# Five years of national airborne pollen monitoring in South Africa: Biome-specific calendars to inform allergy diagnosis and prevention

**DOI:** 10.64898/2025.12.09.693198

**Authors:** Takudzwa Matuvhunye, Dilys M. Berman, Nanike Esterhuizen, Andriantsilavo H. I. Razafimanantsoa, Frank H. Neumann, Dorra Gharbi, Keneilwe Podile, Tshiamo Mmatladi, Boitumelo Langa, Moteng E. Moseri, Linus Ajikah, Angela Effiom, Nikiwe Ndlovu, Lynne J. Quick, Erin Hilmer, Marishka Govender, Shabeer Davids, Andri C. Van Aardt, Linde J.C de Jager, Jubilant V. Sithole, Juanette John, Rebecca M. Garland, Trevor Hill, Jemma Finch, Kama Chetty, Werner Hoek, Marion Bamford, Riaz Y. Seedat, Ahmed I. Manjra, Caryn M Upton, Jonny Peter, the SAPNET consortium

## Abstract

Pollen monitoring is crucial for understanding seasonal patterns, supporting allergy diagnosis and informing early-warning tools to mitigate allergic diseases. The Southern Hemisphere lacks long-term data on pollen seasons, with extremely few from Africa. We present pollen calendars based on five-year data from the South African Pollen Monitoring Network (SAPNET). Airborne pollen from 2019 to 2024 in biomes across South Africa was collected using Hirst-type volumetric spore traps and standard protocols. Daily concentrations were analysed by light microscopy. The five-year mean annual pollen integral (APIn, pollen grains/m³ per year) was calculated for each site. The five-year mean APIn was highest in the Grassland Biome (Bloemfontein, 11654 pg/m³; range 8515 to 14454 pg/m³) and lowest in the Albany Thicket Biome (Gqeberha, 1372; range 853 to 2010 pg/m³). The Grassland Biome (Johannesburg) had the highest averaged tree pollen concentration (7558; range 6575 to 8803 pg/m³). The Savanna Biome (Kimberley) had the highest average grass pollen concentration (3150; 1826 to 3785 pg/m³). The grass family (Poaceae) was the most common pollen type across all biomes. Other common contributing taxa were exotic trees Cupressaceae, *Platanus*, *Morus* and *Betula*. Tree seasons were July to September, whilst grass and weed pollen seasons varied across the different biomes. These five-year pollen calendars provide the first biome-specific national reference for airborne pollen exposure in South Africa. The findings provide baseline data for the clinical management of allergic disease.

## 1 Introduction

Allergic diseases affect approximately 662 million people worldwide, including 400 million with allergic rhinitis (AR) (Lu et al. 2024). The incidence of allergic diseases has increased in recent years possibly due to anthropogenic climate change (Pershad et al. 2025), with the prevalence of AR and asthma among adolescents and university students in South Africa estimated at 40% and 10%, respectively (Zar et al. 2007; Seedat et al. 2018; Mphahlele et al. 2023). Pollen is a dominant aeroallergen driving allergic respiratory diseases. Increased pollen concentrations, changing seasonality and species assemblages have been associated with increased sensitisation, symptomatic AR, and asthma across different populations (Kitinoja et al. 2020; Anuradha et al. 2006; Seedat et al. 2010; Jain et al. 2022). Negative interactions between air pollutants and pollen allergens (Li et al. 2020; Sedghy et al. 2018), together with increasing periods of extreme risk such as thunderstorm asthma events (Davies et al. 2018), highlight the importance of detailed geospatial and temporal information on pollen emissions (Ortega-Rosas et al. 2023).

In Australia, pollen concentrations peak in spring (September to December), with sensitisation of 40-46% being mainly to grasses (Lampugnani et al. 2024; Kam et al. 2016; Davies et al. 2022). In China, spring pollen peaks, primarily from trees, are associated with a lower risk of both allergic rhinitis (AR) and asthma. Autumn pollen peaks are predominantly triggered by weeds (Zhaobin et al. 2024). In Africa, limited records of airborne pollen are available from selected countries, including Nigeria (Adeniyi et al. 2018; Agwu et al. 2005), Algeria (Necib and Boughediri 2016), and Morocco (EIhassani et al. 2022; Raissouni et al. 2024). Pollen from exotic trees such as *Platanus* and *Quercus* is abundant and has been reported as a major allergen among adults in the North West Province of South Africa (Gharbi et al. 2025).

Airborne pollen studies in Africa have been limited to a few cities using different methods (Coetzee and van Zinderen Bakker 1952; Neumann et al. 2025), which has made comparisons difficult. Pollen seasons are influenced by factors such as rainfall, temperature, humidity, wind speed and direction, and vary by geographic location, showing the importance of establishing biome-specific pollen calendars. This underscores the importance of identifying seasonal trends and the specific pollen taxa involved from longer term datasets across a range of biomes.

A pollen calendar is a visual summary of longitudinal trends in timing, duration, and intensity of allergenic pollen over a year, and can support the diagnosis and management of allergic diseases (Lo et al. 2019). Pollen calendars guide allergy sufferers in taking precautions or early preventive medications and assist clinicians in identifying potential triggers and initiating appropriate therapies (Ortega-Rosas et al. 2023). Long-term calendars exist predominantly in the Northern Hemisphere (Cervigón et al. 2024; Camacho et al. 2020). SAPNET, initiated as a national pollen monitoring network in 2019 (Ajikah et al. 2020), has published pollen calendars for seven cities representing the diverse biomes of South Africa, based on two years of airborne monitoring (Esterhuizen et al. 2023). Here, we present the first biome-specific five-year national pollen calendars for South Africa, extending the previous report and demonstrating substantial spatial variability in pollen intensity and seasonality.

## 2 Material and Methods

### 2.1 Sites and airborne pollen sampling

Airborne pollen was collected from August 2019 to August 2024 across seven South African cities representing different biomes, these being, the inland sites: Grassland (Johannesburg), Savanna (Pretoria), Savanna (Kimberley), Grassland (Bloemfontein); and the coastal sites: Fynbos (Cape Town), Indian Ocean Coastal Belt (Durban) and Albany Thicket (Gqeberha, formerly Port Elizabeth) (Fig. 1). A detailed description of the biomes of South Africa is available (Esterhuizen et al. 2023). The standard method for instrument setup, and pollen collection using Hirst-type volumetric spore traps (manufactured by Burkard, UK), and light microscopy were followed, as previously described (Esterhuizen et al. 2023). Detailed descriptions of each biome’s geography and climate can be found in Supplementary Table S1. After seven days of sampling, the Melinex strip was stained and mounted, and analysed using a light microscope at a magnification of X400 (Galán et al. 2014). For each slide representing a 24-h section, three longitudinal transects of the prepared slide were counted (Galán et al. 2014).

**Fig. 1.**
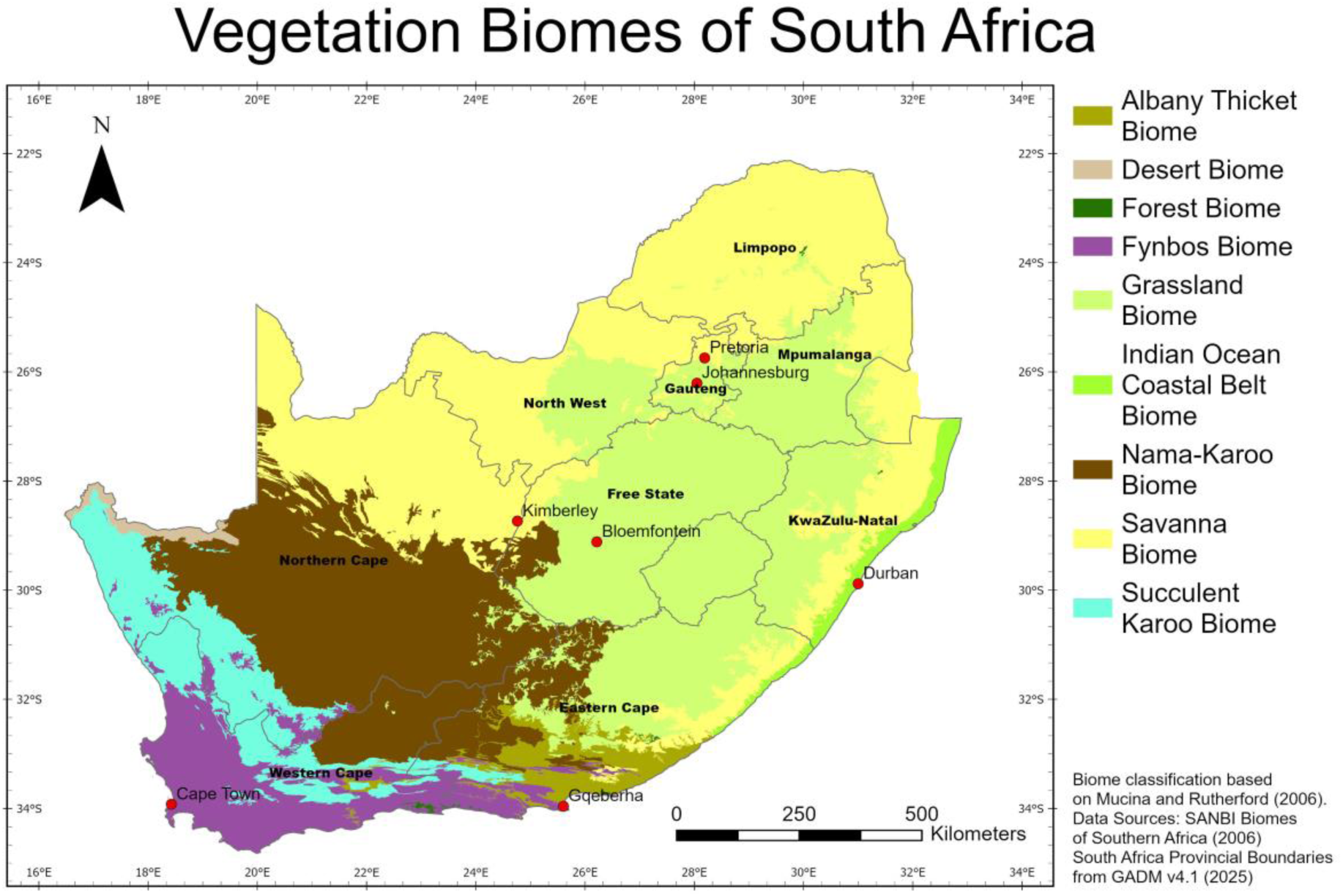
Map of South African biomes (Mucina & Rutherford 2026) showing the seven cities where airborne pollen monitoring was conducted from 2019 to 2024.

Daily pollen concentrations were obtained by converting raw scores into a concentration per cubic meter of air per 24-hour period and applying a correction factor (Esterhuizen et al. 2023). Weekly pollen counts were derived from daily pollen concentrations. Any missing daily data were substituted by the average daily pollen count for that week. The average daily concentrations for the corresponding calendar week in other years were used to fill gaps of greater than six days. This included data for the period 1 July to 11 August 2019, as monitoring of airborne pollen started on 12 August 2019 (Supplementary Fig. S1).

### 2.2 Pollen statistics

The data presented in this paper were calculated by averaging the annual pollen metrics, including the annual pollen integral (APIn), peak count, and peak day. The APIn was calculated as the sum of weekly pollen concentrations for each seasonal year (52 weeks). For all analyses, we defined the seasonal year as 1 July to 30 June of the following year. For the pollen calendars, daily pollen counts were summed to weekly totals and averaged across the 5-year period to generate the average weekly profile over the year. The weekly pollen counts were grouped to determine intensity based on the following scale: 0 (0–3 pg/m³), 1 (3–9 pg/m³), 2 (10–29 pg/m³), 3 (30–99 pg/m³), and 4 (>100 pg/m³) (Esterhuizen et al. 2023). For each city, pollen types contributing at least 3% of the five year mean APIn were included in the calendars. The start and end of the pollen season were defined as the first and last days in the seasonal year when daily grass and weeds (herbaceous shrubs) counts were above 10 pg/m³ and above 15 pg/m³ for trees (Esterhuizen et al. 2023).

## 3 Results

### 3.1 Average Annual Pollen Integral

Overall, the five-year mean APIn was highest in the Grassland Biome (Bloemfontein) (Fig. 2; Table 1). The inland Savanna (Kimberley) had the highest grass pollen, followed by Bloemfontein, with substantially lower counts in the coastal biomes. Johannesburg’s Grassland had the highest APIn for trees, reflecting a strong urban forest component. High tree pollen counts were also observed in Bloemfontein, Pretoria and Cape Town. Weeds were lower than trees and grasses at all sites (Supplementary Fig. S2).

**Fig. 2.**
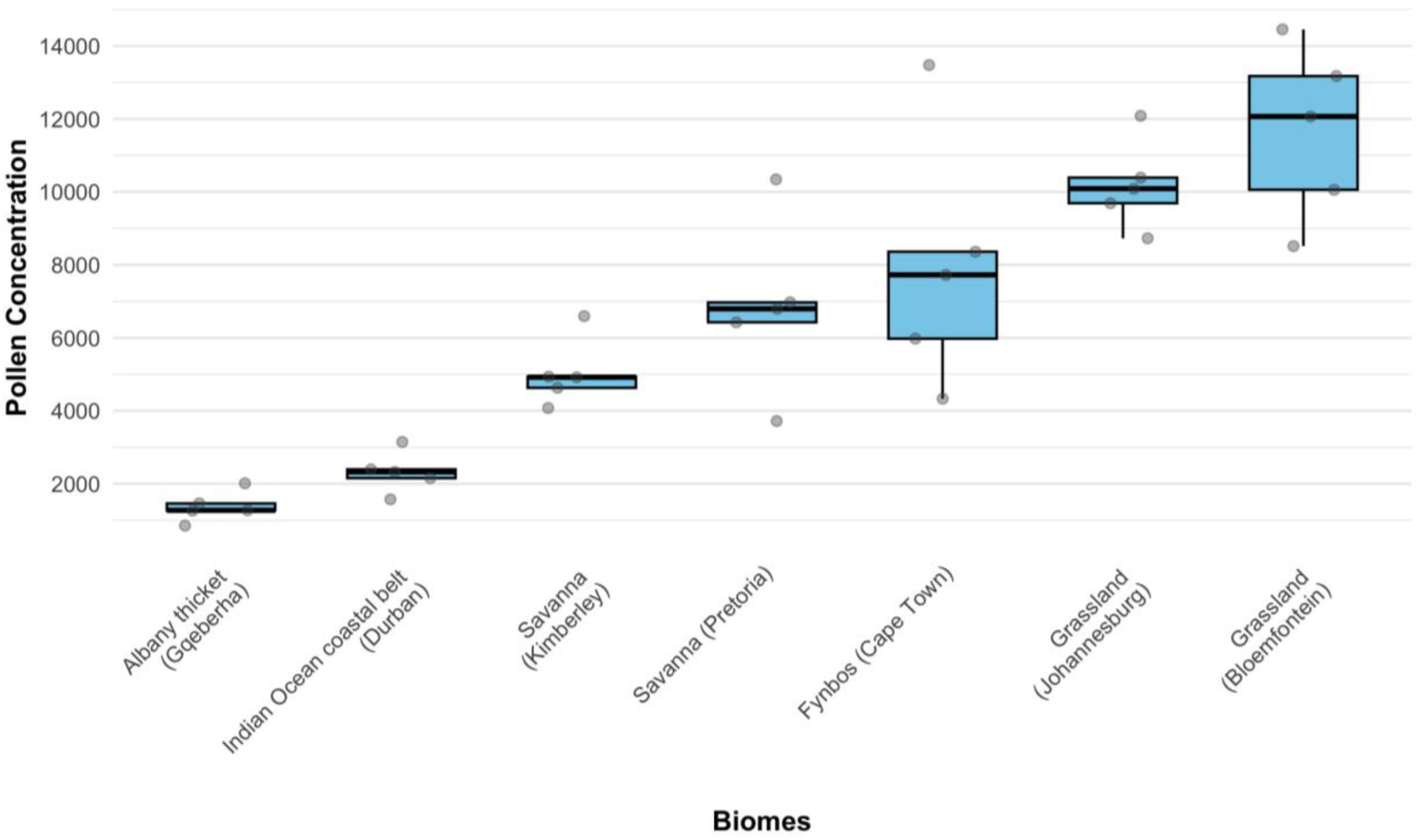
Annual pollen integral for each of the seven South African cities representing distinct biomes over five seasonal years (2019 to 2024). Each dot represents the yearly APIn at each city. The boxplots show the median, interquartile range (IQR), and variability of APIn values within each city.

**Table 1.**
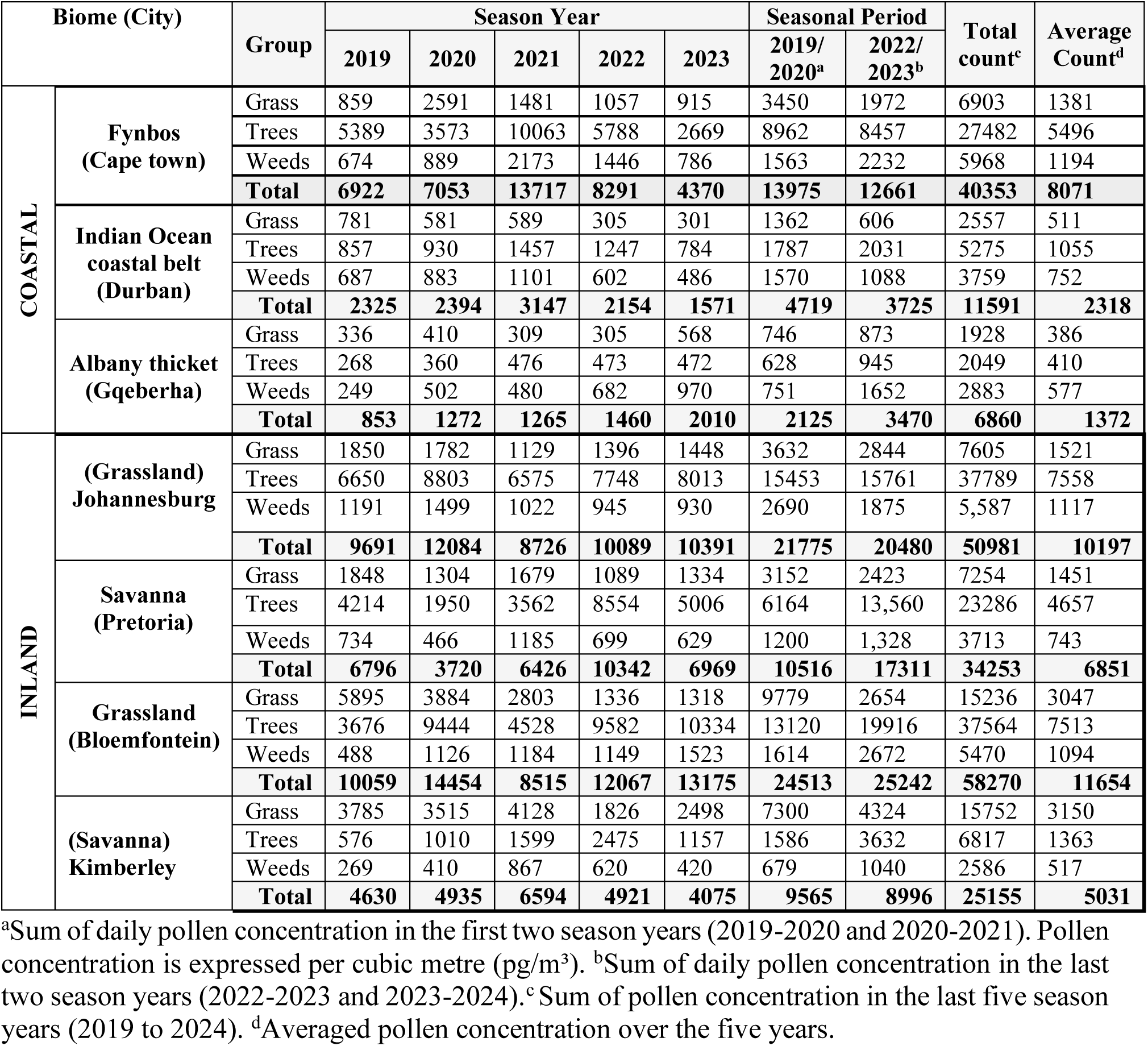
A summary of the annual pollen integral across the South African biomes.

| Biome (City) |  | Group | Season Year |  |  |  |  | Seasonal Period |  | Total count<sup>c</sup> | Average Count<sup>d</sup> |
| --- | --- | --- | --- | --- | --- | --- | --- | --- | --- | --- | --- |
|  |  |  | 2019 | 2020 | 2021 | 2022 | 2023 | 2019/<br>2020<sup>a</sup> | 2022/<br>2023<sup>b</sup> |  |  |
| COASTAL | Fynbos<br>(Cape town) | Grass | 859 | 2591 | 1481 | 1057 | 915 | 3450 | 1972 | 6903 | 1381 |
|  |  | Trees | 5389 | 3573 | 10063 | 5788 | 2669 | 8962 | 8457 | 27482 | 5496 |
|  |  | Weeds | 674 | 889 | 2173 | 1446 | 786 | 1563 | 2232 | 5968 | 1194 |
|  |  | **Total** | **6922** | **7053** | **13717** | **8291** | **4370** | **13975** | **12661** | **40353** | **8071** |
|  | Indian Ocean<br>coastal belt<br>(Durban) | Grass | 781 | 581 | 589 | 305 | 301 | 1362 | 606 | 2557 | 511 |
|  |  | Trees | 857 | 930 | 1457 | 1247 | 784 | 1787 | 2031 | 5275 | 1055 |
|  |  | Weeds | 687 | 883 | 1101 | 602 | 486 | 1570 | 1088 | 3759 | 752 |
|  |  | **Total** | **2325** | **2394** | **3147** | **2154** | **1571** | **4719** | **3725** | **11591** | **2318** |
|  | Albany thicket<br>(Gqeberha) | Grass | 336 | 410 | 309 | 305 | 568 | 746 | 873 | 1928 | 386 |
|  |  | Trees | 268 | 360 | 476 | 473 | 472 | 628 | 945 | 2049 | 410 |
|  |  | Weeds | 249 | 502 | 480 | 682 | 970 | 751 | 1652 | 2883 | 577 |
|  |  | **Total** | **853** | **1272** | **1265** | **1460** | **2010** | **2125** | **3470** | **6860** | **1372** |
| INLAND | (Grassland)<br>Johannesburg | Grass | 1850 | 1782 | 1129 | 1396 | 1448 | 3632 | 2844 | 7605 | 1521 |
|  |  | Trees | 6650 | 8803 | 6575 | 7748 | 8013 | 15453 | 15761 | 37789 | 7558 |
|  |  | Weeds | 1191 | 1499 | 1022 | 945 | 930 | 2690 | 1875 | 5,587 | 1117 |
|  |  | **Total** | **9691** | **12084** | **8726** | **10089** | **10391** | **21775** | **20480** | **50981** | **10197** |
|  | Savanna<br>(Pretoria) | Grass | 1848 | 1304 | 1679 | 1089 | 1334 | 3152 | 2423 | 7254 | 1451 |
|  |  | Trees | 4214 | 1950 | 3562 | 8554 | 5006 | 6164 | 13,560 | 23286 | 4657 |
|  |  | Weeds | 734 | 466 | 1185 | 699 | 629 | 1200 | 1,328 | 3713 | 743 |
|  |  | **Total** | **6796** | **3720** | **6426** | **10342** | **6969** | **10516** | **17311** | **34253** | **6851** |
|  | Grassland<br>(Bloemfontein) | Grass | 5895 | 3884 | 2803 | 1336 | 1318 | 9779 | 2654 | 15236 | 3047 |
|  |  | Trees | 3676 | 9444 | 4528 | 9582 | 10334 | 13120 | 19916 | 37564 | 7513 |
|  |  | Weeds | 488 | 1126 | 1184 | 1149 | 1523 | 1614 | 2672 | 5470 | 1094 |
|  |  | **Total** | **10059** | **14454** | **8515** | **12067** | **13175** | **24513** | **25242** | **58270** | **11654** |
|  | (Savanna)<br>Kimberley | Grass | 3785 | 3515 | 4128 | 1826 | 2498 | 7300 | 4324 | 15752 | 3150 |
|  |  | Trees | 576 | 1010 | 1599 | 2475 | 1157 | 1586 | 3632 | 6817 | 1363 |
|  |  | Weeds | 269 | 410 | 867 | 620 | 420 | 679 | 1040 | 2586 | 517 |
|  |  | **Total** | **4630** | **4935** | **6594** | **4921** | **4075** | **9565** | **8996** | **25155** | **5031** |

<sup>a</sup>Sum of daily pollen concentration in the first two season years (2019-2020 and 2020-2021). Pollen concentration is expressed per cubic metre (pg/m<sup>3</sup>). <sup>b</sup>Sum of daily pollen concentration in the last two season years (2022-2023 and 2023-2024). <sup>c</sup>Sum of pollen concentration in the last five season years (2019 to 2024). <sup>d</sup>Averaged pollen concentration over the five years.

Among the pollen types that contributed at least 3% to the city specific five-year mean, Poaceae was detected in all biomes, whilst *Cupressus* was detected in all biomes, except the Albany thicket. Other common pollen types included *Platanus* (Fynbos, Savanna, Grassland and Grassland), *Morus* (Savanna, Grassland,and the Indian Ocean Coastal Belt) (Fig. 3) and *Betula* (Savanna, Grassland, and the Indian Ocean Coastal Belt).

**Fig. 3.**
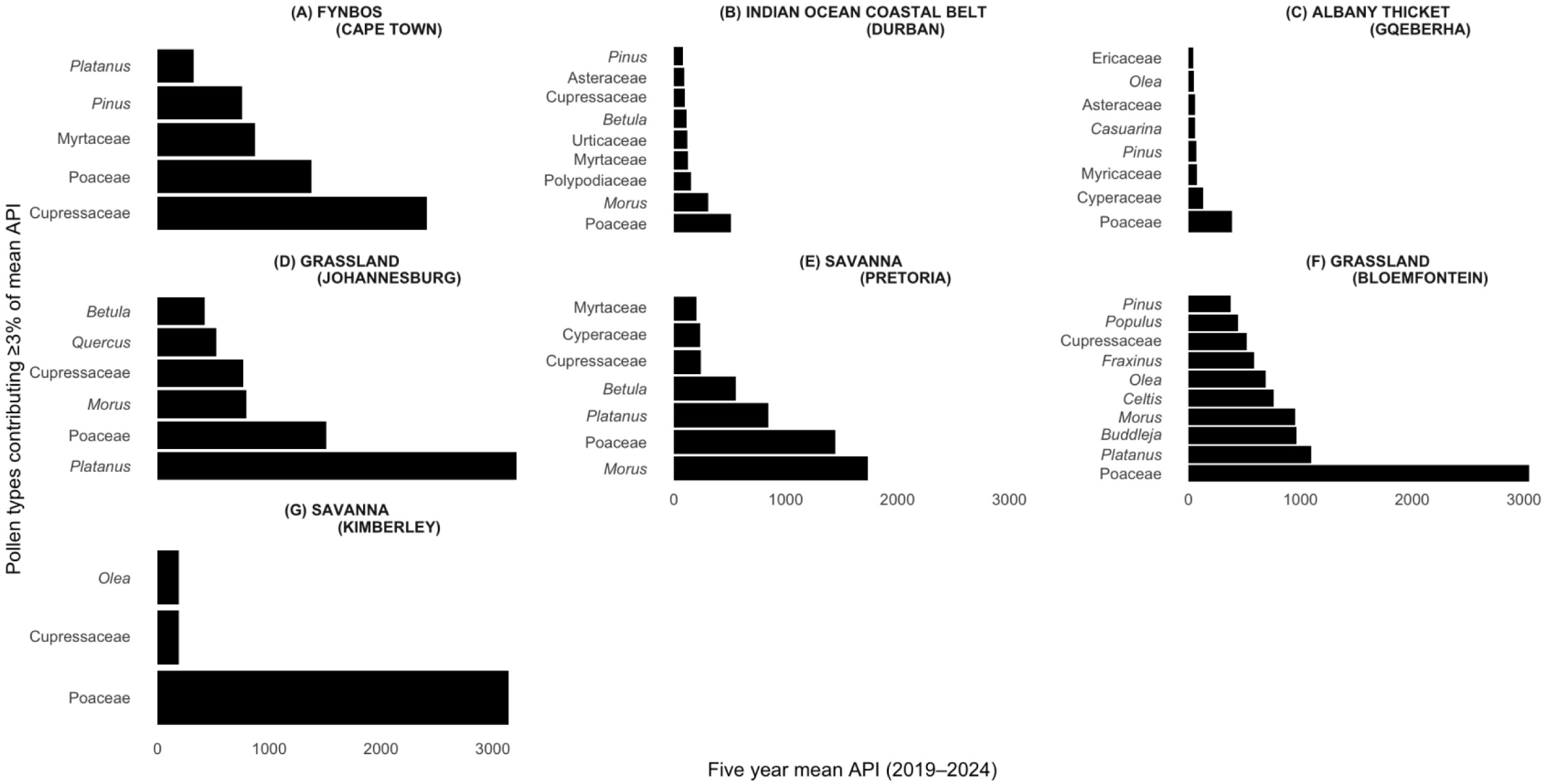
Relative contribution of pollen taxa to the annual pollen integral (APIn) across seven South African cities. Only taxa contributing more than 3% to the five-year mean APIn are shown. Bars represent the cumulative contribution of each taxon from 2019 to 2024, illustrating differences in dominant pollen types across biomes.

### 3.2 Pollen calendars

The start dates (Supplementary Fig. S3) and season durations (Fig. 4) of the pollen seasons for trees, grasses, and weeds varied between taxa and across biomes, with grass seasons being generally longer in inland regions and tree seasons longest in the Fynbos Biome. Across all the biomes, the start month of the tree pollen season was consistently between late July and September. Fynbos (Cape Town) had the longest tree pollen season (mean 297 days), while the Albany Thicket (Gqeberha) had the shortest tree pollen season (mean 116 days) (Table 2). The grass flowering season was longest in the Savanna (Kimberley; mean 247 days), and shortest in the Indian Ocean Coastal Belt Biome (Durban; mean 71 days).

**Fig. 4.**
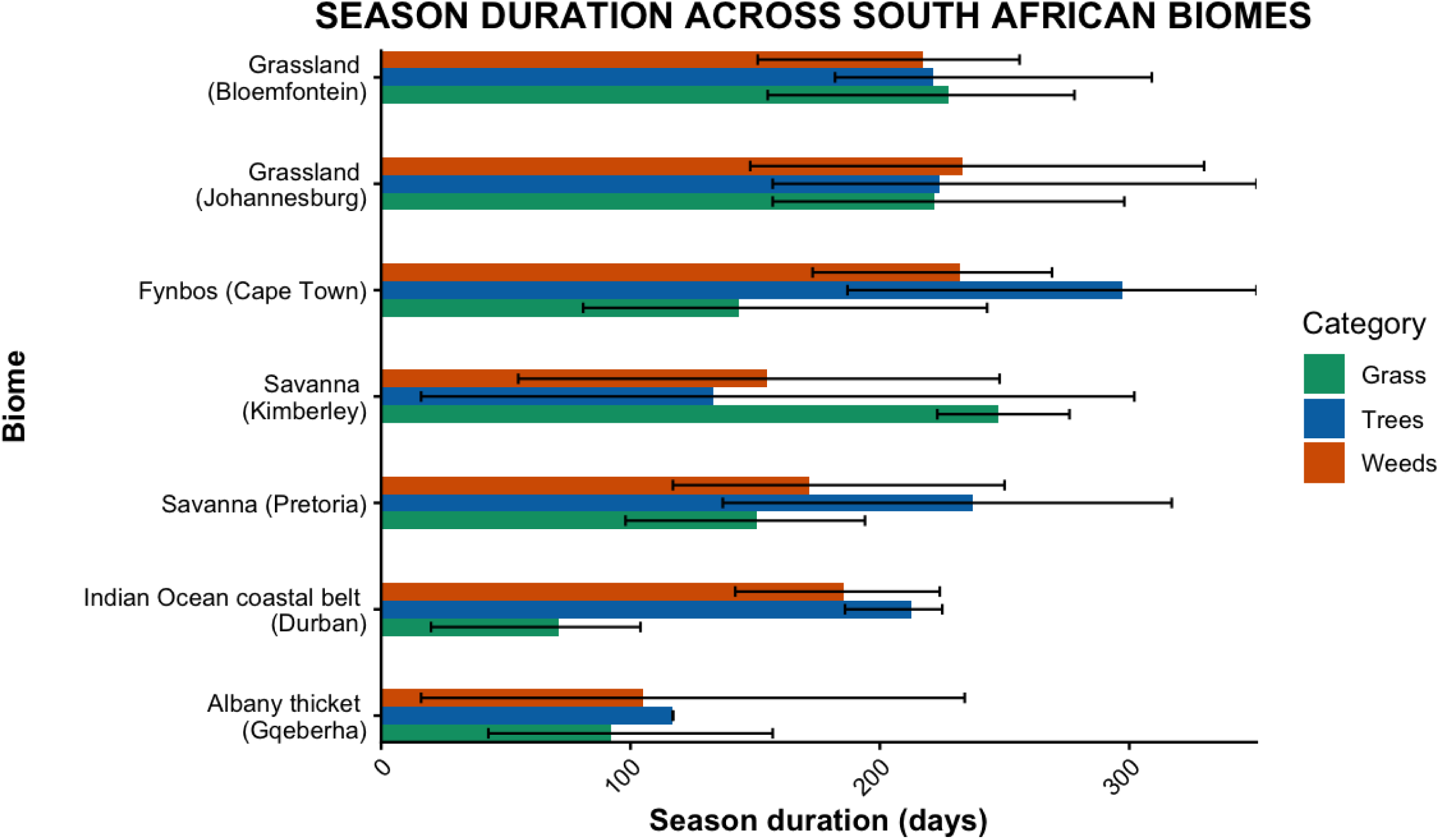
Pollen season duration for grass, tree and weeds across the seven South African cities representing distinct biomes over five seasonal years (2019 to 2024). The bars show the average season duration, while the lines indicate the range.

**Table 2.** Summary of pollen season characteristics from 2019 to 2024 across biomes.

| Biome (City) |  | Pollen Type | Season Start | Season End | Duration in days | Peak Month<br>(Max week and year) |
| --- | --- | --- | --- | --- | --- | --- |
|  |  |  | Mean (range) | Mean (range) | Mean (range) |  |
| COASTAL | Fynbos (Cape Town) | Grass | Sep (6-23 Sep) | Feb (Nov-May) | 144 (81-243) | Oct (26 Oct-1 Nov 2020) |
|  |  | Tree | Jul | May (Jan-Jun) | 297 (187-351) | Sep (13-19 Sep 2021) |
|  |  | Weed | Aug (Jul-Oct) | Apr (Mar-Jun) | 232 (173-269) | Oct (4-10 Oct 2021) |
|  | Indian Ocean Coastal Belt (Durban) | Grass | Feb (Jan-Mar) | Apr (Feb-May) | 71 (20-104) | Feb (17-23 Feb 2020) |
|  |  | Tree | Aug (Jul-Aug) | Mar (Feb-Mar) | 212 (186-225) | Nov (27 Feb-5 Mar 2023) |
|  |  | Weed | Oct (Aug-Jan) | May (Mar-Apr) | 186 (142-224) | Feb (5-11 Apr 2021) |
|  | Albany Thicket (Gqeberha) | Grass | Nov (Aug-Feb) | Feb (Oct-Apr) | 92 (42-157) | Dec (1-7 Apr 2024) |
|  |  | Tree | Sep (Aug-Dec) | April only | 116 | N/A |
|  |  | Weed | Jul/Aug | Dec (Oct-Mar) | 105 (16-234) | Sep (18-24 Sep 2023) |
| INLAND | Grassland (Johannesburg) | Grass | Oct (Jul-Dec) | Jun (May-Jun) | 222 (157-298) | Feb (24 Feb-1 Mar 2020) |
|  |  | Tree | Jul | Feb (Dec-Jun) | 224 (157-279) | Aug (31 Aug-6 Sep 2020) |
|  |  | Weed | Sep (Jul-Dec) | Apr (Jan-Jun) | 233 (148-315) | Dec (5-11 Apr 2021) |
|  | Savanna (Pretoria) | Grass | Nov (Sep-Dec) | Apr (Mar-May) | 150 (98-194) | Jan (24-30 Jan 2022) |
|  |  | Tree | Aug (Jul-Aug) | Mar (Dec-Jun) | 237 (137-317) | Aug (22-28 Aug 2022) |
|  |  | Weed | Oct (Aug-Dec) | Apr/May | 172 (117-250) | Feb (31 Jan-6 Feb 2022) |
|  | Grassland (Bloemfontein) | Grass | Sep | May (Apr-May) | 246 (223-278) | Feb (3-9 Feb 2020) |
|  |  | Tree | Jul-Aug | Mar (Jan-Jun) | 221 (182-309) | Aug (7-13 Sep 2020) |
|  |  | Weed | Sep (Aug-Nov) | Apr (Jan-Jun) | 217 (151-256) | Sep (18-24 Sep 2023) |
|  | Savanna (Kimberley) | Grass | Sep (Aug-Oct) | May | 247 (223-276) | Feb (17-23 Jan 2022) |
|  |  | Tree | Aug (9-25 Aug) | Jan (Oct-Jun) | 163 (69-302) | Sep (6-12 Sep 2021) |
|  |  | Weed | Oct (Aug-Feb) | Mar (Oct-Apr) | 155 (55-248) | Oct (23-29 Aug 2021) |

### 3.3 Biome Characterisation

The biome pollen profiles including start, end, and peak timing for all pollen groups are presented in Table 2.

#### Fynbos Biome (Cape Town)

The Fynbos Biome (Cape Town) showed an APIn that is high among the coastal cities but lower than the highest inland biomes. The biome is characterised by winter rain, high plant biodiversity, and a largely treeless landscape although pines and other exotic trees have been planted (Mucina et al. 2006). The grass season started in September (spring) and lasted 144 days, ending in February (summer) (Fig. 5a and Fig. 6). The Fynbos Biome featured the longest average tree season nationally (297 days), starting consistently in late winter (July). The grass pollen load showed marked inter-annual variability. The pollen profile was dominated by tree taxa (Table 1 and Supplementary Fig. S2). Notably, Cupressaceae accounted for the largest percentage (30%) of the five-year mean APIn. Other significant tree taxa included Myrtaceae (11%), *Pinus* (9%), and *Platanus* (4%). Poaceae contributed substantially (17%). Less common but consistently detected tree pollen types are *Olea* (2.6%) and Urticaceae (2.4%). These low-abundance taxa contribute yearly to the total pollen load.

**Fig. 5.**
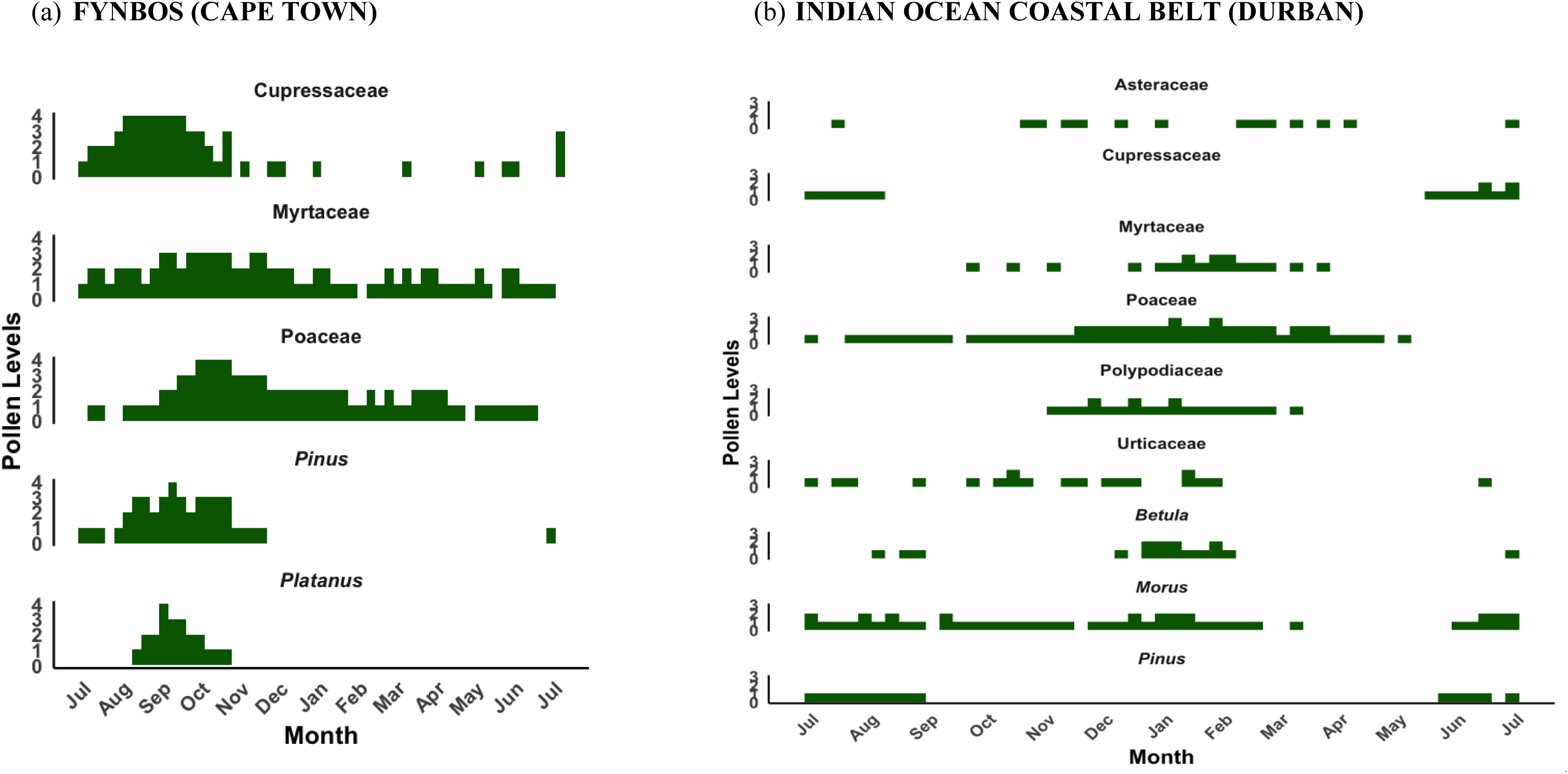
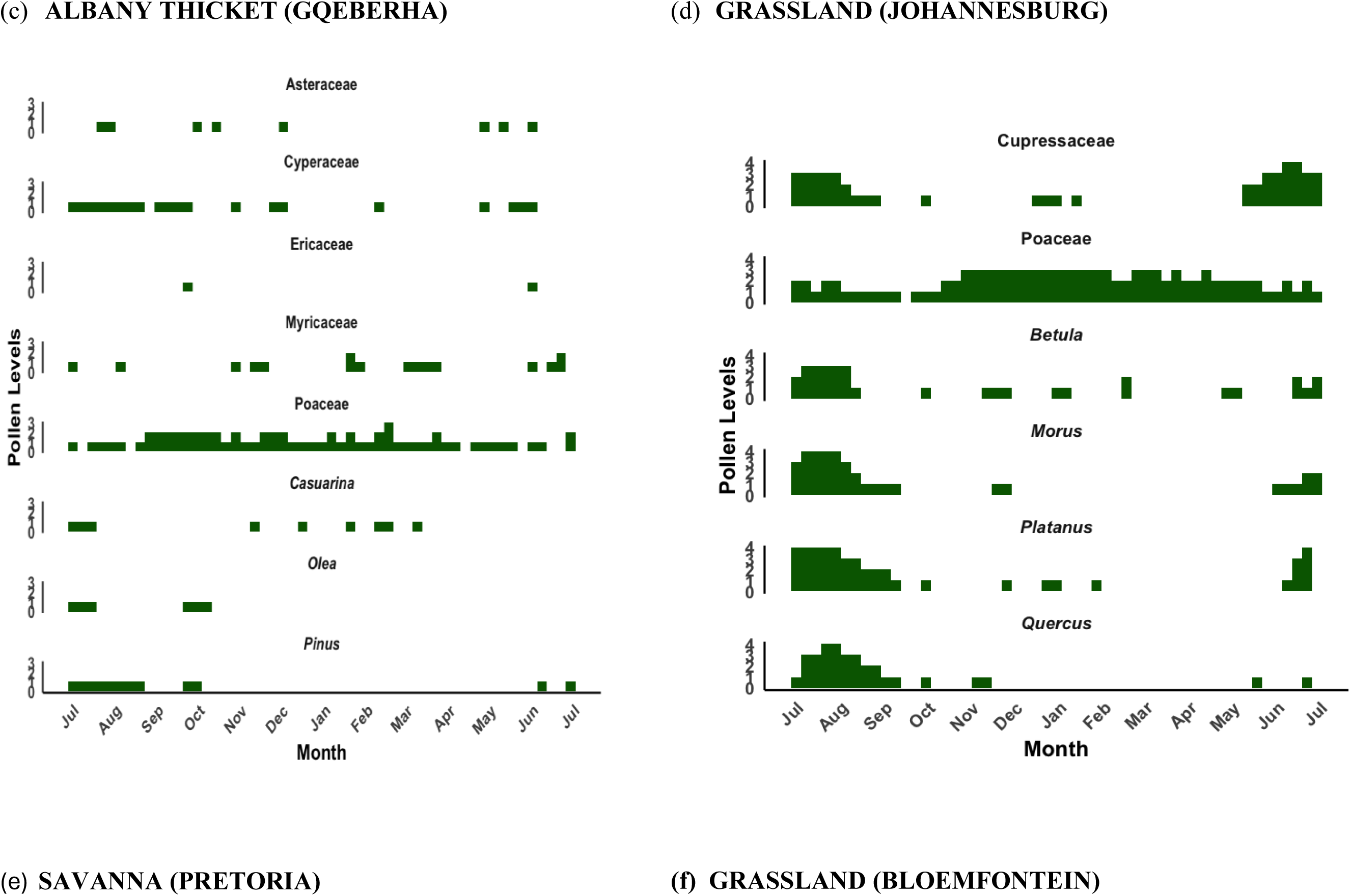
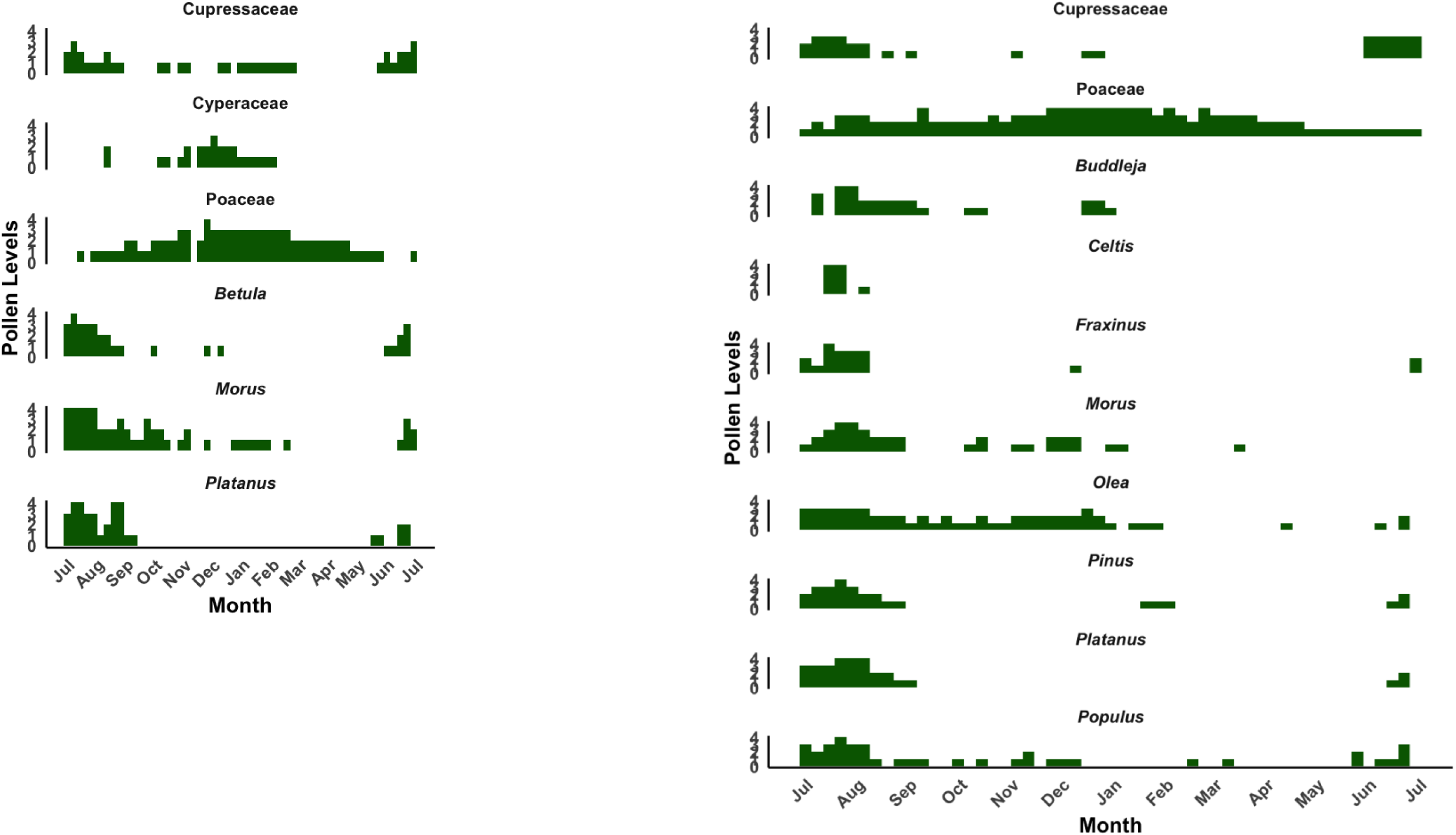
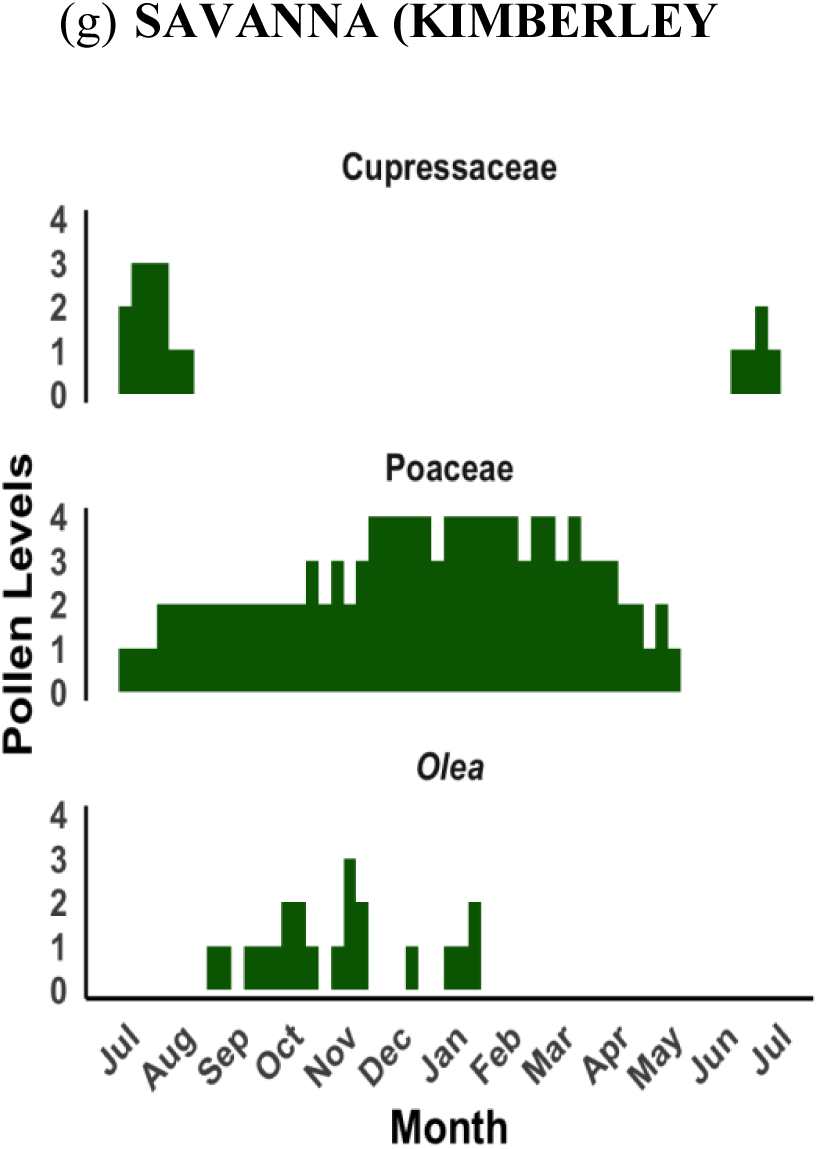
Five-year monthly pollen calendars for seven South African biomes. Stacked blocks represent mean weekly concentrations for each taxon, averaged over five seasonal years (2019 to 2024). Rows represent pollen taxa contributing more than 3% to the annual pollen integral at each biome, shown in A to G. Block height increases with concentration: one block for 0 to 3 pg/m³, two blocks for 3 to 10 pg/m³, three blocks for 10 to 30 pg/m³, and four blocks for concentrations above 30 pg/m³. These calendars highlight differences in timing, duration, and intensity of pollen seasons across biomes.

**Fig. 6.**
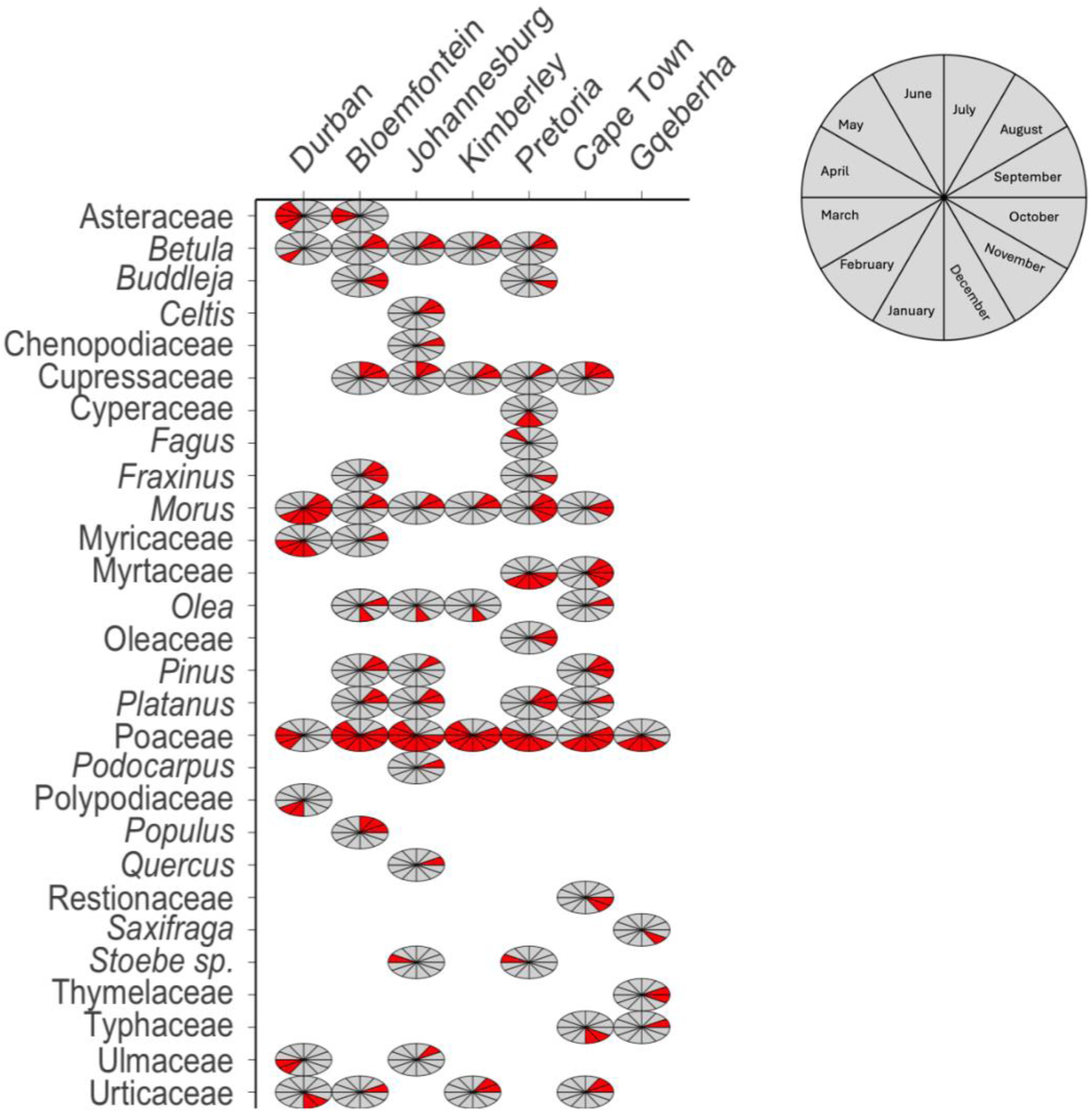
Pollen calendar for taxa with distinct pollen seasons across South African biomes. The pie charts are divided into monthly segments, starting from July, and the red shaded segments depicting the months in which pollen season concencentatration exceeded the seasonal thresholds of 15 pg/m³ for trees, and 10 pg/m³ for grass and weeds.

#### Indian Ocean Coastal Belt Biome (Durban)

The Indian Ocean Coastal Belt Biome (Durban), characterized by (sub) tropical climate with high summer rainfall and abundant tree taxa including palms and mangroves (Mucina et al. 2006), had a low overall pollen catch, consistent with the pattern observed in other coastal biomes (Fig. 2, Table 1). The total pollen catch was evenly distributed across the major groups. Tree pollen was slightly higher than grass and weeds (Table 1). The grass season was the shortest of all the cities measured, lasting an average of 71 days, and starting late in summer. The tree season was long, averaging 212 days, starting in winter. The weed season averaged 186 days, beginning in spring (Fig. 5b). Poaceae contributed 21.9% to the mean APIn, while the exotic tree *Morus*, contributed 13.2%. Other significant trees included Myrtaceae (5.4%), and *Pinus* (3.4%), Polypodiaceae (ferns; 6.6%) were highest within the weed category.

#### The Albany Thicket Biome (Gqeberha, formerly Port Elizabeth)

Albany Thicket Biome (Gqeberha), features dense evergreen, sclerophyllous or succulent shrubland with high biodiversity under predominantly all-year rainfall conditions (Mucina et al. 2006). The lowest five-year mean APIn was observed among all seven cities and this biome featured the shortest pollen seasons nationally for trees and grasses. The tree and grass pollen loads were lower than in the other biomes, and their seasons were shorter, lasting an average of 116 days and 92 days, respectively. Weed seasons were short, lasting an average of 105 days, and showed low to moderate levels of pollen throughout, resulting in no clear seasons (Fig. 5c). Poaceae was the leading contributor, accounting for 28% of the mean APIn. Other prominent taxa contributing more than 3% of the APIn yearly included Cyperaceae (9%), Myricaceae (5%), and *Pinus* (5%).

#### Grassland Biome (Johannesburg)

The Grassland had the highest averaged tree pollen concentration among all monitored sites, mainly contributed by exotic trees. The biome is situated in the summer-rainfall region, largely tree-less and often highly transformed due to agriculture and urbanization, including the introduction of abundant exotic trees (Mucina et al. 2006). The high five-year mean APIn included a high contribution of tree and grass pollen (Table 1). The pollen season was characterised by a similarly long period of high tree concentration that lasted an average of 224 days, starting in mid-winter (Table 2). The grass season was similarly long, lasting an average of 222 days. Weed seasons had the longest average duration nationally at 233 days, beginning in early spring. The pollen profile was highly diverse and dominated by several key tree taxa (Fig. 5d). The leading contributor was the alien tree *Platanus*, accounting for the largest percentage of the mean APIn (31%). Other highly relevant tree taxa included *Morus* (7.8%), Cupressaceae (7.6%), *Quercus* (5.2%), and *Betula* (4.1%). Poaceae also contributed substantially to the overall load (14.8%). Other pollen types consistently detected every year included Asteraceae, *Fraxinus*, and *Celtis* (Supplementary Table 2).

#### Savanna Biome (Pretoria)

Savanna (Pretoria) pollen spectra recorded a high overall APIn (Fig. 2, Table 1). The Savanna Biome is a grass-dominated ecosystem with subtropical trees such as acacias (Mucina et al. 2006). The largest contributor to the APIn was tree pollen, with grass and weed pollen contributing a smaller but substantial portion (Table 1, Supplementary Fig. 2). This biome featured long pollen seasons, where the tree season averaged 237 days, starting early in spring. The grass season was shorter, averaging 150 days, while the weed season averaged 172 days, both starting in spring. Moraceae (25.3%) dominated the tree pollen catch. Poaceae was the second highest contributor (21%). Other significant tree taxa included *Platanus* (12%), *Betula* (8%), Cupressaceae (3.5%), and Myrtaceae (3%) (Supplementary Table S3).

#### Grassland (Bloemfontein)

The Grassland Biome, where Bloemfontein is situated, recorded the highest five-year mean APIn in the study, due to a combination of high tree and grass pollen concentrations. The overall pollen load was dominated by the combined contribution of tree and grass pollen (Table 1, Supplementary Fig. S2). Long pollen seasons were consistently observed for all major groups. The grass flowering season was the longest, averaging 246 days and starting in early spring (Supplementary Fig. S4 and Fig. S5). The tree season was also long, averaging 221 days and beginning in winter. The weed season averaged 217 days, starting in spring. Unlike most other biomes, Poaceae was the only pollen type that consistently contributed more than 3% of the APIn throughout the five years, accounting for 26% of the mean APIn. Other pollen types that were common every year, though contributing slightly below the 3% threshold, included Cyperaceae (2.7%) and Asteraceae (2.4%).

#### Savanna Biome (Kimberley)

Savanna (Kimberley), with low rainfall and an open, grassy landscape (Mucina et al. 2006), recorded the highest averaged grass pollen concentration among all seven monitored sites. The overall five-year mean APIn was characterized by a distinct grass dominance, with grass pollen accounting for the largest contribution to the total load (Table 1). The grass season was notably long, averaging 247 days, and starting relatively early in spring. The tree season was shorter, lasting an average of 163 days, and starting in winter. The weed season was intermediate, averaging 155 days. The pollen profile was dominated by grasses (Poaceae). In terms of tree taxa, Oleaceae and Cupressaceae were the only types that contributed more than 3% of the APIn yearly. Other consistent contributors that were above 2% included Moraceae (2.9%) and *Betul*a (2.7%).

### 3.4 Diversity of pollen taxa

A total of 103 unique pollen taxa were identified across the biomes of South Africa, including 51 trees and 52 weed pollen types (Supplementary Table S4). Weeds were the most diverse pollen types in the Indian Ocean Coastal Belt (Durban) and Albany Thicket Biome (Gqeberha), while trees were more diverse than weeds in the Grassland (Bloemfontein) and the Savanna biomes (Pretoria) (Supplementary Fig. S6). Tree pollen showed greater diversity in September and October in all biomes, except in the Grassland (Johannesburg) where it peaked in August. The grasses included *Zea mays* and species of the Poaceae family. The pollen species of Poaceae that are common in South African biomes, based on data from the Global Biodiversity Information Facility (GBIF), are listed in the supplementary data (35) (Supplementary Fig. 7).

### 3.5 Pollen concentrations in 2019-2021 and 2022-2024

A comparison of overall pollen concentrations between the first two seasonal years (2019-2020 and 2020-2021) and the last two years (2022-2023 and 2023-2024) showed a drop in pollen concentrations over time across most biomes, except in Pretoria (10516 vs 17311 pg/m³), Bloemfontein (24513 vs 25242 pg/m³) and Gqeberha (2125 vs 3470 pg/m³) (Table 1). A similar pattern was observed for grass pollen, with concentrations almost halving over time across all biomes, except in Gqeberha where there was a slight increase (746 vs 873 pg/m³). In contrast, tree pollen showed a different pattern in several biomes, with concentrations doubling during the last two years in Pretoria (6164 vs 13560 pg/m³), Bloemfontein (13120 vs 19916 pg/m³), and Kimberley (1586 vs 3632 pg/m³). There was no clear pattern for weed pollen, although three biomes showed increases: Cape Town (1563 vs 2232 pg/m³), Bloemfontein (1614 vs 2672 pg/m³) and Kimberley (679 vs 1040 pg/m³). Analyses of years with high and low pollen concentrations revealed that seasonal patterns were similar and differences were mainly in the concentration of pollen (Supplementary Fig. S8). The taxa contributing ≥3% of the APIn were consistent across the years (Supplementary Table S2).

## 4 Discussion

We have provided national biome-resolved pollen calendars for South Africa, based on five years of standardised airborne pollen monitoring across seven cities. These calendars extend the earlier two-year report from SAPNET (Esterhuizen et al. 2023) and establish the first long-term evidence base for understanding airborne allergen exposure nationally. Our results offer a practical reference to support area-specific clinical practice, anticipate exacerbation risks, plan public health measures, and to develop early-warning systems for allergic diseases.

Inland biomes, particularly the Grassland (Bloemfontein) and Savanna (Kimberley), had markedly higher annual pollen loads and longer seasons relative to coastal biomes. Potential reasons are complex and might encompass higher rainfall rates at the coast which wash out pollen from the atmosphere, stronger temperature fluctuations including urban heat island effects in more continental, interior cities (Schramm et al. 2021). Coastal biomes such as Cape Town, Durban, and Gqeberha had shorter, concentrated seasons reflecting differences in vegetation structure, land use, humidity, and wind patterns, as previously reported (Esterhuizen et al. 2023; González Minero et al. 1993). Inland grass seasons were greater than 200 days, whereas Durban and Gqeberha had short seasons of less than 100 days. Similar patterns have been reported in Australia and southern Europe, where Mediterranean and subtropical climates are associated with shorter, more intense pollen peaks compared with temperate continental climates which sustain extended flowering, and longer seasons (Frenguelli et al. 2014; Davies et al. 2022; Montiel et al. 2025).

Tree pollen seasons were consistent across all biomes, starting in late winter, a pattern also reported in Europe and North America (Paschalidou et al. 2020; Monroy-Colín et al. 2020). Cape Town had an exceptionally long tree season lasting nearly 300 days, which may blur the distinction between seasonal and perennial allergens. The need for biome-specific pollen calendars rather than referencing calendars from Europe or North America is a result of complex climate zone in South Africa. Here we have shown that risk profiles differ substantially by biome and require region specific interpretation. The dominant allergenic taxa varied by biome. Grasses (Poaceae) were detected in all cities and formed the primary aeroallergen nationally, consistent with global patterns (Medek et al. 2016; Camacho et al. 2020; Ortega-Rosas et al. 2023). Poaceae includes over 12 000 wind-pollinated species and is a major allergen source in Europe (García-Mozo 2017; Frisk et al. 2023).

Tree pollen contributed disproportionately to the pollen spectra of Johannesburg and Pretoria, where the urban forest is dominated by allergenic alien tree species such as *Platanus*, Cupressaceae, *Quercus*, *Betula, Pinus,* and Myrtaceae (mostly pollen of Australian exotic *Eucalyptus*) (Caillaud et al. 2014; Huang et al. 2024). Birch (*Betula*), although alien to South Africa, was detected in all biomes and is a major cause of allergic rhinitis in the Northern Hemisphere (Biedermann et al. 2019; D’Amato et al. 1998), highlighting exposure not reflected in typical local allergen panels (Murray et al. 2022; Pedretti et al. 2025). Weed pollen was the least abundant group but showed taxonomic diversity in coastal areas, where fynbos like erica and protea species were regularly identified from sampling. We observed a decline in pollen concentrations between the first two years and the last two years. This anomaly requires further investigation together with metereological parameters and landuse changes that may have affected pollen concentrations.

Strengths of this study include the use of standardised Hirst-based methods across multiple biomes, enabling quantitative comparison between cities, and the generation of clinically accessible weekly pollen calendars. However, a limitation is coverage, with pollen monitoring restricted to only seven urban centres, leaving many provinces and rural areas without direct coverage, although other cities have been mapped as reported previously (Gharbi et al. 2024; Cadman et al. 1994). The calendars therefore provide a national reference framework of substantial data but may not capture local exposure in underrepresented provinces. Another limitation is the inability of light microscopy to determine Poaceae beyond the family level, despite its importance as the dominant allergenic group nationally.

These findings have significant implications for healthcare and public health. Pollen calendars can guide diagnostic testing, inform treatment plans, and advise patients on when to initiate or escalate preventive therapy. For public health agencies, the data provide a foundation for development of an early warning system that integrates pollen, air quality, and weather information to anticipate surges in allergic rhinitis and asthma. Maintaining and expanding pollen monitoring in South Africa is therefore a core component of environmental health surveillance. Without continuous data, changing exposure cannot be tracked, and early warnings cannot be implemented or scaled. Strengthening the network through expanded geographic coverage, incorporation of molecular identification (such as DNA metabarcoding for Poaceae (Van Haeften et al. 2024) and fungal spores (Banchi et al. 2020), and integration into national health information systems will be critical for supporting allergy care, reducing respiratory morbidity and mortality, and preparing for the impacts of climate variability. The calendars presented here provide both a baseline and a compelling rationale for sustaining and enhancing pollen monitoring as part of South Africa’s broader climate and health response.

## 5 Conclusion

In this study, we have provided the first five-year, biome-specific pollen calendars for South Africa. Our biomes show substantial variation in pollen exposure between regions, taxa, and years. Poaceae and alien tree taxa such as *Platanus*, Cupressaceae, *Betula* and Myrtaceae are the dominant aeroallergens. These findings establish a national reference for pollen exposure patterns and underscore the need for ongoing monitoring. Expansion to map unexplored areas and to understand local variability and the impact of alien taxa is necessary.

## Supporting information

https://uctcloud-my.sharepoint.com/:w:/g/personal/mtvtak006_myuct_ac_za/IQB4iXi5mvNhQ5Fq_o_7XqmkAYtUsPkPsbpbD2lu8ZploJk

## Data availability

Additional results are in the Supplementary, then all datasets used or analysed during the current study are available from the corresponding author upon reasonable request.

## Acknowledgements

The South African Pollen Monitoring Network acknowledges the following parties for their logistical support, pollen trap management, and pollen data collection towards this manuscript: Wisahl Wallace, Noejfah Jardien, Sarah Pedretti (UCT Lung Institute, Cape Town); Kevwe Eweto, Prosper Bande (University of the Witwatersrand, Johannesburg); Kristen Nienaber, Lize Joubert, Mawethu Ndiki, Marius Müller (University of the Free State, Bloemfontein.

## Contributions

**Takudzwa Matuvhunye** led the data analysis and writing, and contributed to data interpretation and curation. **Dilys M. Berman** led the study conceptualization and project administration, and contributed to investigation, data curation, interpretation, review, and editing. **Nanike Esterhuizen** and **Dorra Gharbi** led data curation and contributed to project administration, review, and editing. **Andriantsilavo H. I. Razafimanantsoa**, **Keneilwe Podile**, **Tshiamo Mmatladi**, **Boitumelo Langa**, **Moteng E. Moseri**, **Linus Ajikah**, **Angela Effiom**, **Nikiwe Ndlovu**, **Erin Hilmer**, **Marishka Govender**, **Shabeer Davids**, J**.C. Linde de Jager**, and **Jubilant V. Sithole** contributed to the investigation, review, and editing. **Lynne J. Quick**, **Frank H**. **Neumann**, **Andri C. van Aardt**, **Juanette John**, **Rebecca M. Garland**, **Trevor Hill**, **Jemma Finch**, **Kama Chetty**, **Werner Hoek**, **Marion Bamford, Riaz Y. Seedat, Ahmed I. Manjra**, and **Caryn M. Upton** contributed to study conceptualization, supervison, review, and editing. Jonny Peter led the funding acquisition and supervision, and contributed to data analysis, writing, interpretation, review, and editing.

## Supplemenray material

Supplementary file 1

## References

Adeniyi TA, Adeonipekun PA, Olowokudejo JD (2018) Annual Records of Airborne Pollen of Poaceae in Five Areas in Lagos, Nigeria. Grana 57 (4):284–291. doi:10.1080/00173134.2017.1356865

Agwu COC, Njokuocha RC, Mezue O (2005) The Study Of Airborne Pollen And Spores Circulating At “Head Level” In Nsukka Environment. Bio-Research 2 (2):7–14. doi:10.4314/br.v2i2.28552

Ajikah L, Neumann F, Berman D, Peter J (2020) Aerobiology in South Africa: A new hope! South African Journal of Science 116. doi:10.17159/sajs.2020/8112

Anuradha B, Vijayalakshmi V, Latha GS, Priya VHS, Murthy K (2006) Profile of pollen allergies in patients with asthma, allergic rhinitis and urticaria in Hyderabad. INDIAN JOURNAL OF CHEST DISEASES AND ALLIED SCIENCES 48 (3):221

Banchi E, Ametrano CG, Tordoni E, Stanković D, Ongaro S, Tretiach M, Pallavicini A, Muggia L, Verardo P, Tassan F (2020) Environmental DNA assessment of airborne plant and fungal seasonal diversity. Science of the Total Environment 738:140249

Biedermann T, Winther L, Till SJ, Panzner P, Knulst A, Valovirta E (2019) Birch pollen allergy in Europe. Allergy 74 (7):1237–1248. 10.1111/all.13758

Cadman A, Dames J, Treblanche A (1994) Airspora concentrations in the Vaal Triangle: monitoring and potential health effects. 1. Pollen.

Caillaud D, Martin S, Segala C, Besancenot JP, Clot B, Thibaudon M (2014) Effects of airborne birch pollen levels on clinical symptoms of seasonal allergic rhinoconjunctivitis. Int Arch Allergy Immunol 163 (1):43–50. doi:10.1159/000355630

Camacho I, Caeiro E, Nunes C, Morais-Almeida M (2020) Airborne pollen calendar of Portugal: a 15-year survey (2002–2017). Allergologia et Immunopathologia. doi:10.1016/j.aller.2019.06.012

Cervigón P, Ferencova Z, Cascón Á, Romero-Morte J, Galán Díaz J, Sabariego S, Torres M, Montserrat Gutiérrez-Bustillo A, Rojo J (2024) Progressive pollen calendar to detect long-term changes in the biological air quality of cities in the Madrid Region, Spain. Landscape and Urban Planning 247:105053. 10.1016/j.landurbplan.2024.105053

Coetzee J, van Zinderen Bakker E (1952) The pollen spectrum of the southern Middleveld of the Orange Free State. South African Journal of Science 48 (9):275

D’Amato G, Spieksma FTM, Liccardi G, Jäger S, Russo M, Kontou-Fili K, Nikkels H, Wüthrich B, Bonini S (1998) Pollen-related allergy in Europe. Allergy 53 (6):567–578

Davies J, Thien F, Hew M (2018) Thunderstorm asthma: controlling (deadly) grass pollen allergy. Bmj 360:Article number: k432

Davies JM, Smith BA, Milic A, Campbell B, Van Haeften S, Burton P, Keaney B, Lampugnani ER, Vicendese D, Medek D, Huete A, Erbas B, Newbigin E, Katelaris CH, Haberle SG, Beggs PJ (2022) The AusPollen partnership project: Allergenic airborne grass pollen seasonality and magnitude across temperate and subtropical eastern Australia, 2016-2020. Environ Res 214 (Pt 1):113762. doi:10.1016/j.envres.2022.113762

EIhassani L, Boullayali A, Janati A, Achmakh L, Bouziane H (2022) Aerobiological study of airborne pollen in Tétouan (NW of Morocco): diversity, intensity and calendar. Aerobiologia 38 (4):483–499

Esterhuizen N, Berman DM, Neumann FH, Ajikah L, Quick LJ, Hilmer E, Van Aardt A, John J, Garland R, Hill T (2023) The South African pollen monitoring network: Insights from 2 years of national aerospora sampling (2019–2021). Clinical and Translational Allergy 13 (11):e12304

Frenguelli G, Ghitarrini S, Tedeschini E (2014) Climatic change in Mediterranean area and pollen monitoring. Flora Mediterranea 24:99–107

Frisk CA, Adams-Groom B, Smith M (2023) Isolating the species element in grass pollen allergy: A review. Sci Total Environ 883:163661. doi:10.1016/j.scitotenv.2023.163661

Galán C, Smith M, Thibaudon M, Frenguelli G, Oteros J, Gehrig R, Berger U, Clot B, Brandao R, group EQw (2014) Pollen monitoring: minimum requirements and reproducibility of analysis. Aerobiologia 30 (4):385–395

García-Mozo H (2017) Poaceae pollen as the leading aeroallergen worldwide: A review. Allergy 72 (12):1849–1858. 10.1111/all.13210

Gharbi D, Berman D, Neumann FH, Hill T, Sidla S, Cillers SS, Staats J, Esterhuizen N, Ajikah L, Moseri ME, J. Quick L, Hilmer E, Van Aardt A, John J, Garland R, Finch J, Hoek W, Bamford M, Seedat RY, I. Manjra A, Peter J (2024) Ambrosia (ragweed) pollen — A growing aeroallergen of concern in South Africa. World Allergy Organization Journal 17 (12):101011. 10.1016/j.waojou.2024.101011

Gharbi D, Neumann FH, Podile K, McDonald M, Linde J-h, Frampton M, Liebenberg JL, Cilliers S, Mmatladi T, Nkosi P, Paledi K, Piketh S, Staats J, Burger RP, Havenga H, Garland RM, Bester P, Lebre PH, Ricci C (2025) Exposure to outdoor aerospora and associated respiratory health risks among adults in Potchefstroom, North-West province, South Africa. Frontiers in Allergy Volume 6 - 2025. doi:10.3389/falgy.2025.1568669

González Minero FJ, Herrero Villacorta B, Candau P (1993) Latitudinal study of allergenic pollen in two Spanish cities. J Investig Allergol Clin Immunol 3 (6):304–310

Huang Z, Li A, Zhu H, Pan J, Xiao J, Wu J, Han Y, Zhong L, Sun X, Wang L, Hu L, Wang C, Ma X, Qiao Z, Zhang M, Yuan L, Liu X, Tang J, Li Y, Yu H, Zheng Z, Sun B (2024) Multicenter study of seasonal and regional airborne allergens in Chinese preschoolers with allergic rhinitis. Scientific Reports 14 (1):4754. doi:10.1038/s41598-024-54574-z

Jain S, Jain A, Gupta SK (2022) Study of Allergen Patterns in Cases of Moderate to Severe Persistent Allergic Rhinitis in Central India. Indian J Otolaryngol Head Neck Surg 74 (Suppl 2):888–893. doi:10.1007/s12070-020-01954-2

Kam AW, Tong WW, Christensen JM, Katelaris CH, Rimmer J, Harvey RJ (2016) Microgeographic factors and patterns of aeroallergen sensitisation. Medical Journal of Australia 205 (7):310–315

Kitinoja MA, Hugg TT, Siddika N, Rodriguez Yanez D, Jaakkola MS, Jaakkola JJK (2020) Short-term exposure to pollen and the risk of allergic and asthmatic manifestations: a systematic review and meta-analysis. BMJ Open 10 (1):e029069. doi:10.1136/bmjopen-2019-029069

Lampugnani ER, Silver JD, Burton P, Nattala U, Katelaris CH (2024) Seasonal Patterns and Allergenicity of Casuarina Pollen in Sydney, Australia: Insights from 10 Years of Monitoring and Skin Testing. Atmosphere 15 (6):719

Li CH, Sayeau K, Ellis AK (2020) Air Pollution and Allergic Rhinitis: Role in Symptom Exacerbation and Strategies for Management. Journal of Asthma and Allergy 13 (null):285–292. doi:10.2147/JAA.S237758

Lo F, Bitz CM, Battisti DS, Hess JJ (2019) Pollen calendars and maps of allergenic pollen in North America. Aerobiologia (Bologna) 35 (4):613–633. doi:10.1007/s10453-019-09601-2

Lu MY, Shobnam N, Livinski AA, Saksena S, Salters D, Biete M, Myles IA (2024) Examining allergy related diseases in Africa: A scoping review protocol. PLOS ONE 19 (2):e0297949. doi:10.1371/journal.pone.0297949

Medek DE, Beggs PJ, Erbas B, Jaggard AK, Campbell BC, Vicendese D, Johnston FH, Godwin I, Huete AR, Green BJ, Burton PK, Bowman DMJS, Newnham RM, Katelaris CH, Haberle SG, Newbigin E, Davies JM (2016) Regional and seasonal variation in airborne grass pollen levels between cities of Australia and New Zealand. Aerobiologia 32 (2):289–302. doi:10.1007/s10453-015-9399-x

Monroy-Colín A, Maya-Manzano JM, Silva-Palacios I, Tormo-Molina R, Pecero-Casimiro R, Gonzalo-Garijo Á, Fernández-Rodríguez S (2020) Phenology of Cupressaceae urban infrastructure related to its pollen content and meteorological variables. Aerobiologia 36 (3):459–479. doi:10.1007/s10453-020-09645-9

Montiel N, Hidalgo PJ, Adame JA, González-Minero F (2025) Pollen season variations among anemophilous species in an Atlantic-influenced mediterranean environment: a long term study (1993–2022). International Journal of Biometeorology 69 (1):109–122. doi:10.1007/s00484-024-02796-1

Mphahlele R, Lesosky M, Masekela R (2023) Prevalence, severity and risk factors for asthma in school-going adolescents in KwaZulu Natal, South Africa. BMJ Open Respir Res 10 (1). doi:10.1136/bmjresp-2022-001498

Mucina L, Rutherford MC, Powrie LW (2006) Vegetation Atlas of South Africa, Lesotho and Swaziland. The vegetation of South Africa, Lesotho and Swaziland:748–789

Murray L, Rooyen CV, Berg SVd, Green RJ (2022) Allergic sensitisation in South Africa: allergen-specific ige-component testing (ISAC). Current Allergy & Clinical Immunology 35 (1):3–8. doi:doi:10.10520/ejc-caci-v35-n1-a4

Necib A, Boughediri L (2016) Airborne pollen in the El-Hadjar town (Algeria NE). Aerobiologia 32 (2):277–288. doi:10.1007/s10453-015-9398-y

Neumann FH, Gharbi D, Ajikah L, Scott L, Cilliers S, Staats J, Berman D, Moseri ME, Podile K, Ndlovu N (2025) Ecological and allergenic significance of atmospheric pollen spectra from a Grassland-Savanna ecotone in North West province, South Africa. Palynology 49 (2):2411234

Ortega-Rosas CI, Gutiérrez-Ruacho OG, Brito-Castillo L, Calderón-Ezquerro MC, Guerrero-Guerra C, Amaya-García V (2023) Five-year airborne pollen calendar for a Sonoran Desert city and the relationships with meteorological variability. Int J Biometeorol 67 (11):1853–1868. doi:10.1007/s00484-023-02546-9

Paschalidou AK, Psistaki K, Charalampopoulos A, Vokou D, Kassomenos P, Damialis A (2020) Identifying patterns of airborne pollen distribution using a synoptic climatology approach. Science of The Total Environment 714:136625. 10.1016/j.scitotenv.2020.136625

Pedretti S, Sittmann A, Von Hagen A, Peter J (2025) Analysis of the multicomponent ALEX array data to examine patterns of sensitization in Cape Town, South Africa. Frontiers in Allergy 6:1572509

Pershad AR, Krishnan R, Lee E, Gardiner L, Hughes E, Tummala N (2025) How Climate Change Is Impacting Allergic Rhinitis: A Scoping Review. Laryngoscope 135 (8):2670–2682. doi:10.1002/lary.32124

Raissouni I, Boullayali A, Recio M, Bouziane H (2024) Variations, trends and forecast models for the airborne Olea europaea pollen season in Tétouan (NW of Morocco). International Journal of Biometeorology 68 (12):2613–2625. doi:10.1007/s00484-024-02772-9

Schramm PJ, Brown C, Saha S, Conlon K, Manangan A, Bell J, Hess J (2021) A systematic review of the effects of temperature and precipitation on pollen concentrations and season timing, and implications for human health. International journal of biometeorology 65 (10):1615–1628

Sedghy F, Varasteh AR, Sankian M, Moghadam M (2018) Interaction Between Air Pollutants and Pollen Grains: The Role on the Rising Trend in Allergy. Rep Biochem Mol Biol 6 (2):219–224

Seedat R, Claassen J, Claassen A, Joubert G (2010) Mite and cockroach sensitisation in patients with allergic rhinitis in the Free State. South African Medical Journal 100 (3):160–163

Seedat RY, Sujee M, Ismail W, Vallybhai NY, Cassim MI, Khan S, Solwa A, Joubert G (2018) Allergic rhinitis in medical students at the University of the Free State. South African Family Practice 60 (4):121–125. doi:10.1080/20786190.2018.1437869

Van Haeften S, Campbell BC, Milic A, Addison-Smith E, Al Kouba J, Huete A, Beggs PJ, Davies JM (2024) Environmental DNA analysis of airborne poaceae (grass) pollen reveals taxonomic diversity across seasons and climate zones. Environmental Research 247:117983

Zar HJ, Ehrlich RI, Workman L, Weinberg EG (2007) The changing prevalence of asthma, allergic rhinitis and atopic eczema in African adolescents from 1995 to 2002. Pediatric Allergy and Immunology 18 (7):560-565. 10.1111/j.1399-3038.2007.00554.x

Zhaobin S, Yuxin Z, Xingqin A, Na G, Ziming L, Shuwen Z, Yinglin L, Wenxi R, Yaqin B, Jingyi X, Shihong L (2024) Effects of airborne pollen on allergic rhinitis and asthma across different age groups in Beijing, China. Sci Total Environ 912:169215. doi:10.1016/j.scitotenv.2023.169215

