## Supplementary material for "Five years of national airborne pollen monitoring in South Africa: Biome-specific calendars to inform allergy diagnosis and prevention": https://uctcloud-my.sharepoint.com/:w:/g/personal/mtvtak006_myuct_ac_za/IQB4iXi5mvNhQ5Fq_o_7XqmkAYtUsPkPsbpbD2lu8ZploJk

**SUPPLEMENTARY FIGURES**

**Fig. S1** Daily pollen concentrations from 2019 to 2024 for the seven biomes (A to G) across South Africa. For each biome, the original data with gaps is shown in (i), and the data after gaps were filled using weekly averages is shown in (ii).

(a) **FYNBOS BIOME (CAPE TOWN)**

(i)

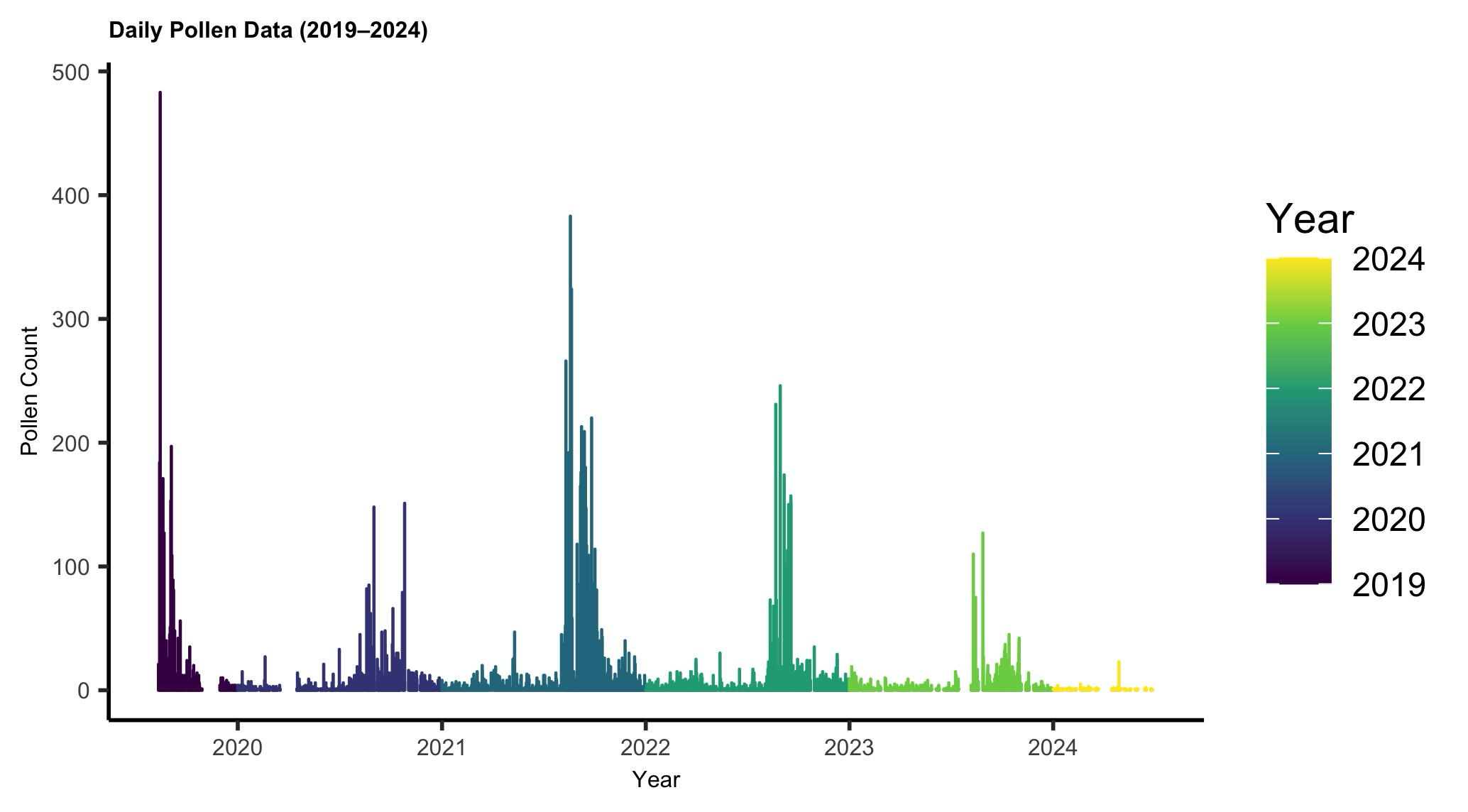

(ii)

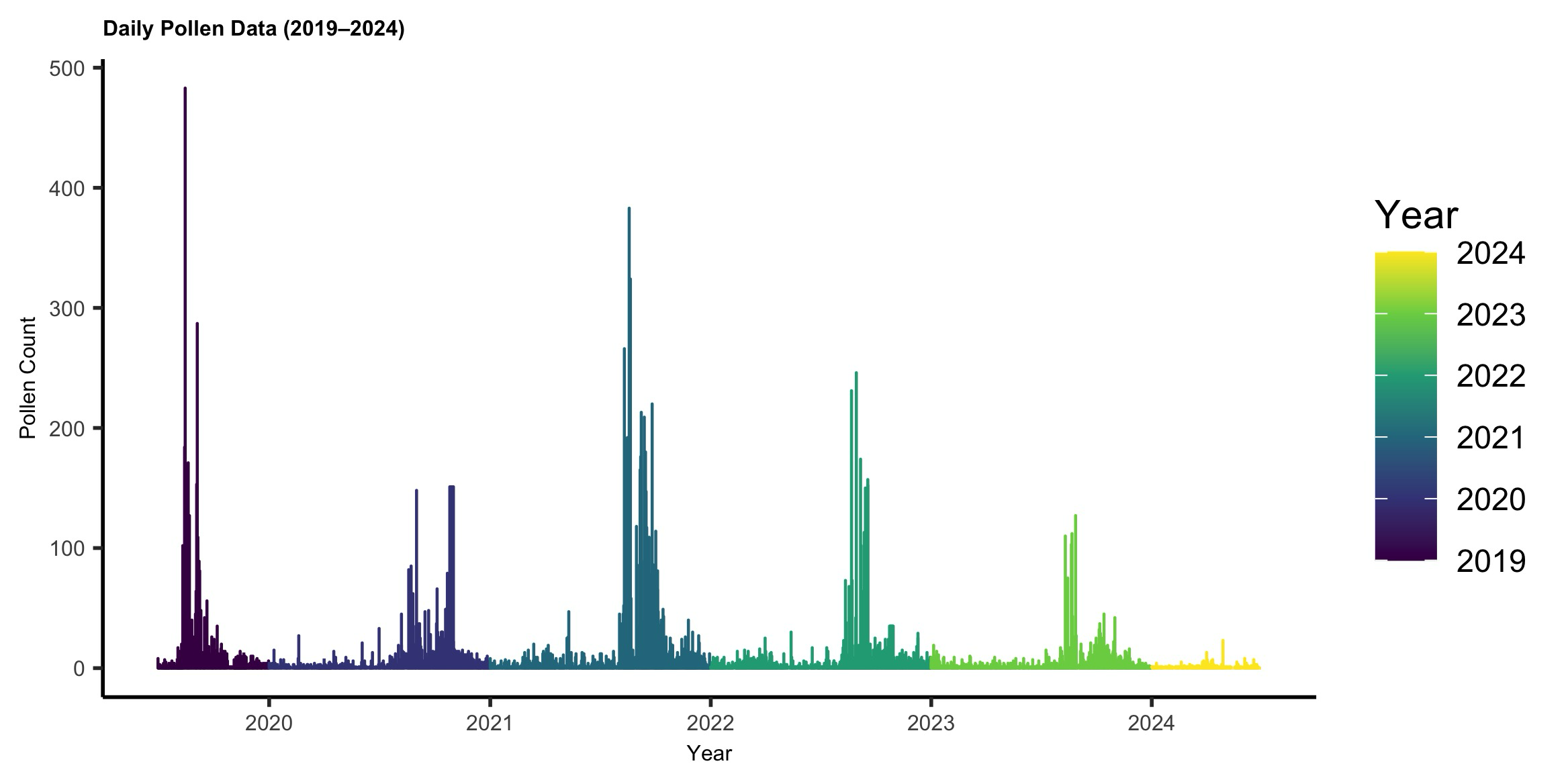

(b) **SAVANNA BIOME (PRETORIA)**

(i)

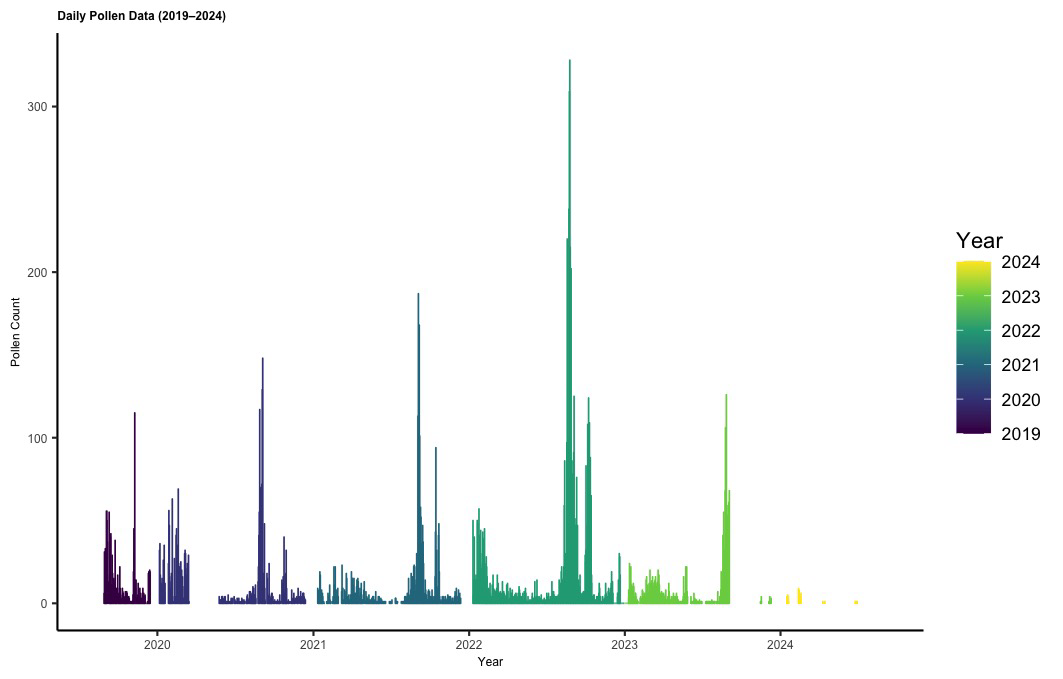

(ii)

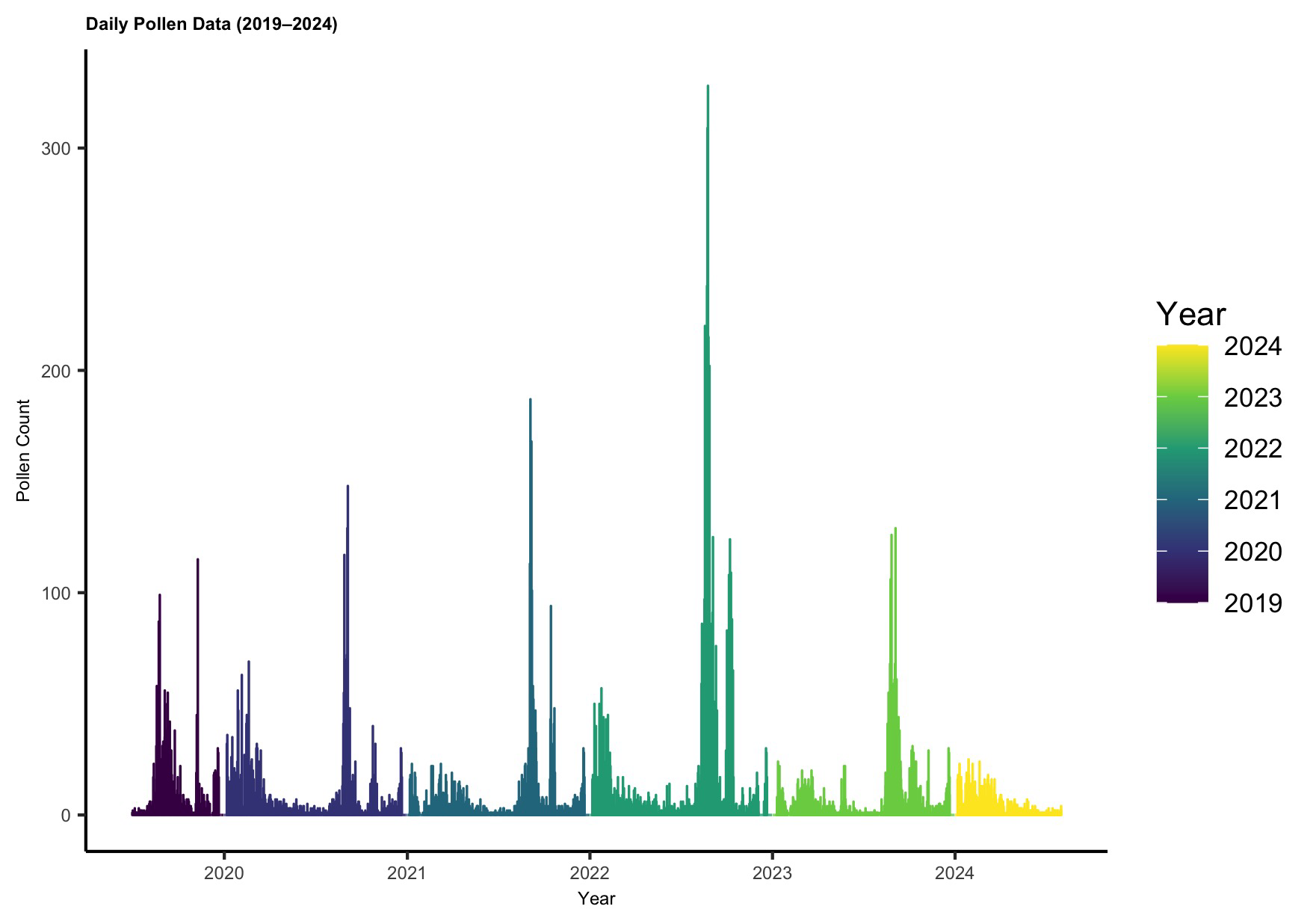

(c) **GRASSLAND BIOME (JOHANNESBURG)**

**(i)**

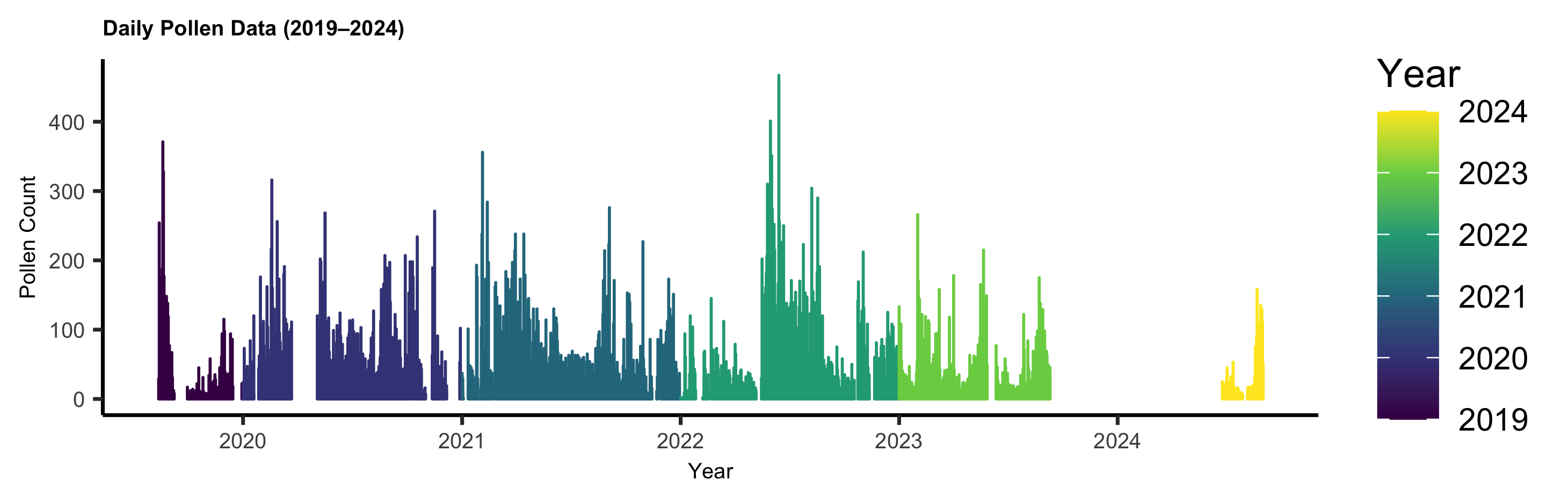
(ii)

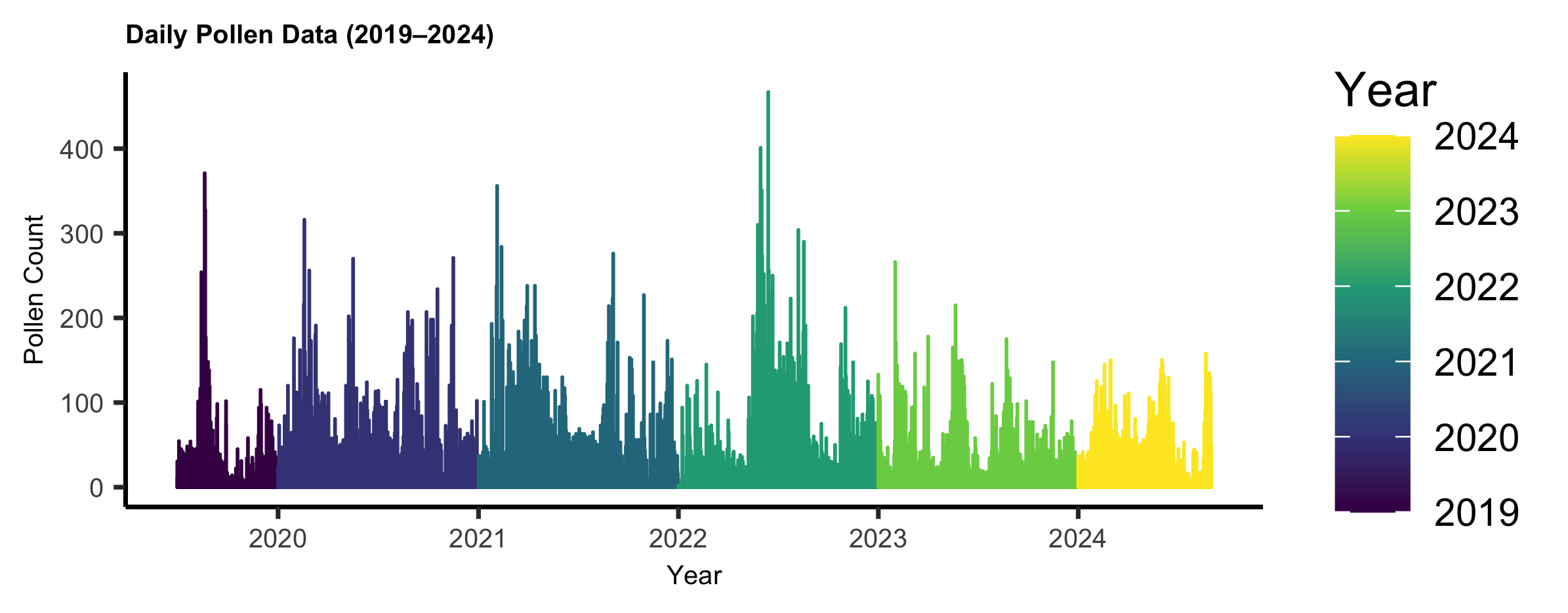

(d) **INDIAN OCEAN COASTAL BELT BIOME (DURBAN)**

**(i)**

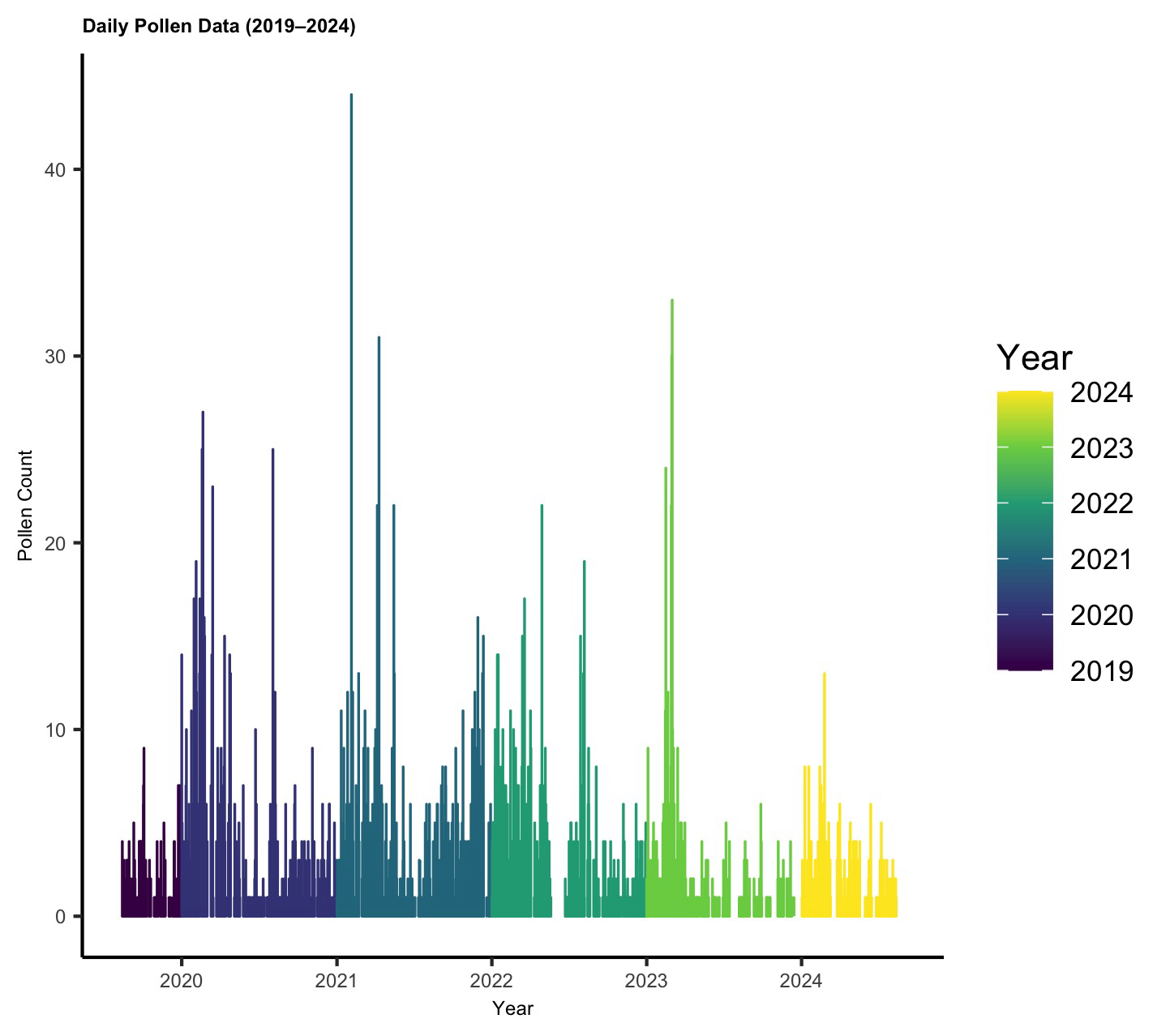

(ii)

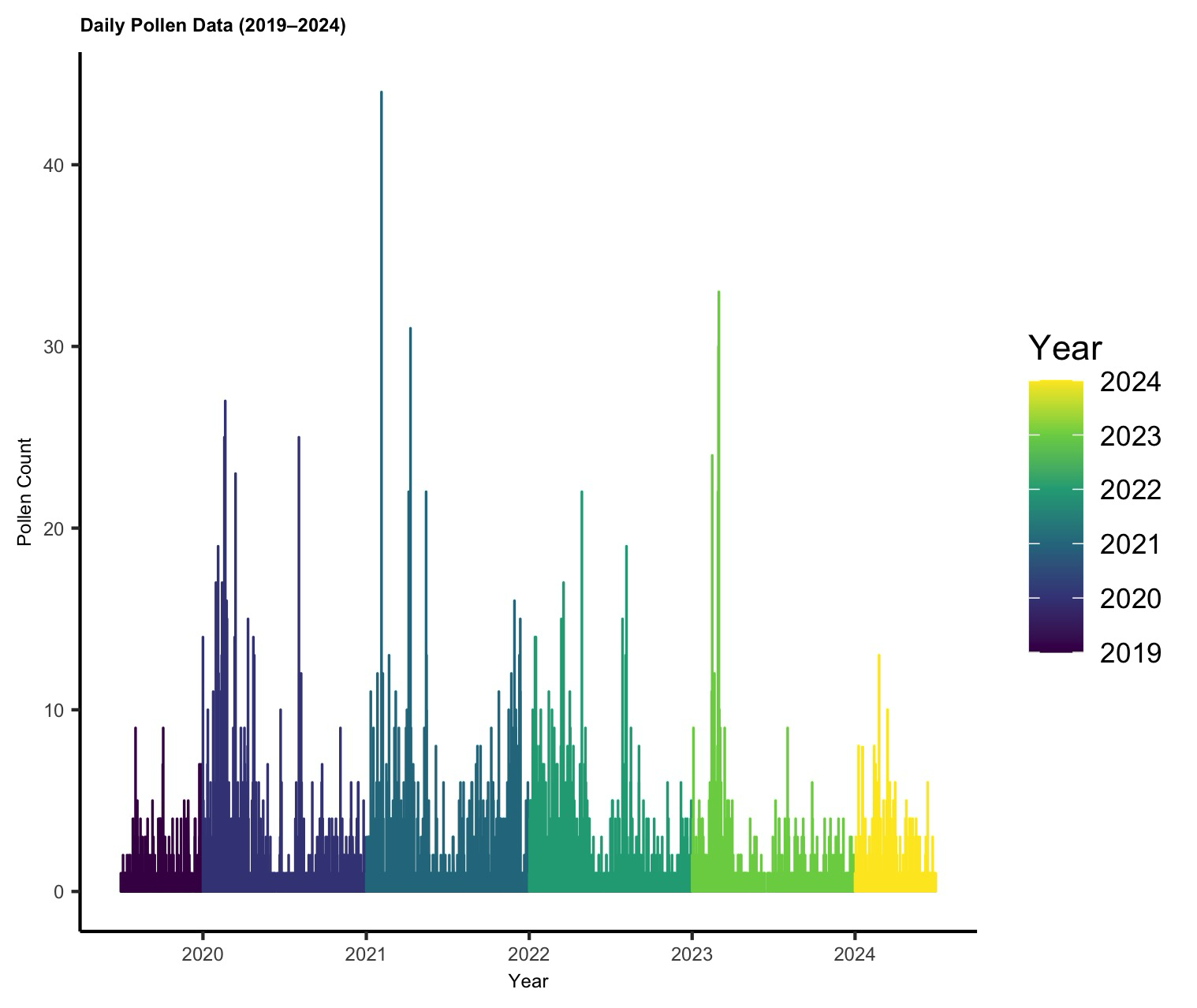

(e) **SAVANNA BIOME (KIMBERLEY)**

**(i)**

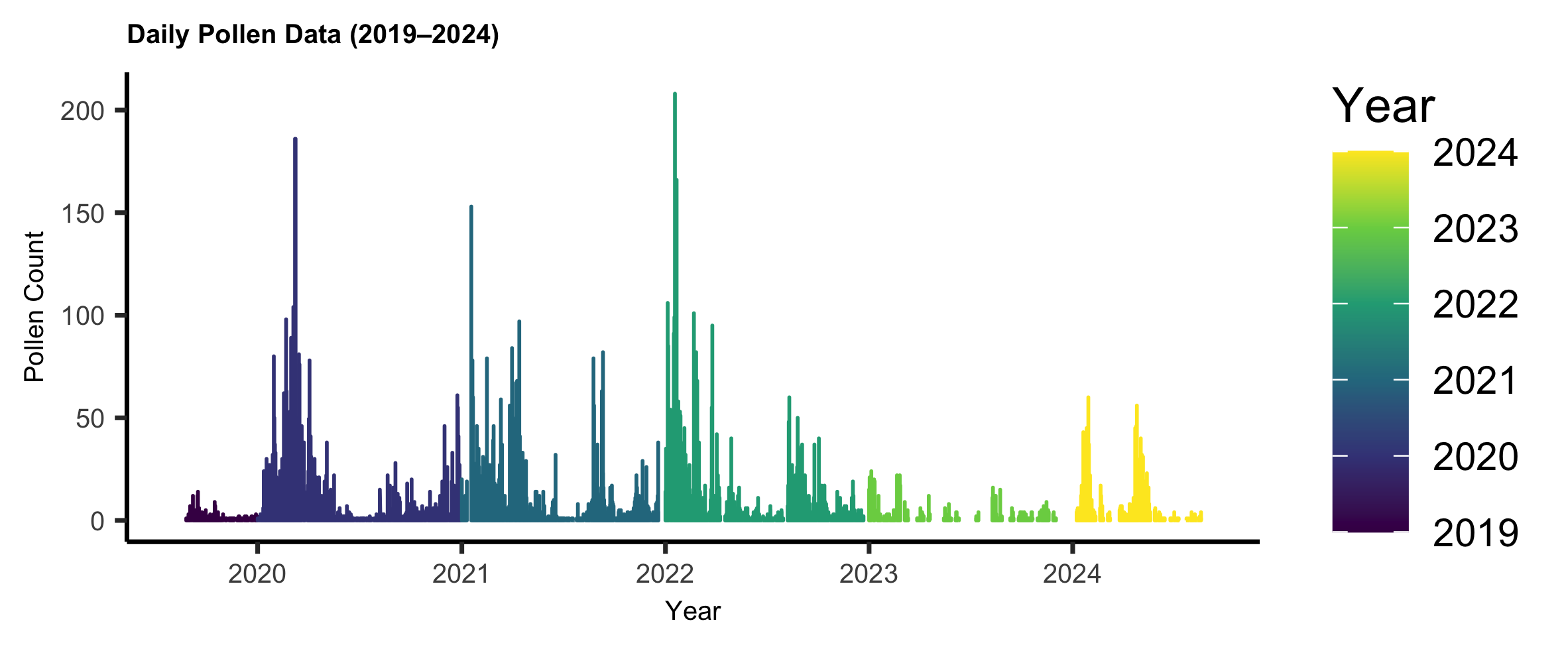

**(ii)**

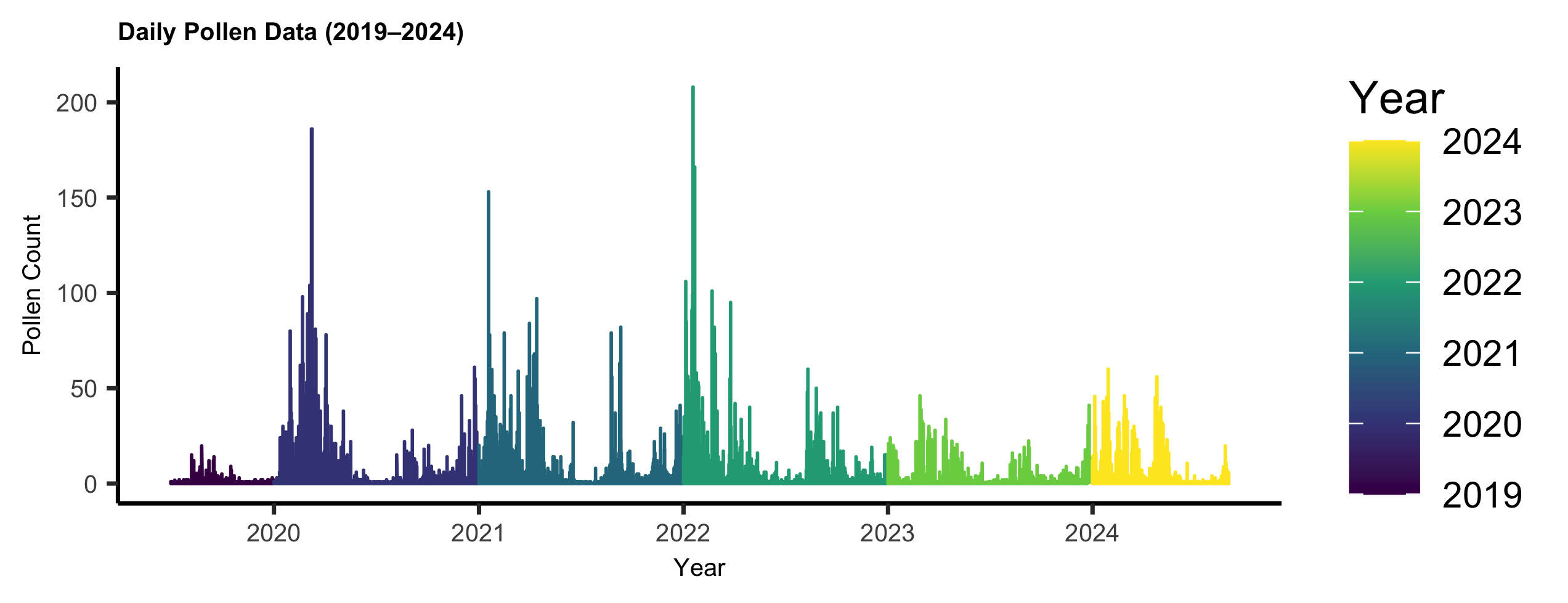

(f) **ALBANY THICKET BIOME (GQERBERHA)**

(i)

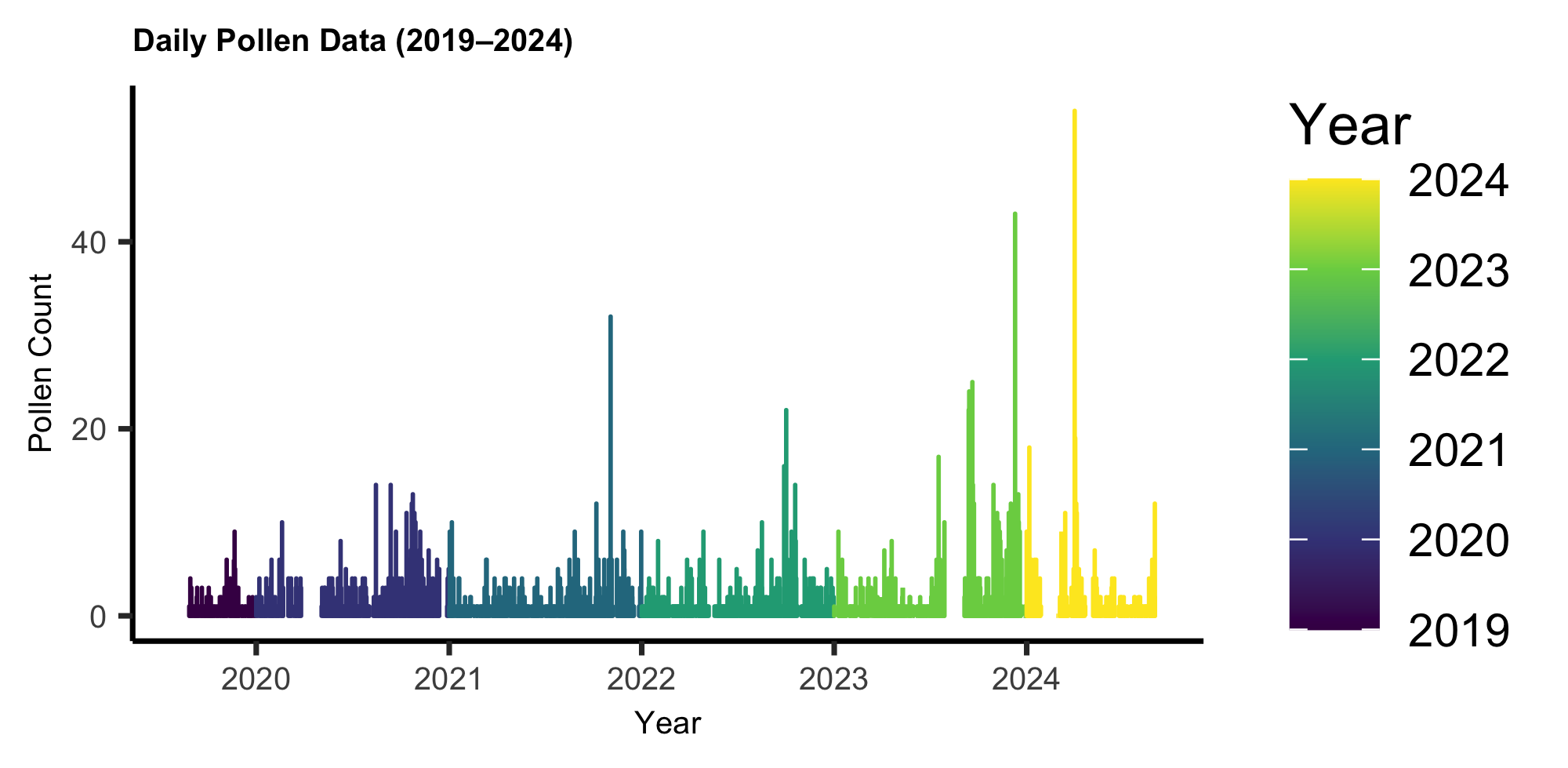

(ii)

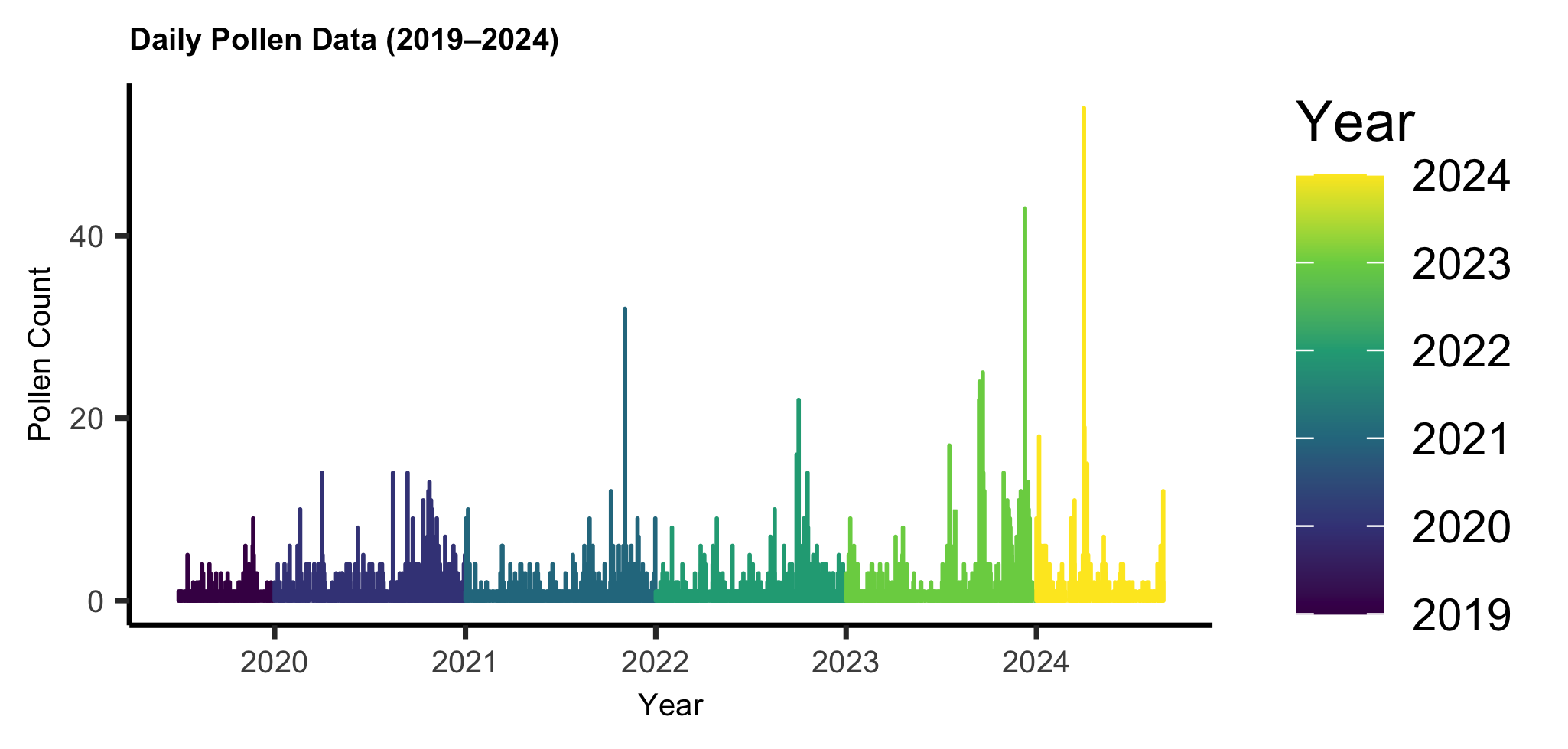

(g) **GRASSLAND (BLOEMFONTEIN)**

(i)

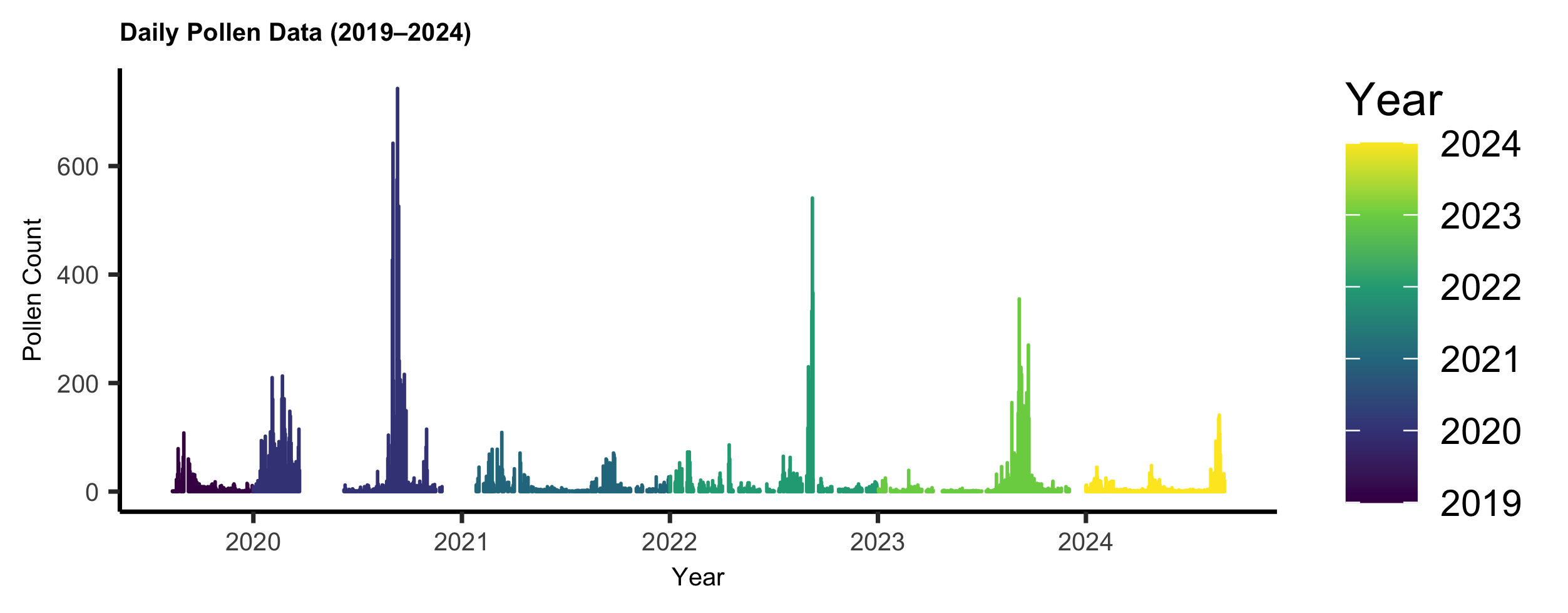

(ii)

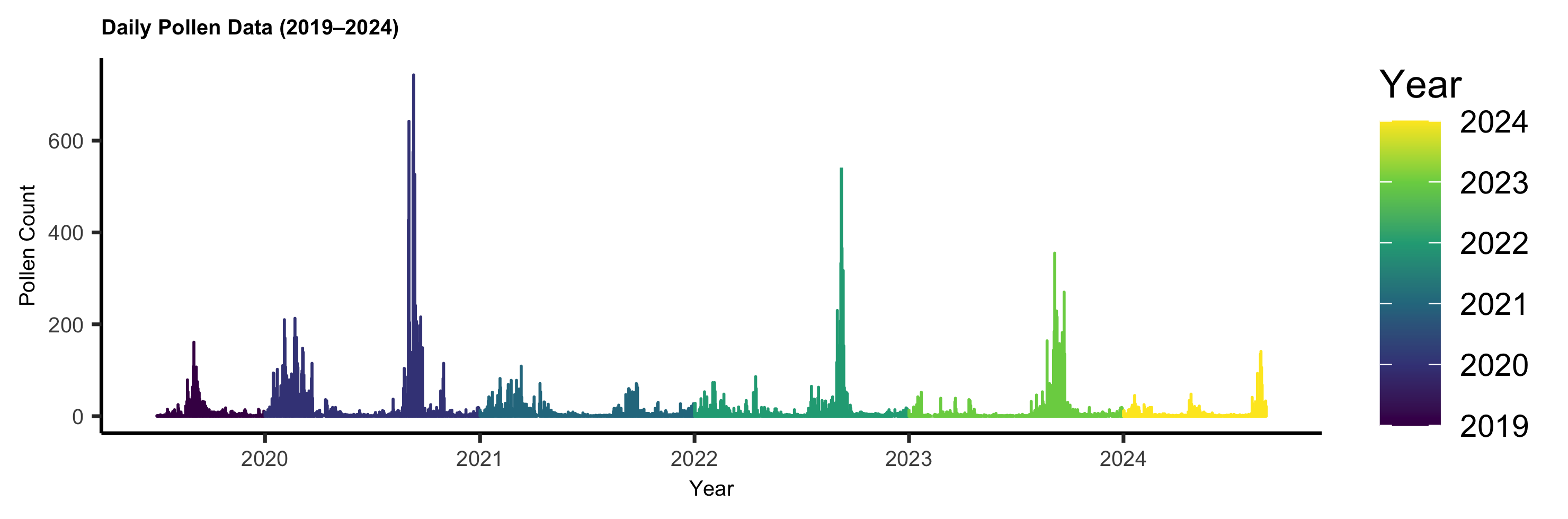

**Fig. S2** The averaged pollen integral across the cities in South Africa.

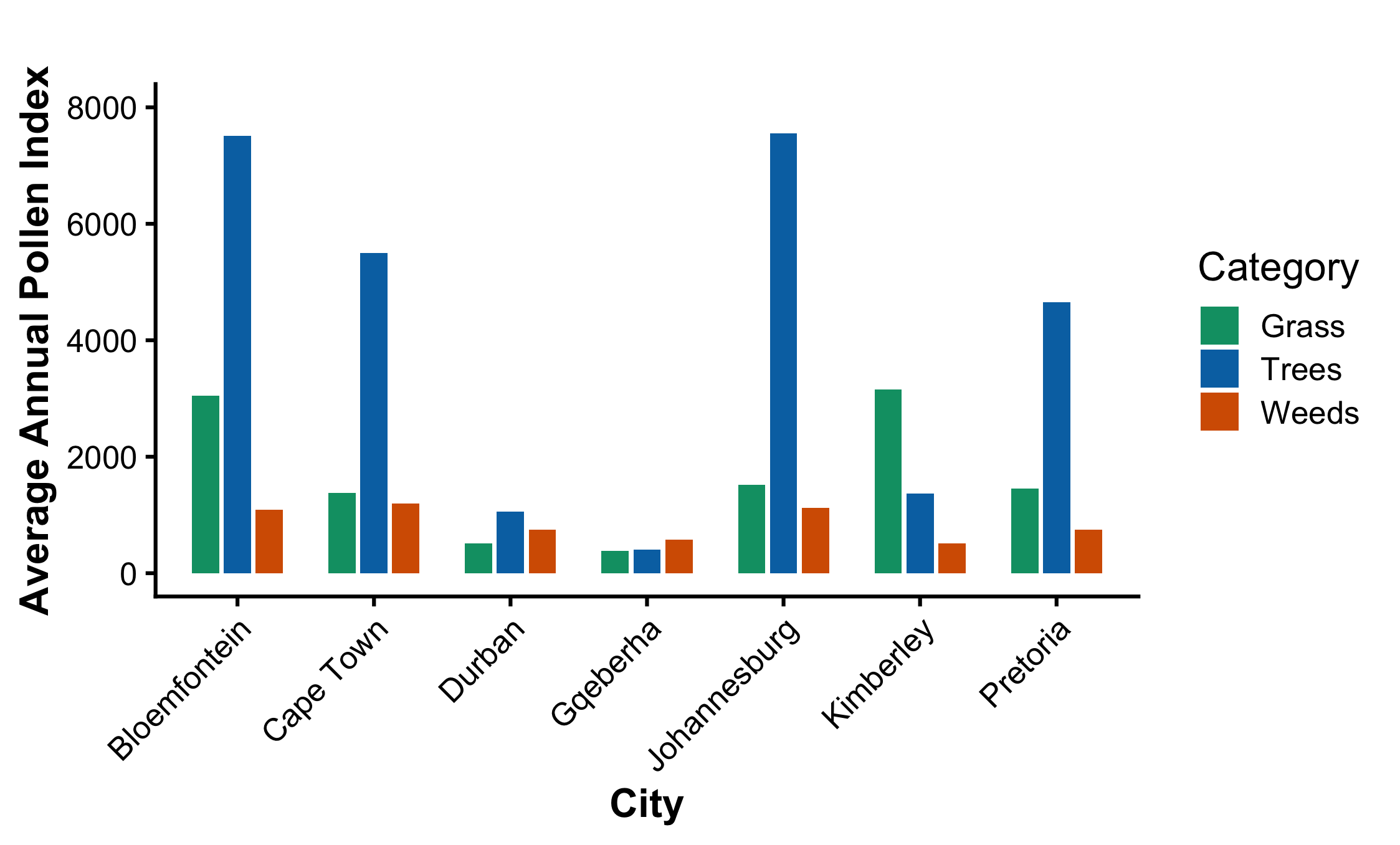

Average Annual Pollen Integral

**Fig. S3** The start day number of the pollen season by each group across the five years.

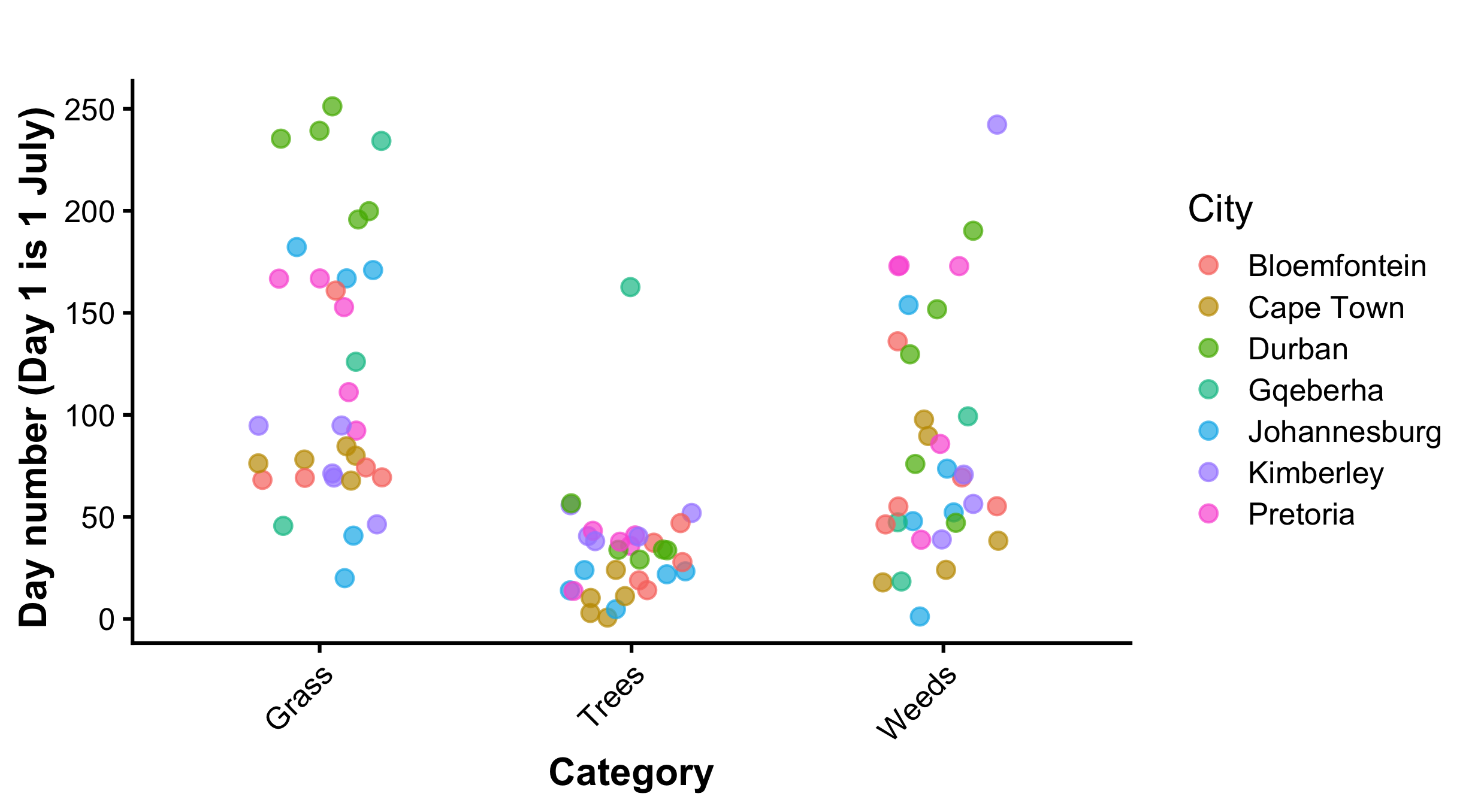

**Fig. S4** A heatmap showing climate and weather seasons of cities in South Africa

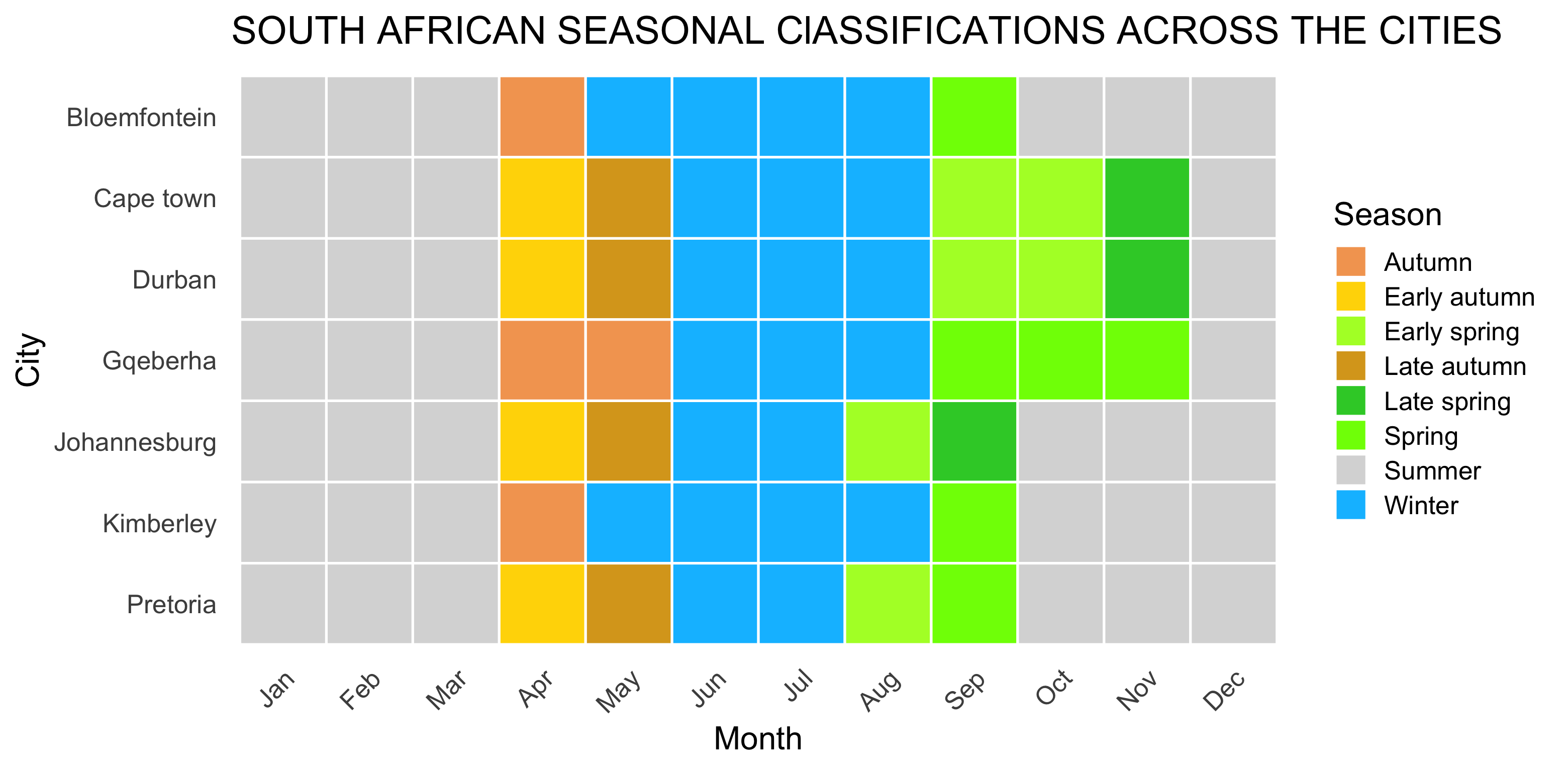

**Fig. S5** A heatmap showing grass, tree and weed pollen seasons and peak months across the cities in South Africa

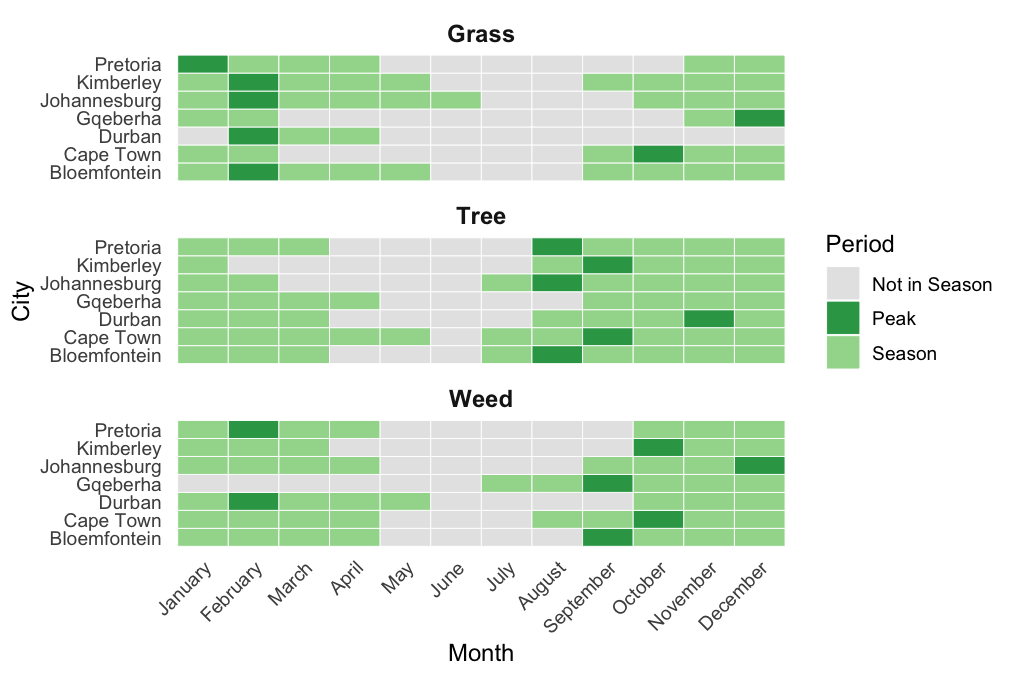

**Fig. S6** The number of pollen taxa detected at each city across the five years and grouped by month.

(a) **FYNBOS (CAPE TOWN)**

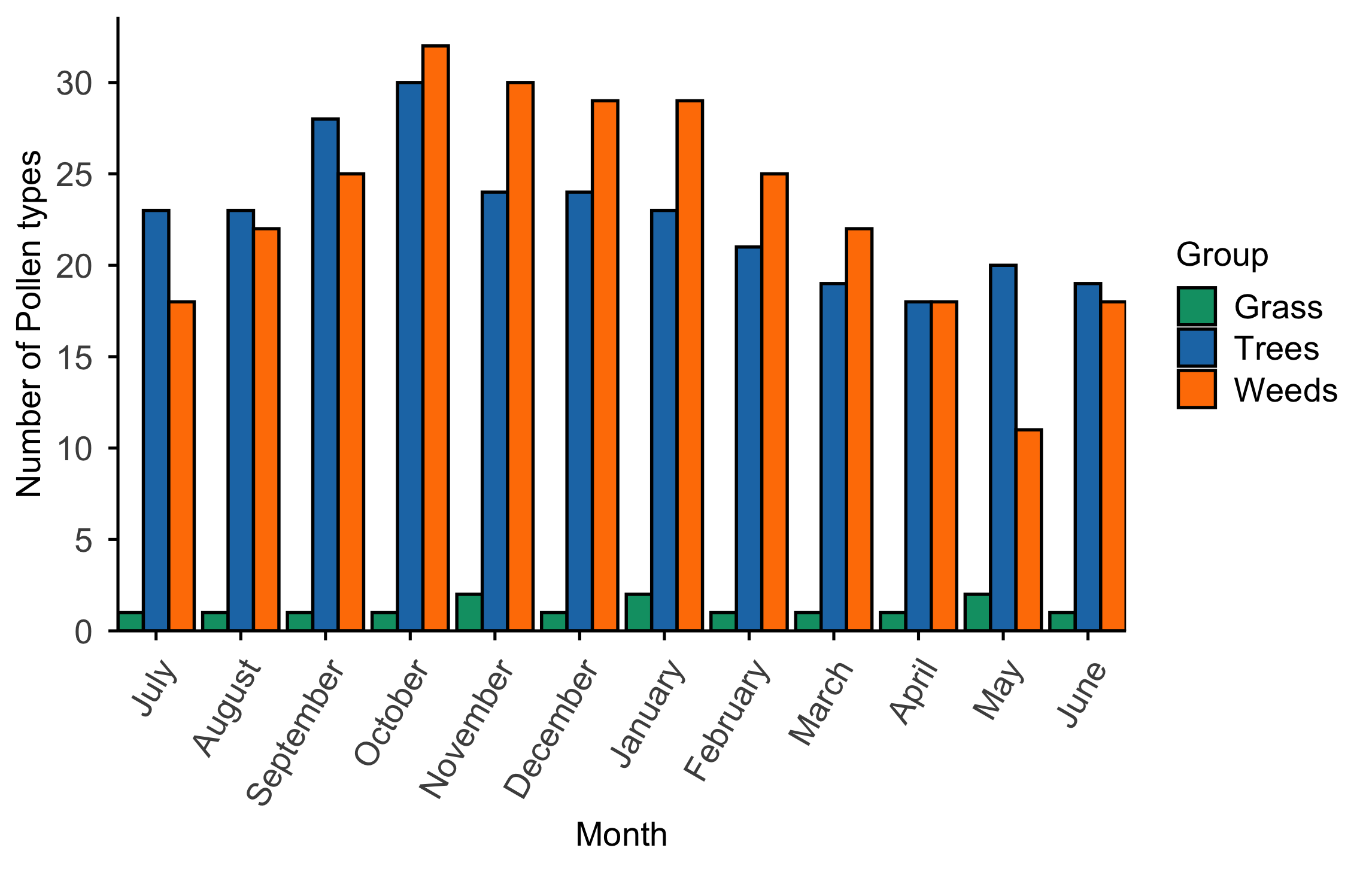

(b) **INDIAN OCEAN COASTAL BELT (DURBAN)**

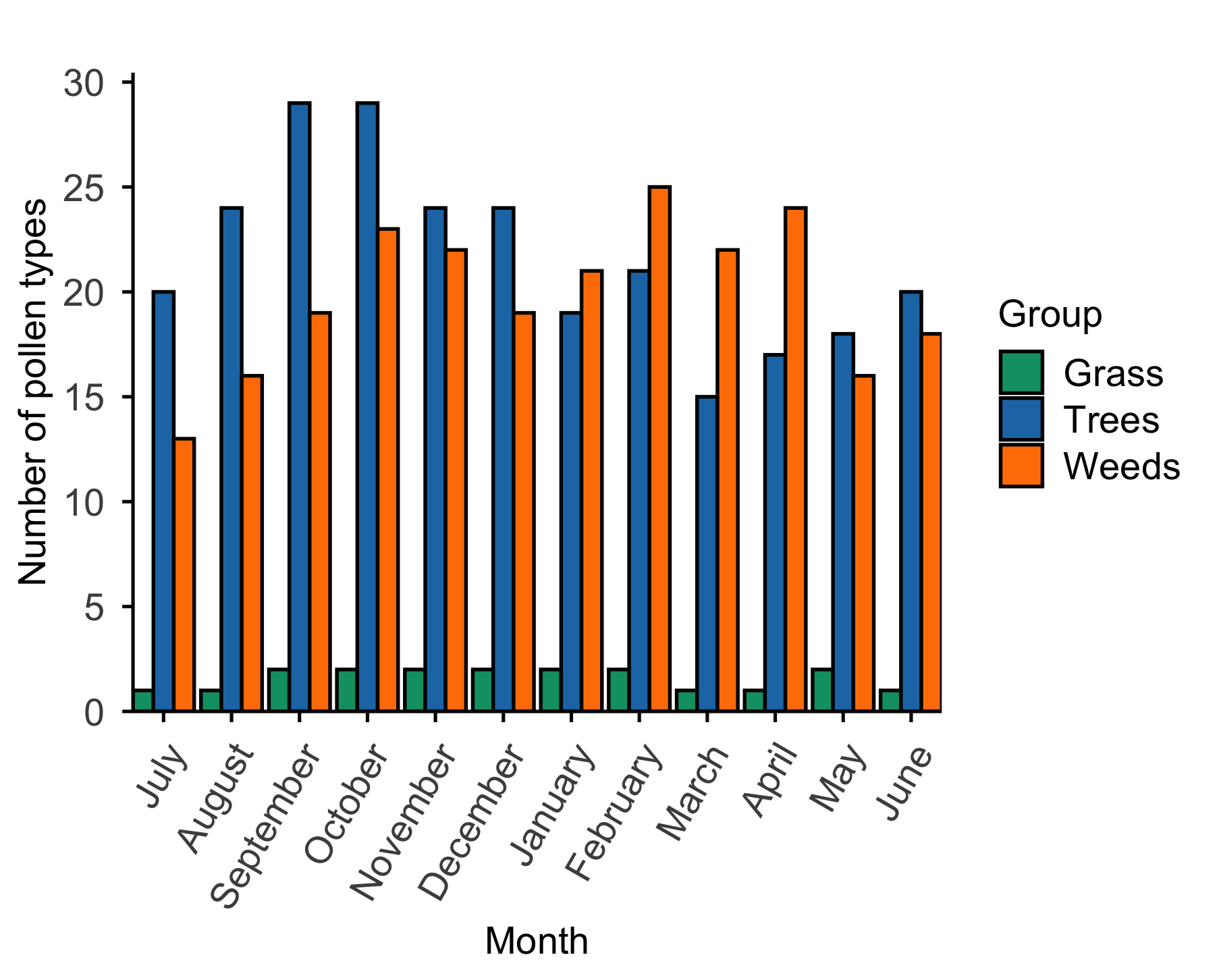

(c) **ALBANY THICKET (GQEBERHA)**

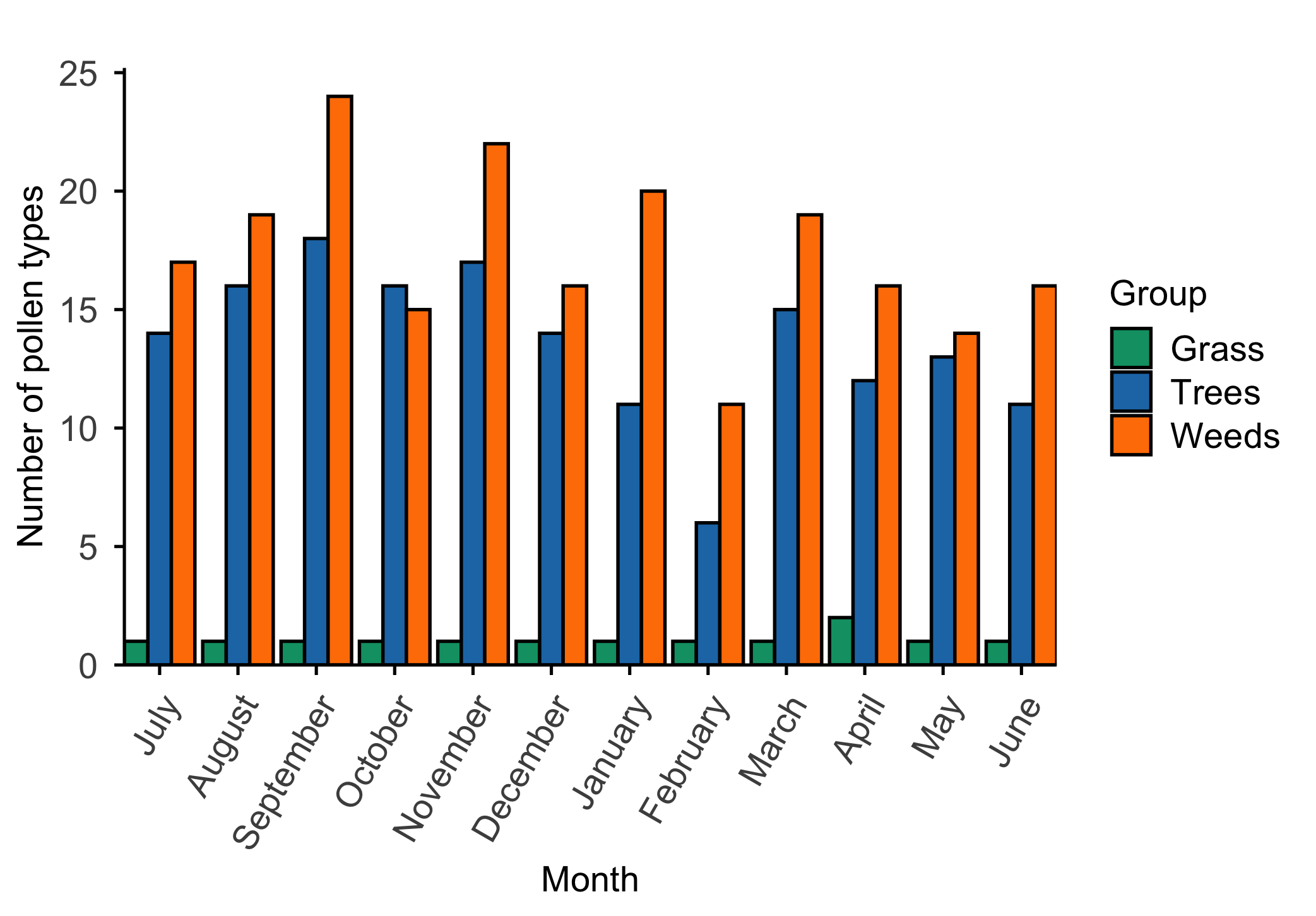

(d) **GRASSLAND (JOHANNESBURG)**

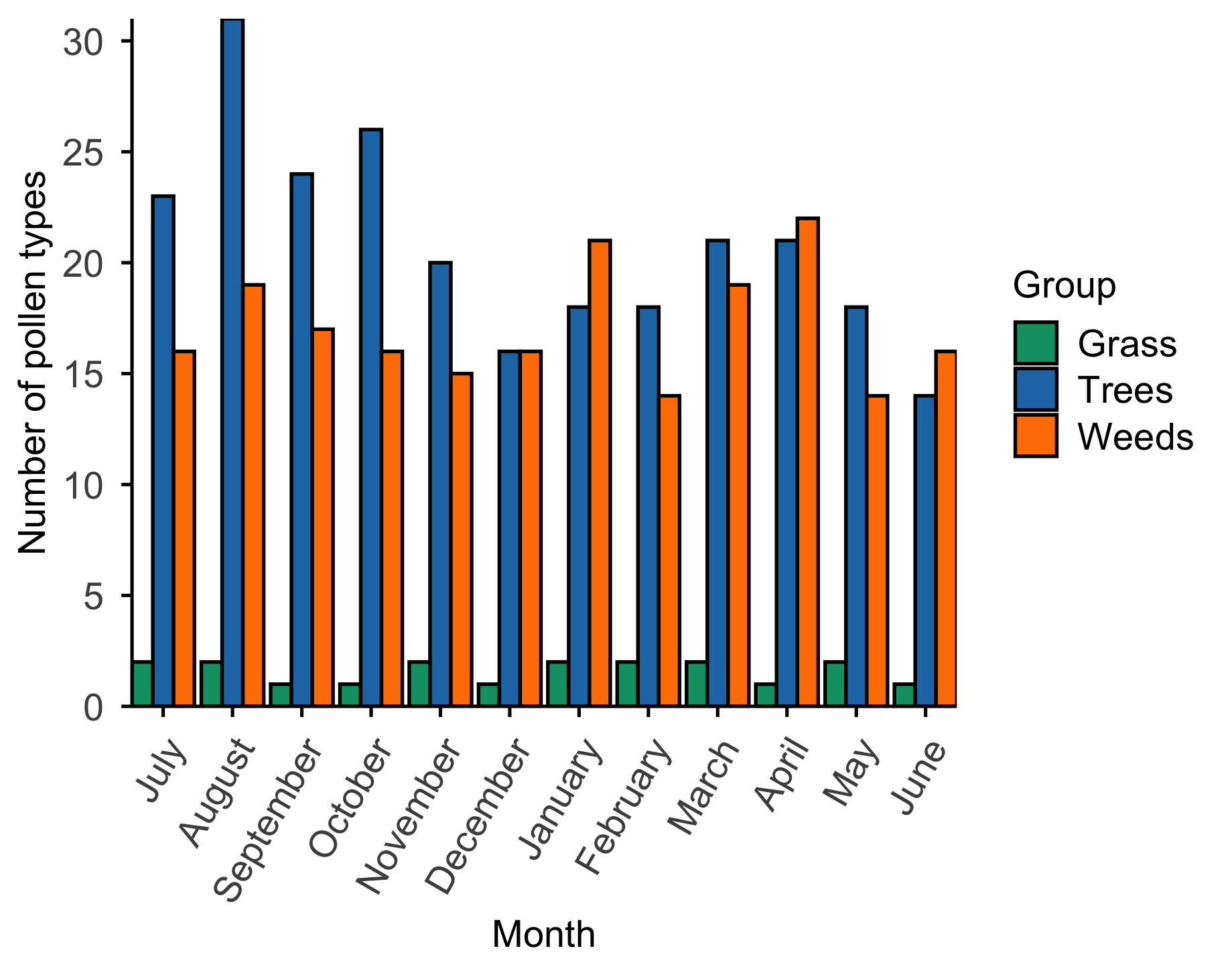

(e) **SAVANNA (PRETORIA)**

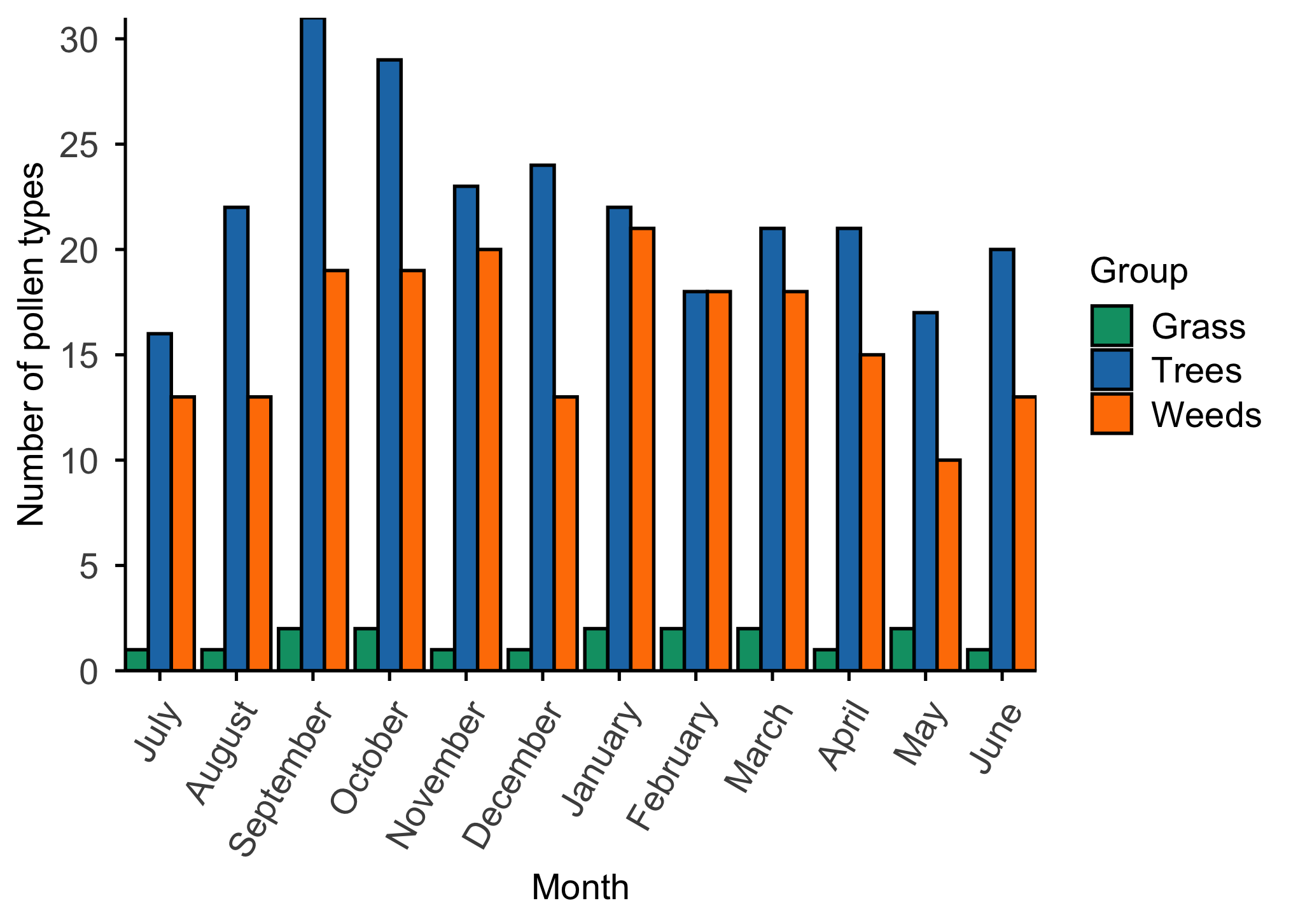

(f) **GRASSLAND (BLOEMFONTEIN)**

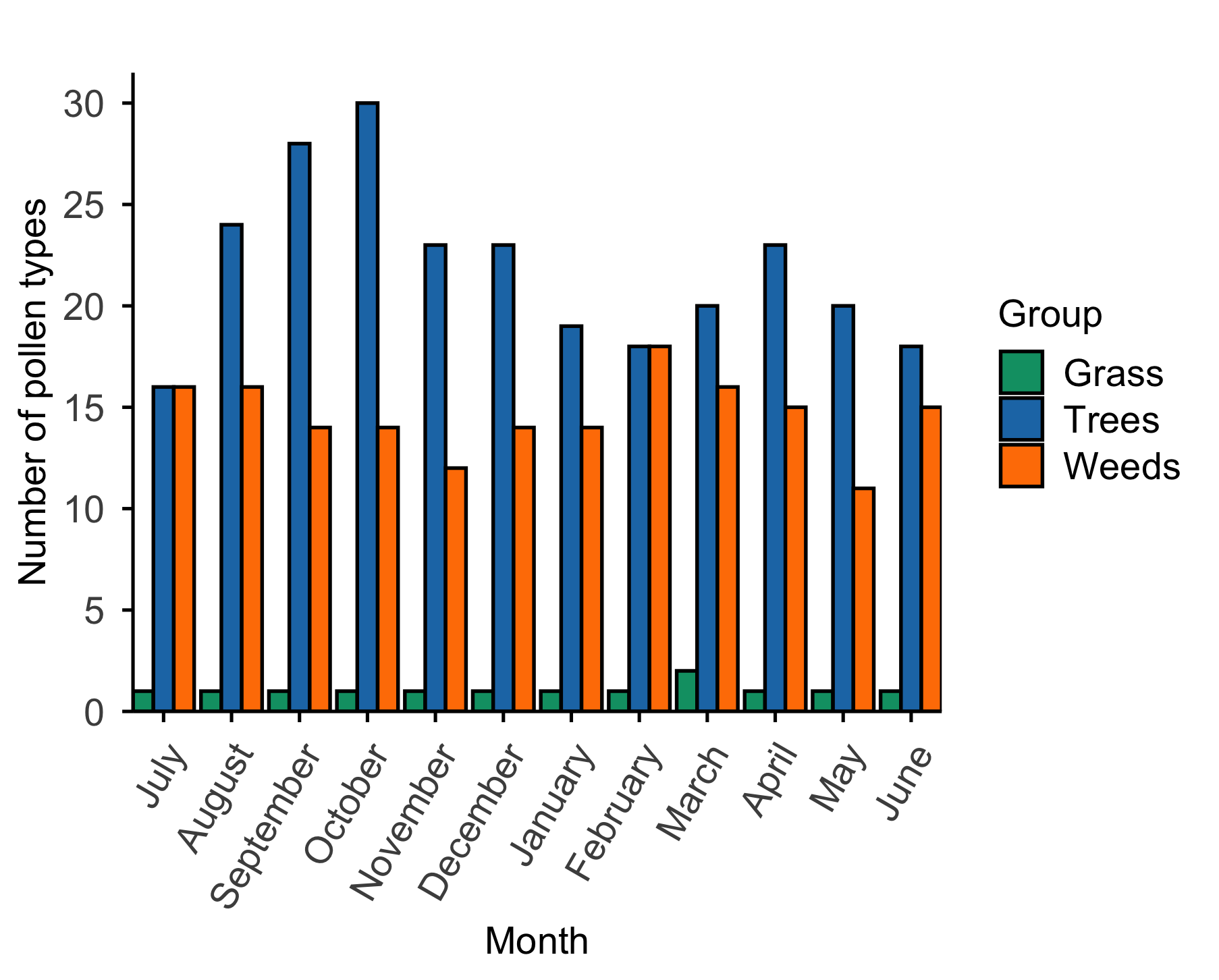

(g) **SAVANNA (KIMBERLEY)**

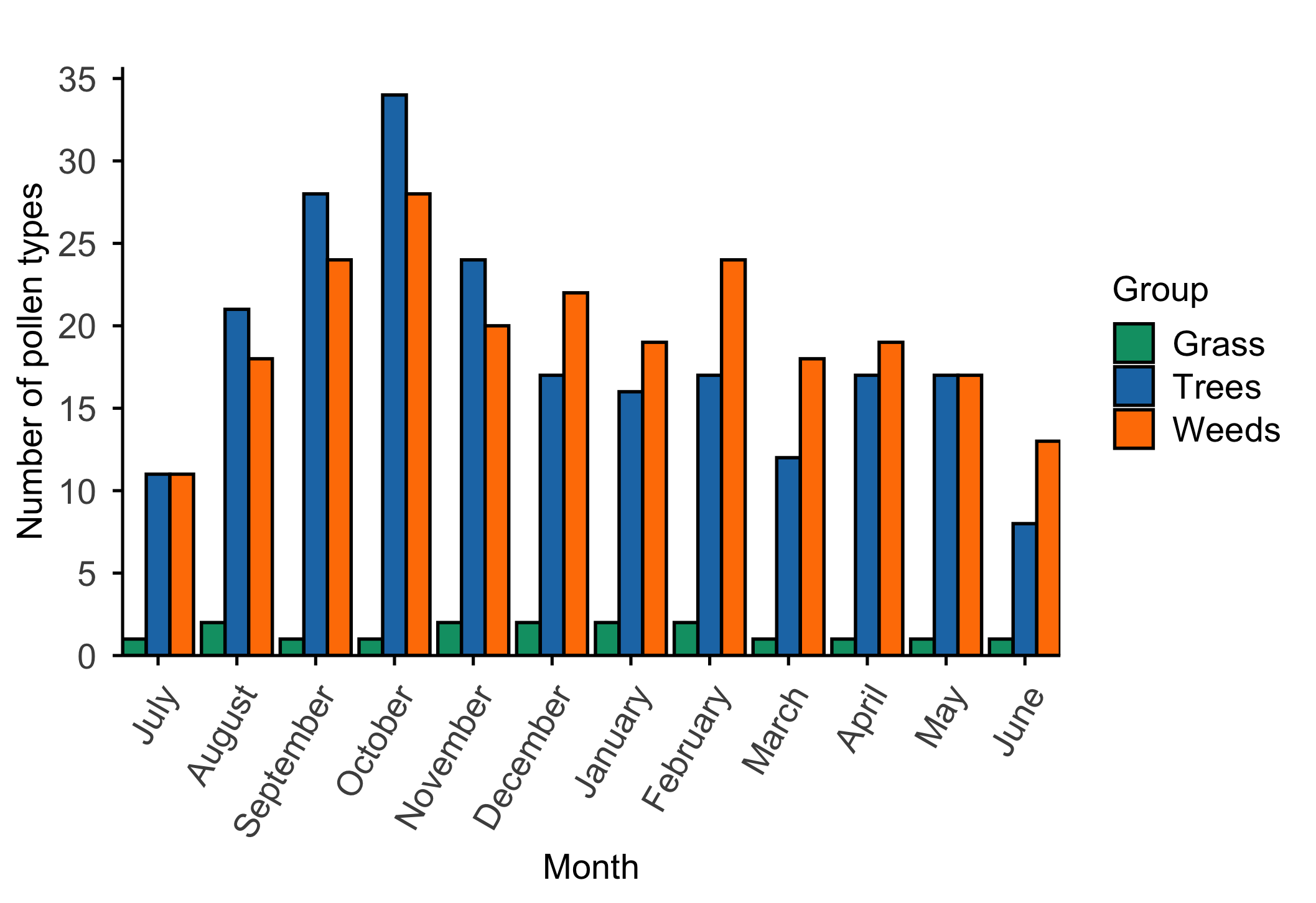

**Fig. S7** List of the top 20 grass species that occur in each province of South Africa based on the global biodiversity information facility (35).

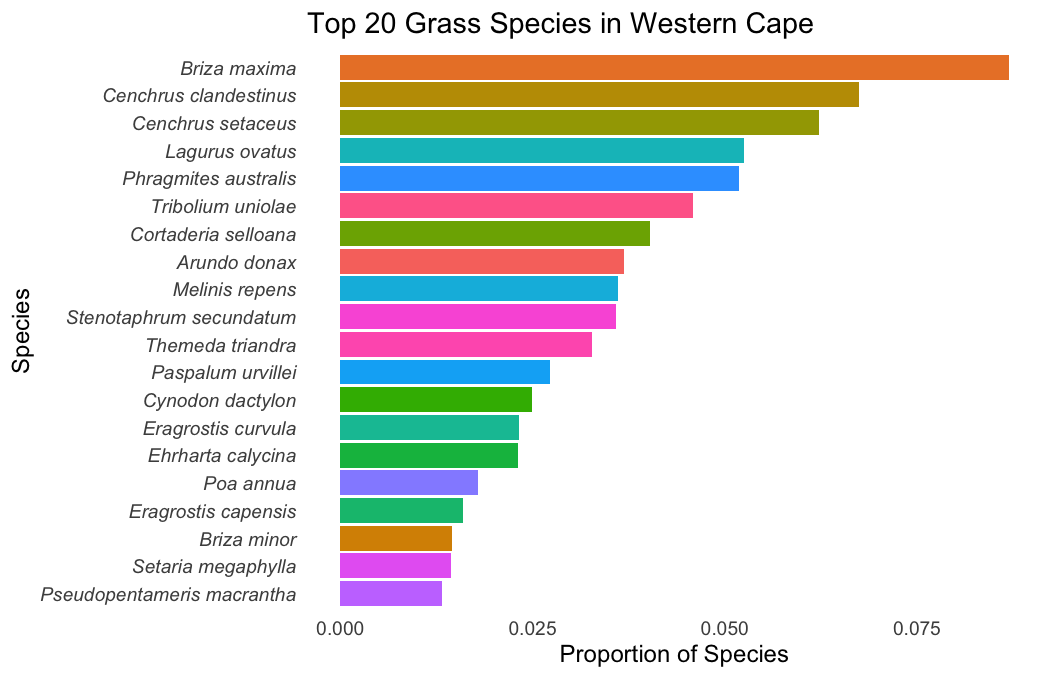

(b)

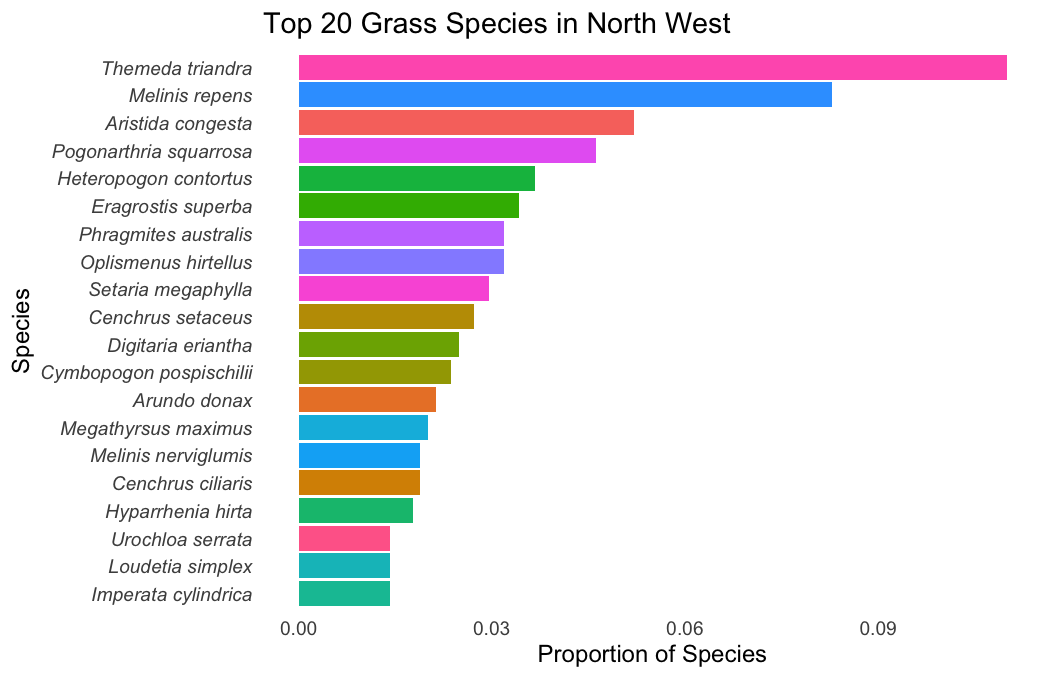

(c)

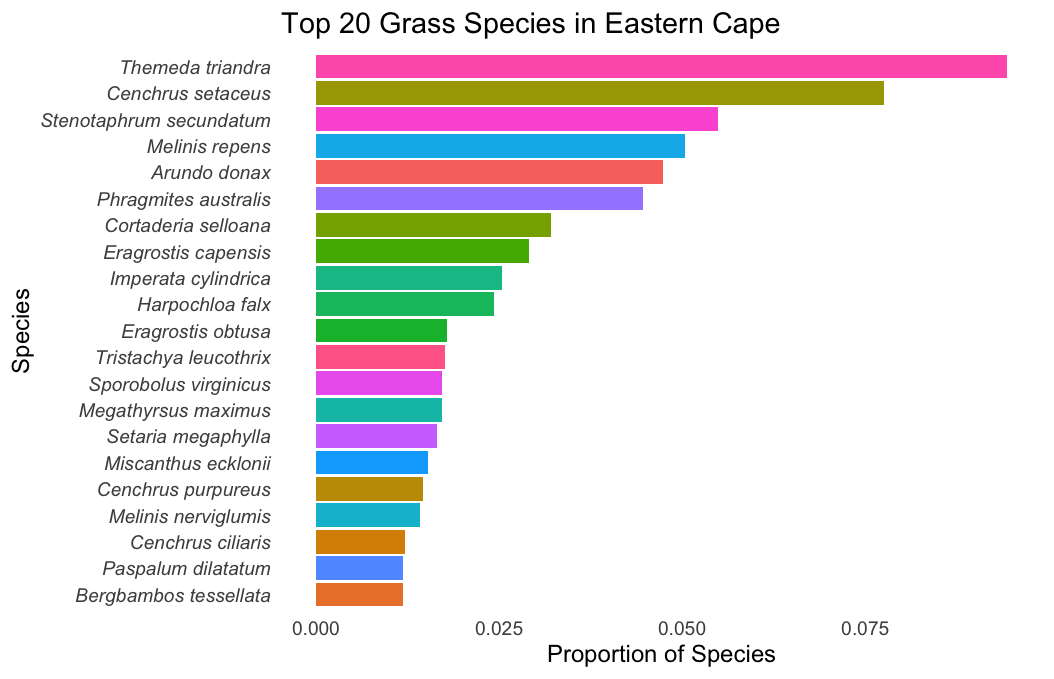

(d)

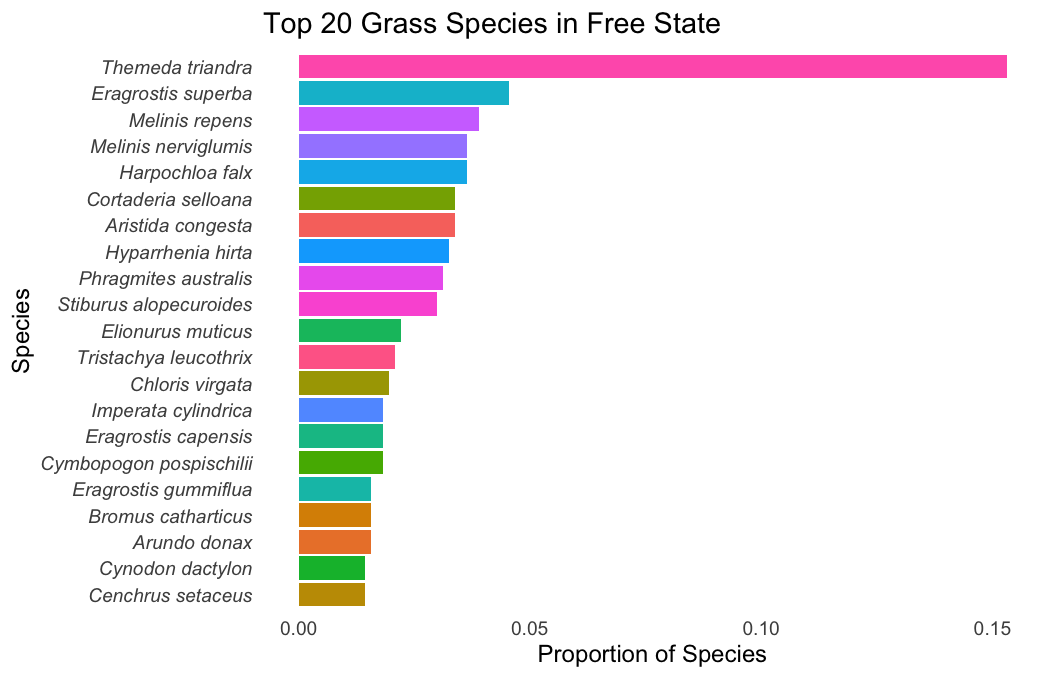

(e)

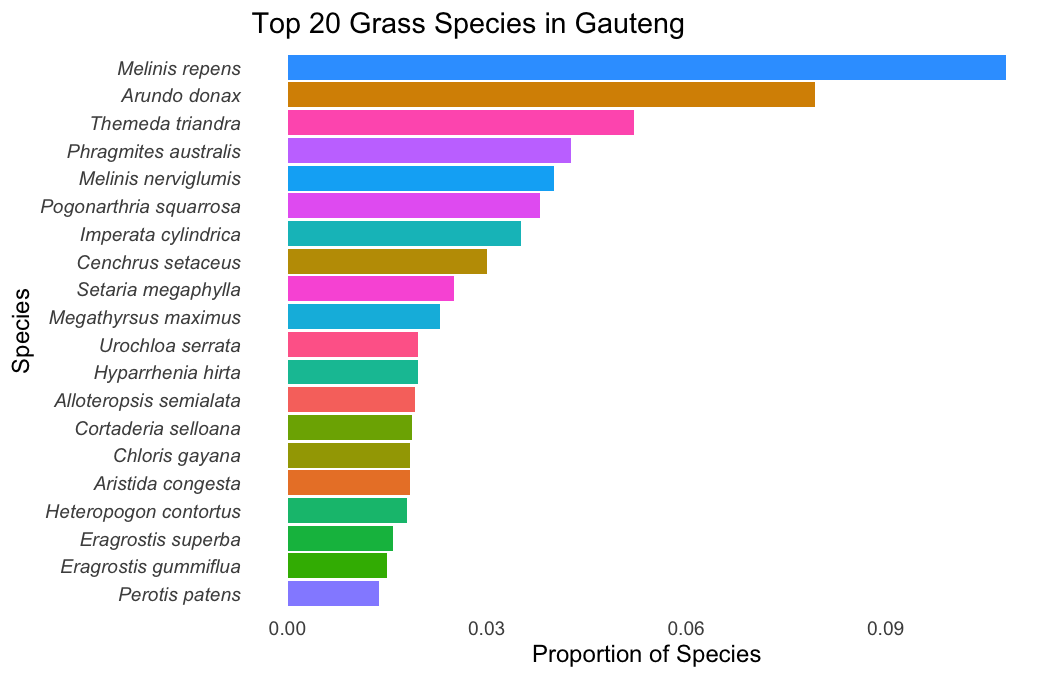

(f)

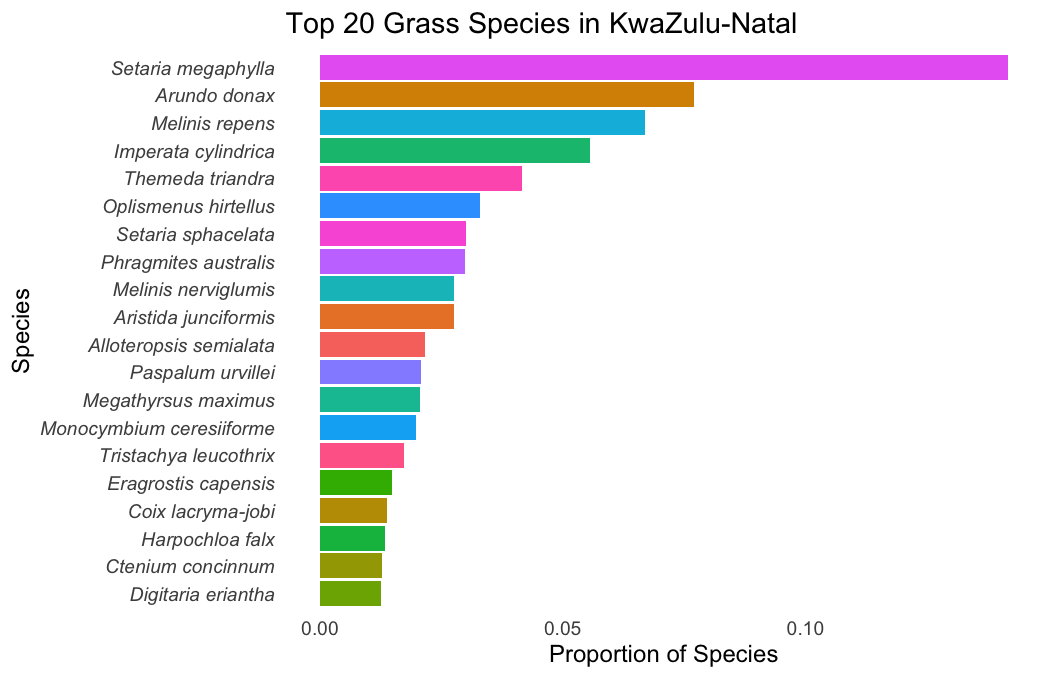

**(g)**

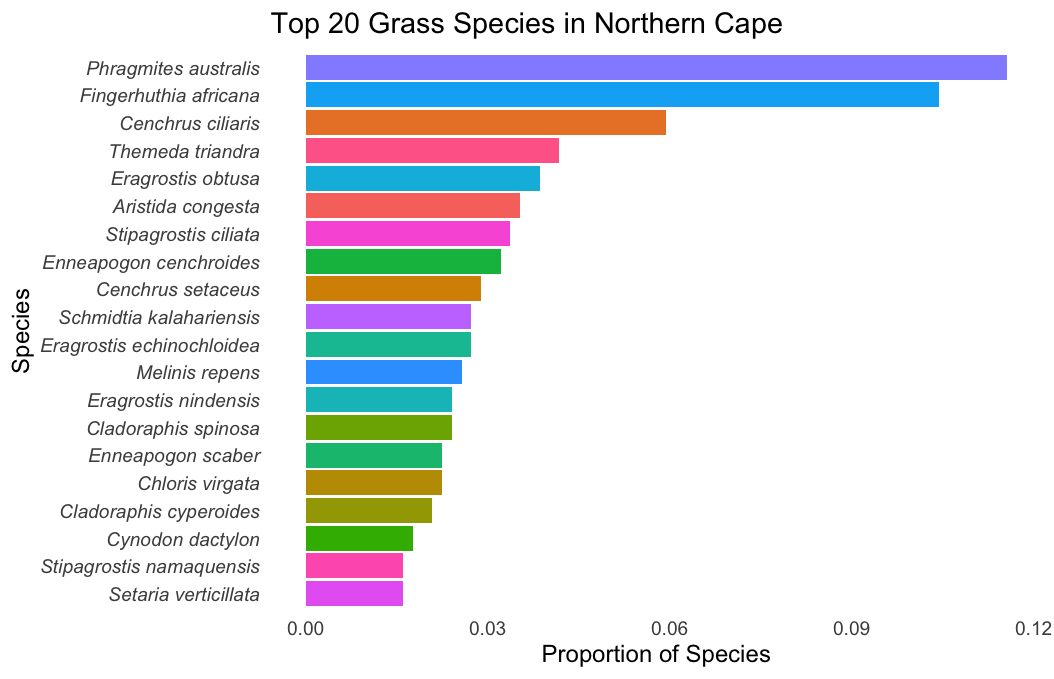

**Fig. S8** Pollen trends of taxa in seasons with high and low pollen concentrations. The graphs shows pollen concentration of different taxa in years with the highest and the lowest pollen levels (A-G).

(a) **FYNBOS (CAPE TOWN)**

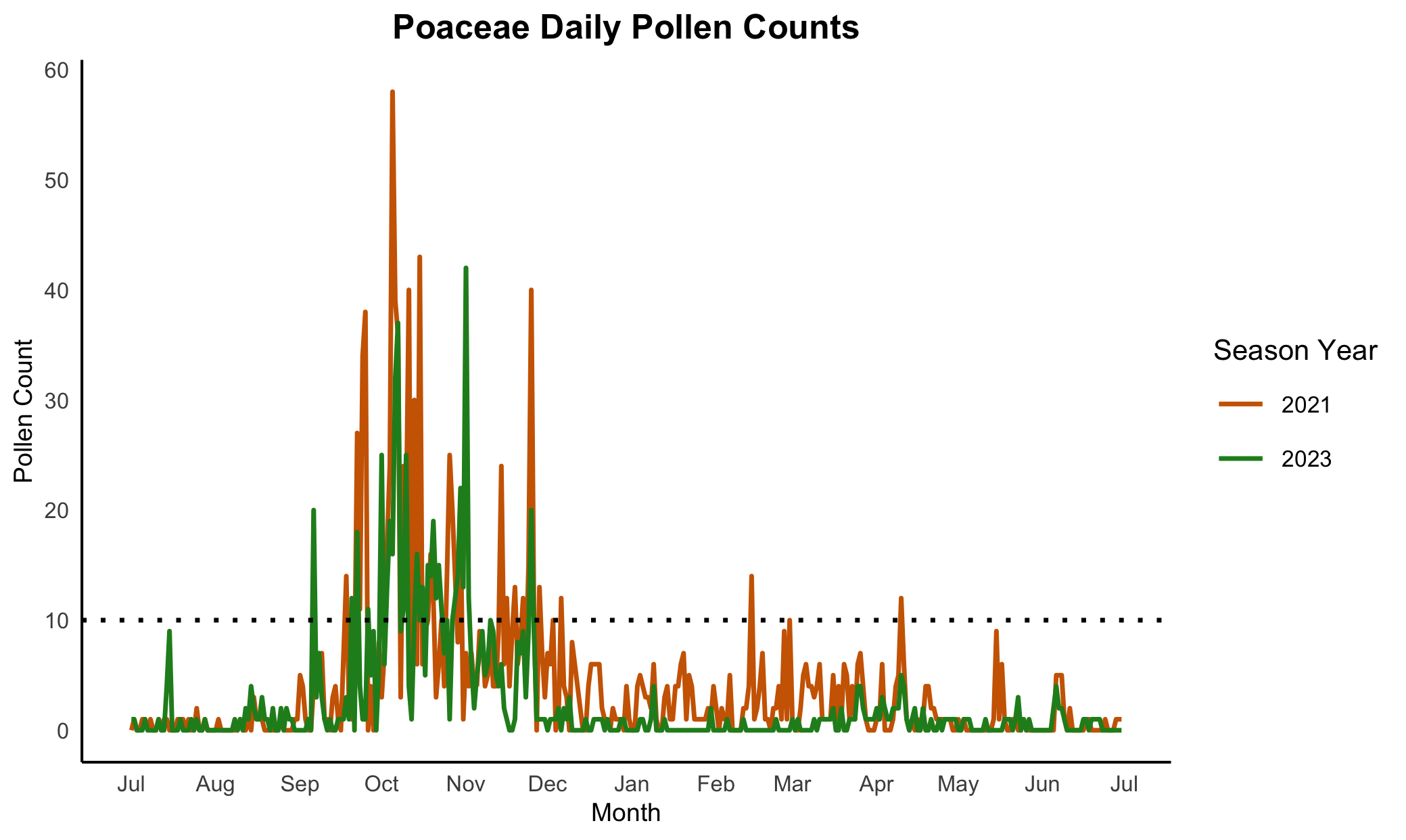

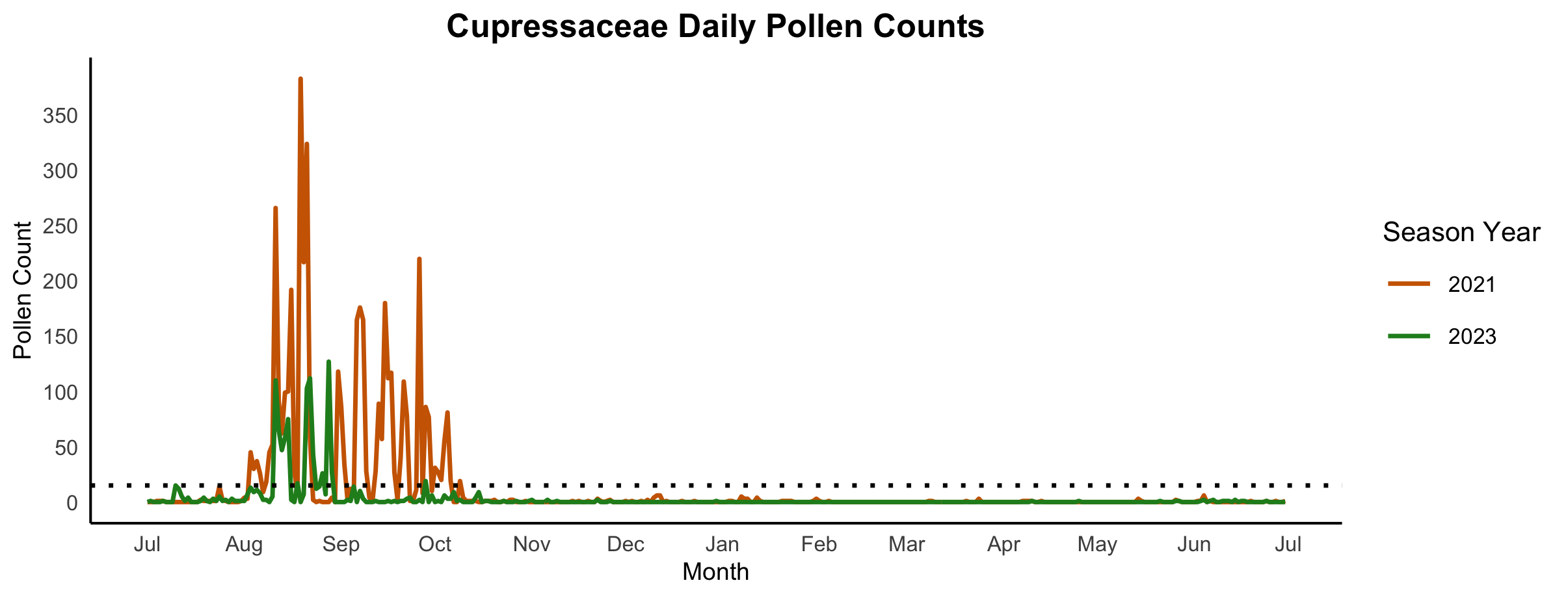

(b) **GRASSLAND (JOHANNESBURG)**

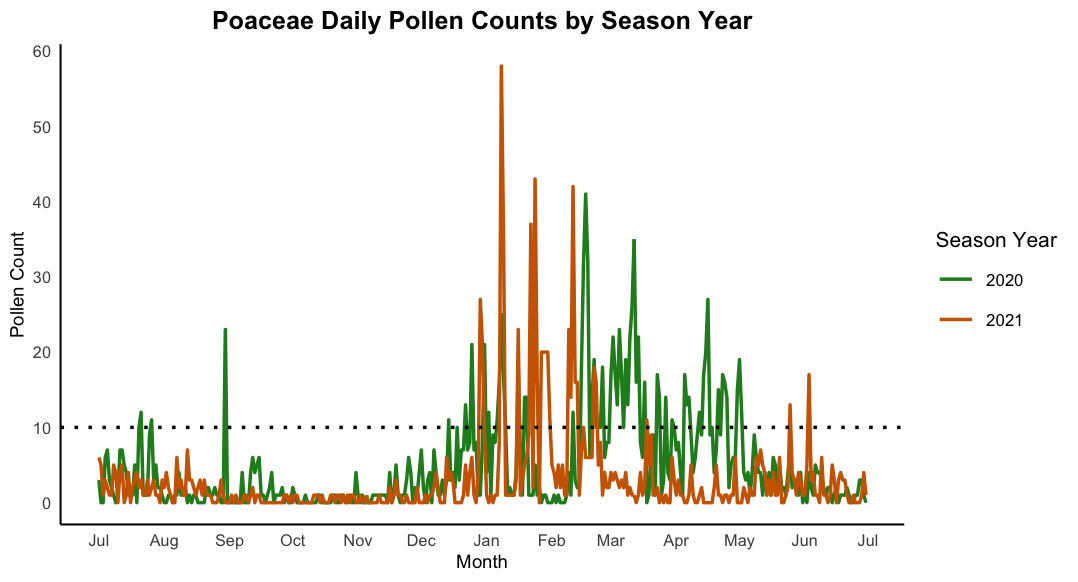

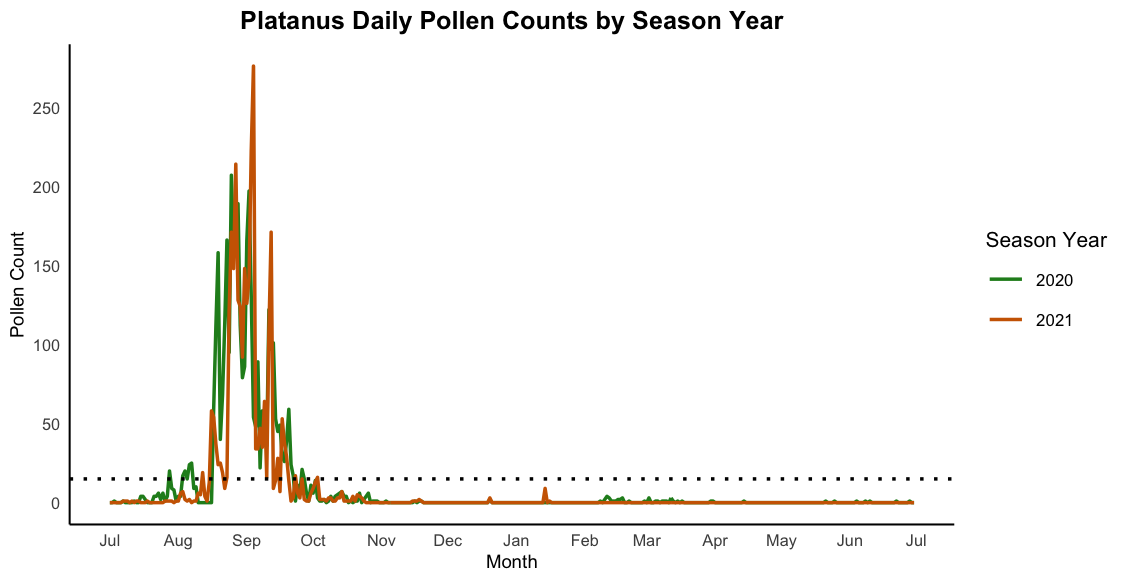

(c) **SAVANNA (PRETORIA)**

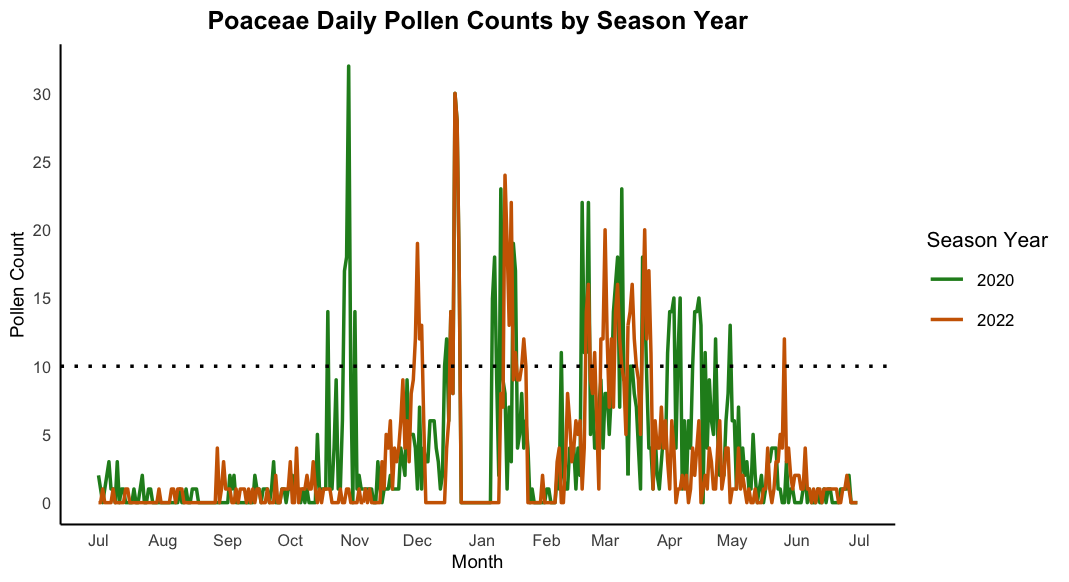

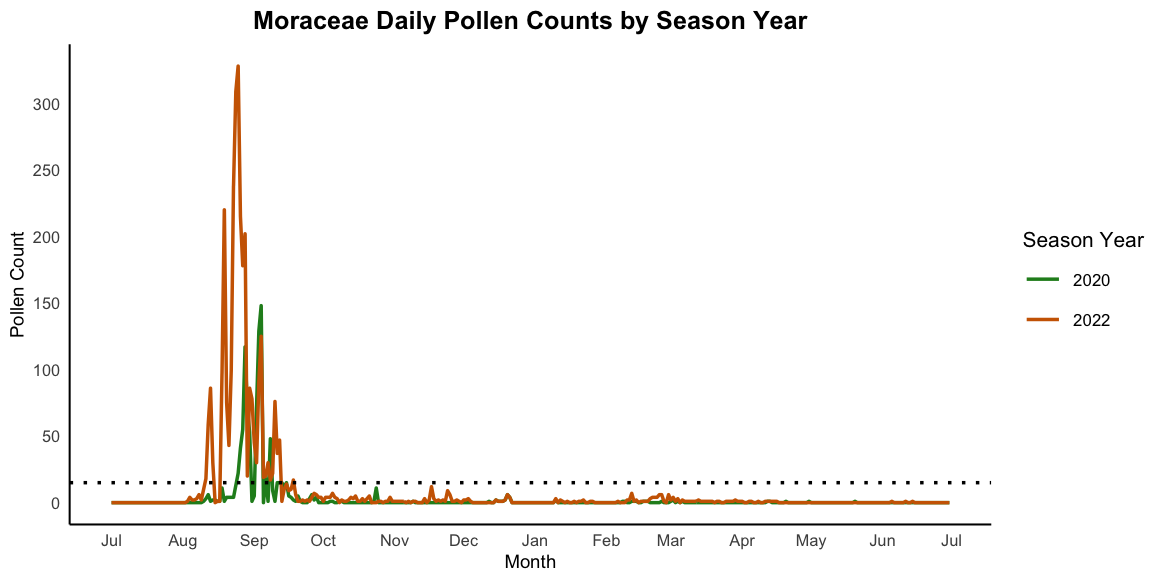

(d) **INDIAN OCEAN COASTAL BELT (DURBAN)**

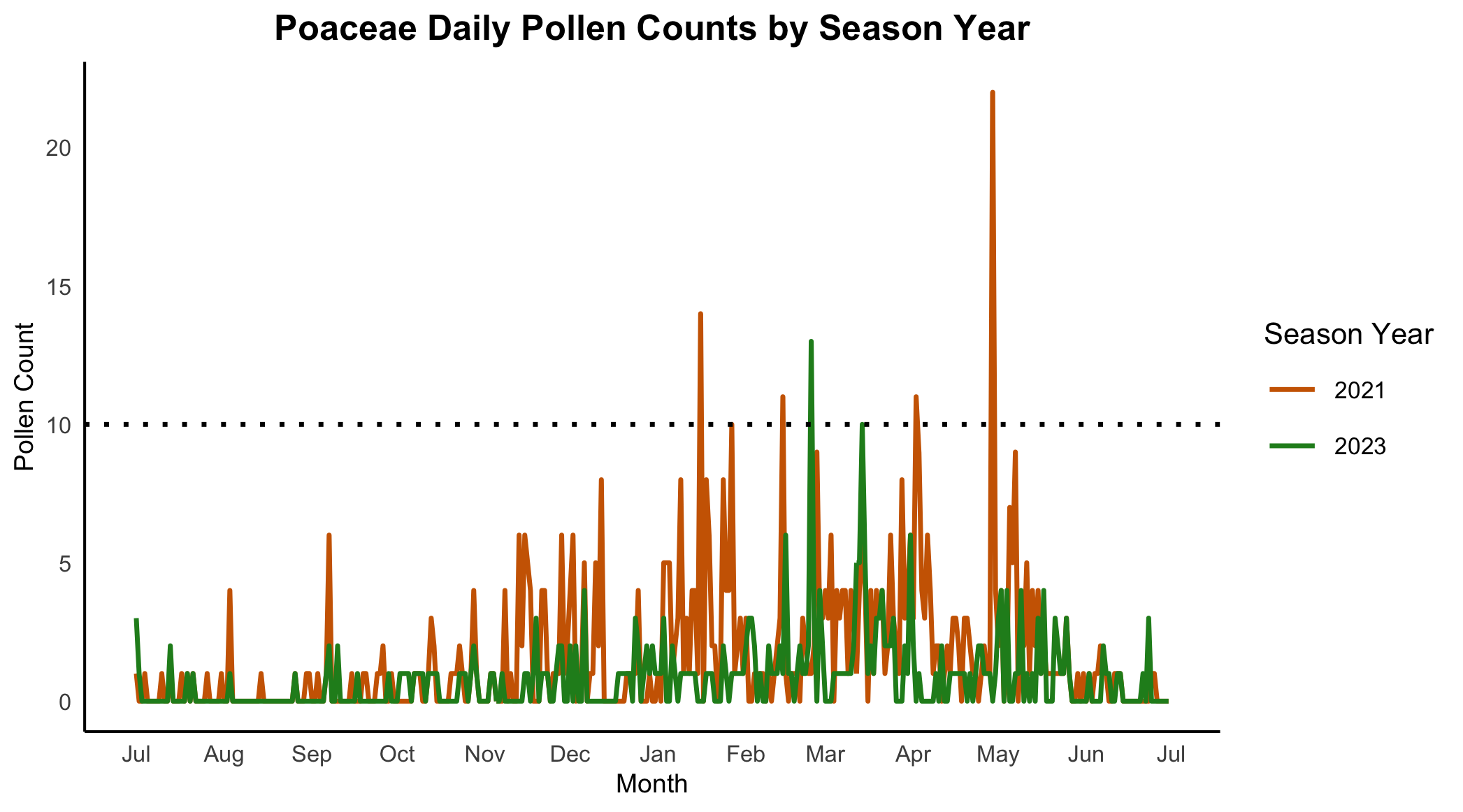

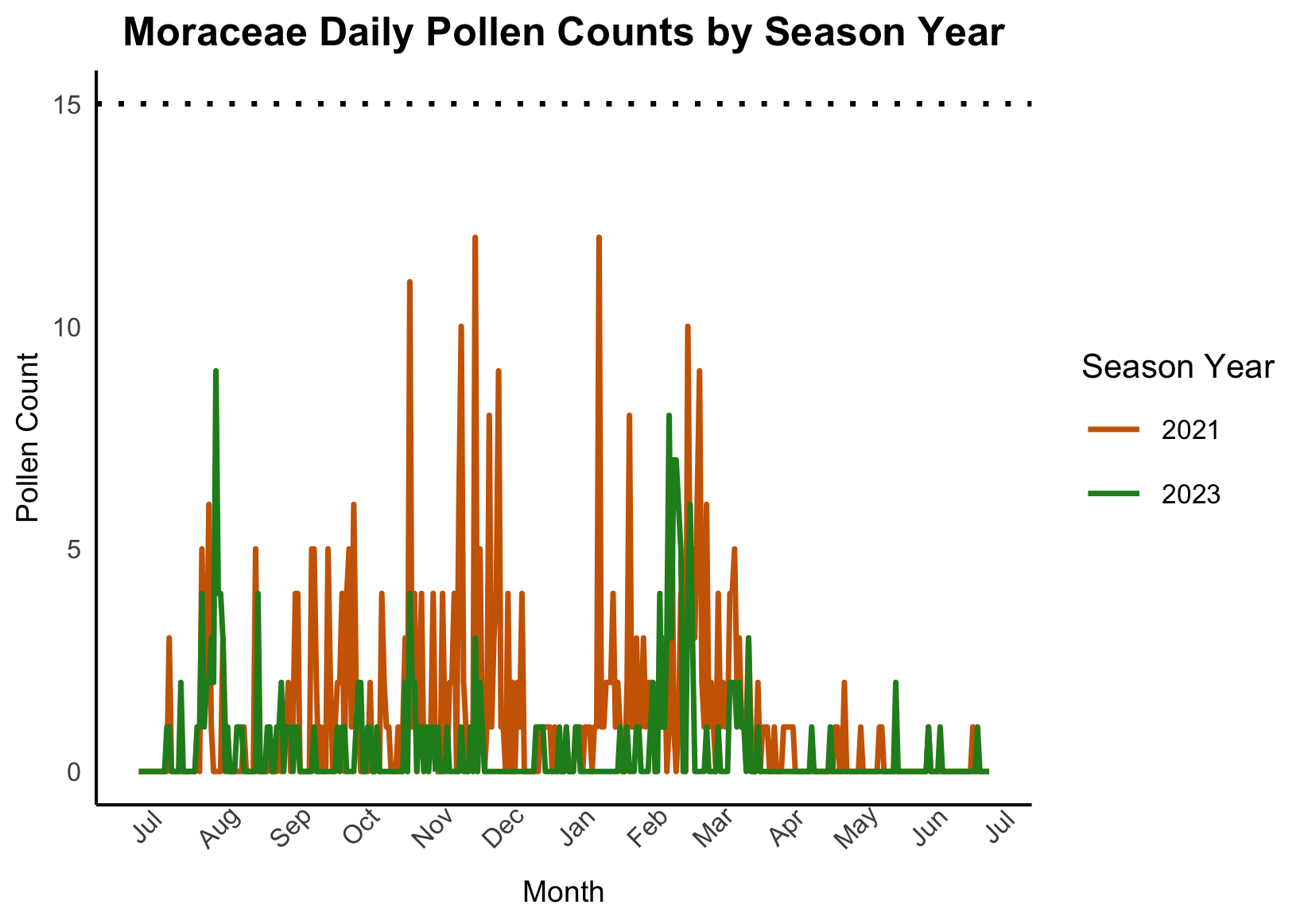

(e) **GRASSLAND (BLOEMFONTEIN)**

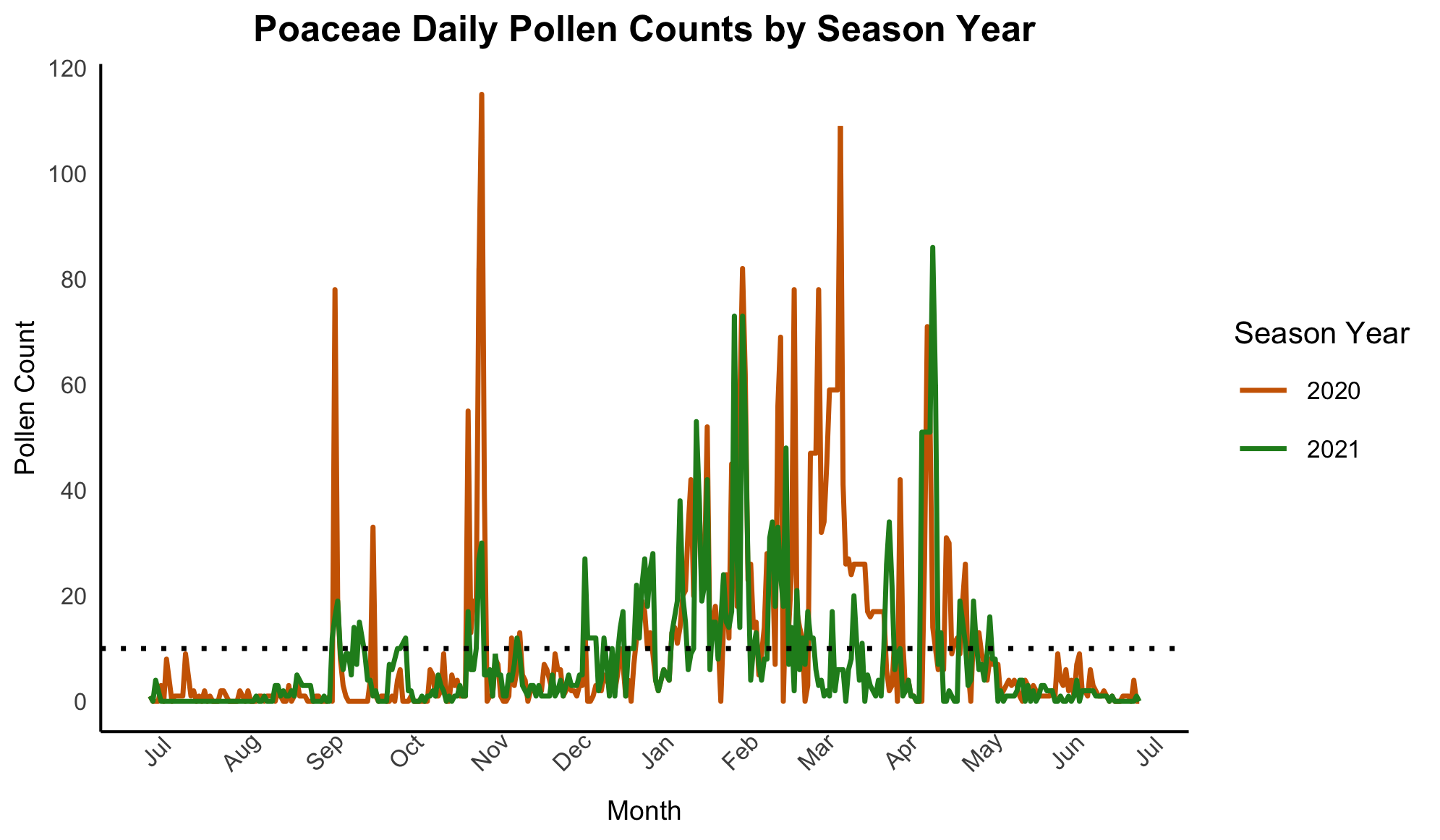

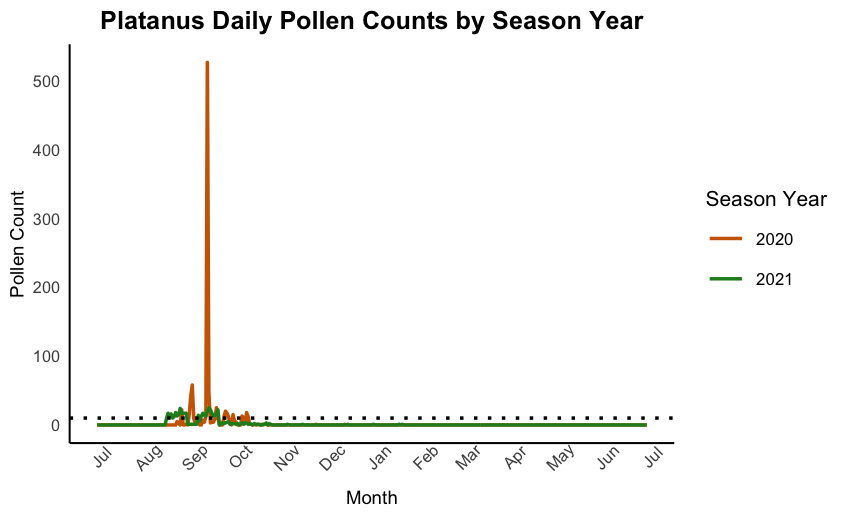

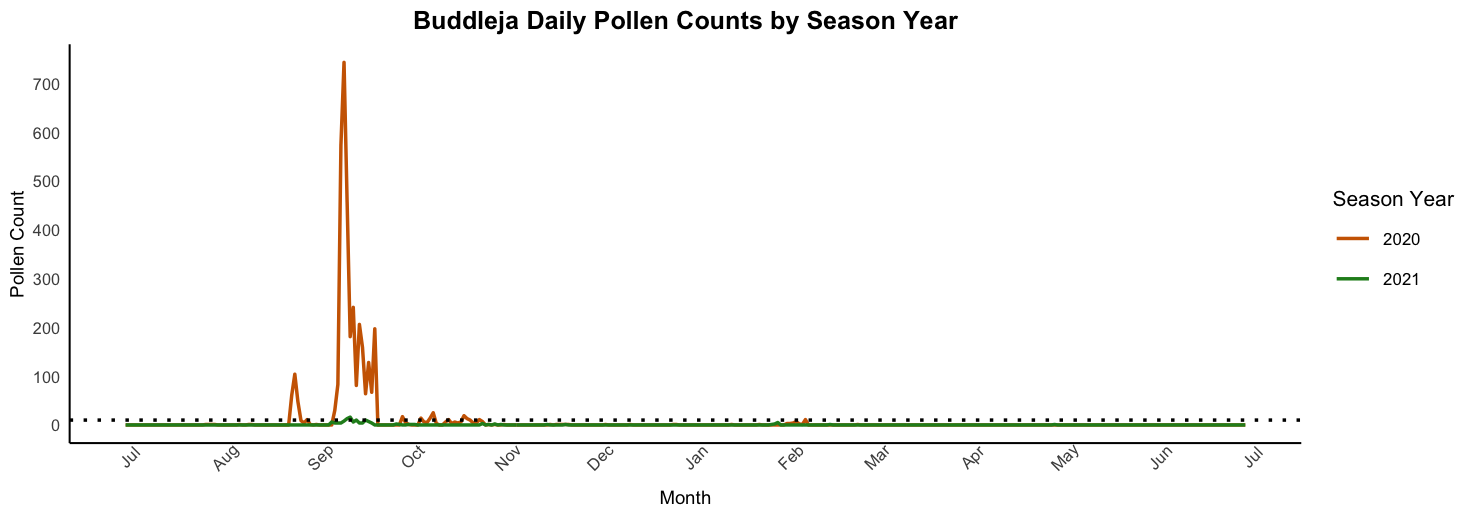

(f) **ALBANY THICKET (GQEBERHA)**

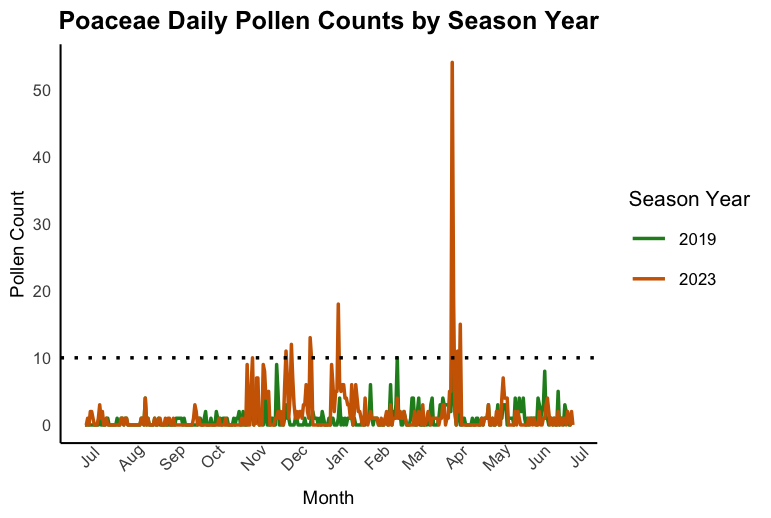

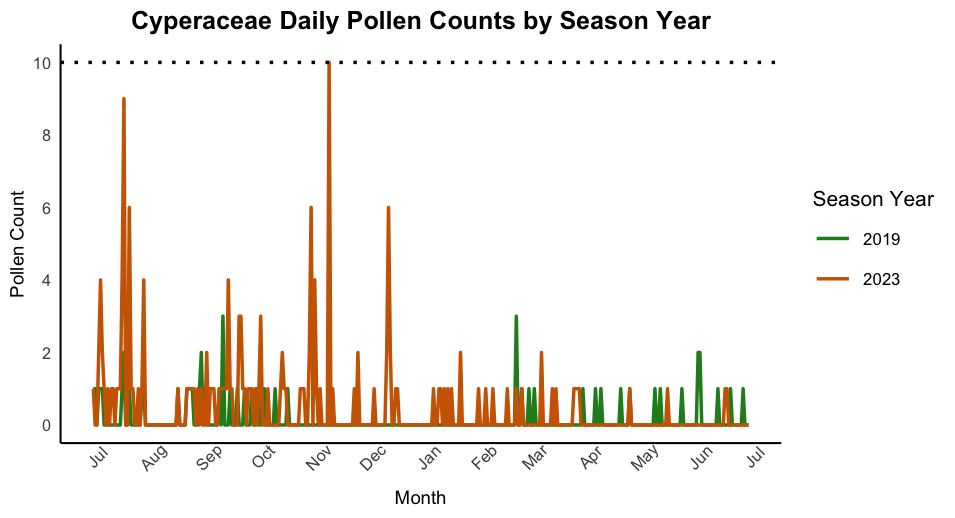

(g) **SAVANNA (KIMBERLEY)**

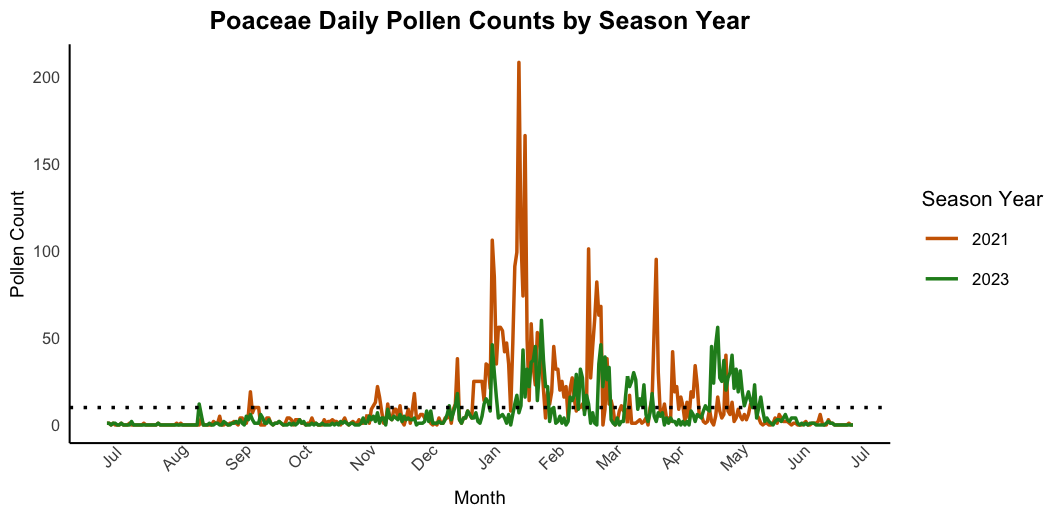

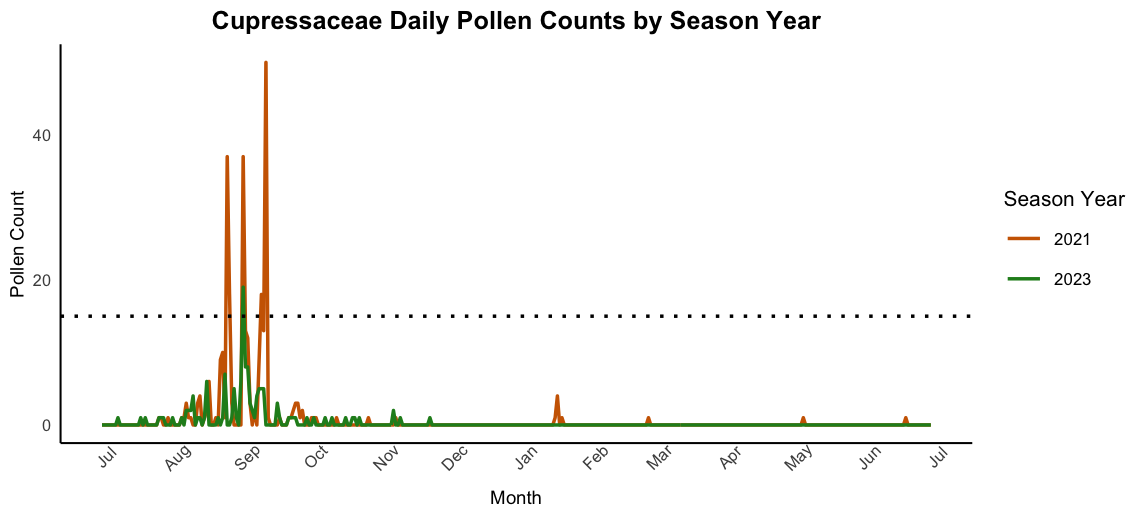

**SUPPLEMENTARY TABLES**

**Table S1** The geography and climate of the biomes in South Africa

| **Biome** | **City** | **Köppen**  **Climate /**  **Humidity**  **Range (%)** | **Rainfall Pattern /**  **Annual Average Rainfall** | **Average  Temperatures**  **(Min-Max)** | **Elevation** | **Population** | **GPS Coordinates** | **Dominant**  **Wind Pattern** |
| --- | --- | --- | --- | --- | --- | --- | --- | --- |
| Fynbos | Cape Town | Mediterranean  60-85% | Winter  515 mm | 11.0 °C-21.5 °C | 0 - 1,590 m | 4.8 million | 33°55′31″S  18°25′26″E | Strong South-easterly |
| Grassland | Johannesburg | Subtropical  Highland  40-75% | Summer  713 mm | 9.5 °C-21.9 °C | 1753 m | 8 million | 26°12′16″S  28°2′44″E | Mostly northern winds, with some western or southern winds |
| Savanna | Pretoria | Subtropical  45-80% | Summer  673 mm | 12.0 °C-25.0 °C | 1400 m | 2.9 million | 25°44′46″S  28°11′17″E | Relatively light winds |
| Grassland | Bloemfontein | Semi-arid  30-65% | Summer  559 mm | 7.0 °C- 24.4 °C | 1395 m | 0.7 million | 29°07′S  26°13′E | South-west winds, with occasional northerly/easterly winds |
| Savanna | Kimberley | Semi-arid  25%-60% | Summer  414 mm | 10.9 °C- 26.1 °C | 1184 m | 0.2 million | 28°44′18″S  24°45′50″E | Variable directions |
| Indian Ocean  Coastal Belt | Durban | Humid-  Subtropical  70-95% | Summer  1019 mm | 16.5 °C-25.2 °C | 8 m | 3.6 million | 29°53′S  31°03′E | North-west winds |
| Albany  Thicket | Gqeberha | Oceanic  65-85% | Year-round  624 mm | 13.5 °C-22.3 °C | 60 m | 1.15 million | 33°57′29″S  25°36′00″E | **South-westerly** in winter, and **north-easterly** in summer |

**Table S2** Summary of pollen taxa that contributed at least 3% of the APIn across the seven biomes in South Africa

|  | **Biome/ City** | **2019** | **2020** | **2021** | **2022** | **2023** |
| --- | --- | --- | --- | --- | --- | --- |
| **COASTAL** | Fynbos (Cape Town) | Cupressaceae, *Pinus*, Poaceae, Myrtaceae, *Platanus*, *Olea* | Poaceae, Cupressaceae, *Pinus*, Myrtaceae, *Platanus*, Restionaceae | Cupressaceae, Myrtaceae  Poaceae, *Pinus*, *Platanus* | Cupressaceae, *Pinus*  Poaceae, Myrtaceae  Urticaceae, *Platanus* | Cupressaceae, Poaceae  Myrtaceae, *Pinus*  Plantaginaceae, *Morus* |
|  | Indian Ocean Coastal Belt (Durban) | Poaceae, *Betula,* Polypodiaceae, *Morus*, Myrtaceae, Asteraceae, *Pinus*, *Ambrosia*, Cyperaceae | Poaceae, *Morus*  Polypodiaceae,  Asteraceae, Myrtaceae, *Betula*, Cupressaceae, *Pinus* | Poaceae, *Morus*  Urticaceae, Myrtaceae, Polypodiaceae, *Pinus*, Cupressaceae, Plantaginaceae, Asteraceae, Cyperaceae, | *Morus*, Poaceae, Urticaceae, Ulmaceae, Myrtaceae, Cupressaceae, Polypodiaceae, Myricaceae, *Pinus* | Poaceae, *Morus*  *Betula*, *Ambrosia*  Polypodiaceae, Myrtaceae  Cupressaceae, *Pinus*, Urticaceae |
|  | The Albany thicket (Gqeberha) | Poaceae, *Pinus*, Cyperaceae, Asteraceae  *Casuarina*, Proteaceae, Myrtaceae | Poaceae, Cyperaceae  *Olea*, Myricaceae, Asteraceae, *Casuarina*, Ericaceae, Caryophyllace, Anacardiacee, *Pinus* | Poaceae, *Casuarina*  Cyperaceae, *Olea,* *Amaranthus*, *Pinus*, *Stoebe*, Anacardiacee, Myricaceae, Asteraceae, Ericaceae | Poaceae, Myricaceae, Thymelaceae, Cyperaceae  *Amaranthus*, Myrtaceae *Pinus*, Asteraceae, *Rhus* / *Searsia*, *Anthospermm* | Poaceae, Typhaceae  Cyperaceae, *Saxifraga*  *Pinus*, Myricaceae, Asteraceae, Restionaceae, Ericaceae |
| **INLAND** | Grassland  (Johannesburg) | *Platanus*, Poaceae, Cupressaceae, *Morus*, *Betula,* *Quercus*, Asteraceae | Platanus, Poaceae, *Morus*, Cupressaceae, *Quercus*, *Betula*, Asteraceae | *Platanus*, Poaceae *Morus*, Cupressaceae, *Betula*, *Fraxinus,* *Quercus* | *Platanus,* Poaceae, Cupressaceae, *Quercus*, *Celtis*, *Morus*, Ulmaceae, Myrtaceae, *Olea* | *Platanus*, Poaceae, Cupressaceae, *Morus*, *Celtis*, *Quercus*, *Fraxinus* |
|  | Savanna  (Pretoria) | Poaceae, *Morus*  *Platanus*, *Betula*, Cupressaceae, Myrtaceae | Poaceae, *Morus,*  Cupressaceae, Stoebe-type | Poaceae, *Morus*  Cyperaceae, *Platanus*, *Betula*, *Prosopis*, Asteraceae, Myrtaceae | *Morus*, *Platanus*  Poaceae, *Betula*, *Fraxinus*, Cupressaceae | *Morus*, Poaceae  *Platanus*, *Betula,* Cupressaceae |
|  | Grassland (Bloemfontein) | Poaceae, Oleaceae  *Buddleja*, Cupressaceae  *Morus*, *Fraxinus* | Poaceae, *Buddleja Fraxinus*, *Platanus*, Cupressaceae, *Pinus*, Asteraceae | Poaceae, *Morus*, *Celtis*  *Platanus*, Myricaceae, *Searsia* (*Rhus*)*,* Asteraceae, *Olea* | *Celtis*, Poaceae, Cupressaceae, *Platanus*, Moraceae, *Populus*, *Olea,* *Buddleja* | *Platanus*, *Morus*, Poaceae, *Populus*, *Olea*, Cyperaceae, *Pinus*, *Betula* |
|  | Savanna Kimberley | Poaceae | Poaceae, *Olea*  Cupressaceae | Poaceae, Urticaceae, Cupressaceae, *Celtis*  *Olea*, Anacardiaceae | Poaceae, *Betula*, Cupressaceae, *Morus*, *Platanus*, Anacardiaceae | Poaceae, Oleaceae  *Celtis*, Cupressaceae, *Morus* |

**Table S3** The average percentage contribution of pollen taxa to the APIn

|  | **CAPE TOWN** | **DURBAN** | **GQEBERHA** | **JOHANNESBURG** | **PRETORIA** | **BLOEMFONTEIN** | **KIMBERLEY** | **All CITIES** |
| --- | --- | --- | --- | --- | --- | --- | --- | --- |
| Classification | Percentage | Percentage | Percentage | Percentage | Percentage | Percentage | Percentage | Average Percentage |
| *Acacia* | 0.04 | 0.36 | 0.99 | 0.14 | 0.24 | 0.56 | 0.08 | 0.34 |
| *Acer* | 0.02 | 0.03 | 0.09 | 0.01 | 0 | 0 | 0.05 | 0.03 |
| Aizoaceae | 0.06 | 0.28 | 0.04 | 0 | 0 | 0.08 | 0.08 | 0.08 |
| *Amaranthus* | 0 | 0 | 2.64 | 0.46 | 0.18 | 0.02 | 0 | 0.47 |
| Amaryllidaceae | 0 | 0 | 0.44 | 0.05 | 0.01 | 0 | 0.01 | 0.07 |
| *Ambrosia* | 0 | 2.79 | 0 | 0 | 0.08 | 0 | 0.18 | 0.43 |
| Anacardiaceae | 1.13 | 1.28 | 1.69 | 0.04 | 0.06 | 0.55 | 1.86 | 0.94 |
| *Anthospermum* | 0 | 0 | 1.40 | 0 | 0.06 | 0 | 0 | 0.21 |
| Apiaceae | 0.05 | 0.06 | 0.01 | 0.01 | 0 | 0 | 0 | 0.02 |
| Araucariaceae | 0.07 | 0.16 | 0.10 | 0 | 0 | 0 | 0.05 | 0.05 |
| Arecaceae | 0.14 | 0.24 | 0.10 | 0.11 | 0.03 | 0.01 | 0 | 0.09 |
| *Artemisia* | 0.25 | 1.05 | 1.27 | 0.16 | 0.46 | 0.38 | 0.25 | 0.55 |
| Asphodelaceae | 0.03 | 0.01 | 0.35 | 0 | 0.01 | 0.02 | 0.14 | 0.08 |
| Asteraceae | 1.25 | 4.02 | 4.26 | 2.41 | 2.03 | 2.37 | 1.74 | 2.58 |
| *Betul*a | 0.33 | 4.85 | 0.21 | 4.13 | 8.05 | 1.44 | 2.7 | 3.11 |
| Boraginaceae | 0.05 | 0.19 | 0 | 0 | 0.01 | 0.01 | 0 | 0.04 |
| Brassicaceae | 0.02 | 0.10 | 0.01 | 0 | 0.01 | 0 | 0.06 | 0.03 |
| *Buddleja* | 0 | 0.01 | 0 | 0.02 | 0.34 | 8.28 | 0.19 | 1.26 |
| *Cannabis* | 0 | 0.01 | 0 | 0.46 | 0 | 0 | 0 | 0.07 |
| *Carya* | 0.01 | 0.68 | 0.23 | 0.03 | 0.05 | 0.02 | 0.58 | 0.23 |
| Caryophyllaceae | 0.15 | 0.19 | 1.76 | 1.45 | 0.46 | 0.02 | 0.27 | 0.62 |
| *Casuarina* | 0.21 | 0.71 | 4.31 | 0.52 | 0.10 | 0.16 | 0.35 | 0.91 |
| *Cedrus* | 0.38 | 0.12 | 0.90 | 0 | 0.01 | 0 | 0.02 | 0.21 |
| *Celtis* | 0.04 | 0.87 | 0.01 | 2.70 | 0.87 | 6.52 | 2.06 | 1.87 |
| Chenopodiaceae | 0.98 | 1.17 | 1.06 | 0.98 | 0.86 | 1.01 | 1.74 | 1.11 |
| *Citrus* | 0 | 0 | 0 | 0 | 0.01 | 0 | 0 | 0 |
| Combretaceae | 0.05 | 0.32 | 0.03 | 0.64 | 1.61 | 0.72 | 1.19 | 0.65 |
| Crassulaceae | 0.01 | 0.01 | 0 | 0 | 0 | 0.32 | 0 | 0.05 |
| Cupressaceae | 30.23 | 4.27 | 0.79 | 7.55 | 3.48 | 4.47 | 3.816 | 7.80 |
| Cyperaceae | 1.37 | 2.91 | 9.26 | 1.62 | 3.39 | 2.74 | 0.493 | 3.11 |
| *Delonix* | 0.04 | 0 | 0 | 0 | 0 | 0.01 | 00 | 0.01 |
| Dodonaea | 0 | 0 | 0.63 | 0.01 | 0.27 | 0.01 | 0 | 0.13 |
| Ebenaceae | 0.11 | 0.03 | 0.03 | 0 | 0 | 0 | 0 | 0.02 |
| Ericaceae | 0.88 | 0.25 | 3.02 | 0.51 | 0.33 | 0.16 | 0.09 | 0.75 |
| *Erodium* | 0.01 | 0.01 | 0.01 | 0 | 0 | 0 | 0 | 0.01 |
| *Erythrina* | 0 | 0 | 0 | 0.02 | 0.12 | 0.01 | 0 | 0.02 |
| Euclea | 0.01 | 0 | 0.03 | 0 | 0.07 | 0.02 | 0.05 | 0.03 |
| Euphorbiaceae | 0.14 | 0.27 | 1.40 | 0.40 | 0.36 | 0.01 | 0.33 | 0.41 |
| Fabaceae | 0.03 | 0.08 | 0.09 | 0.02 | 0.40 | 0.02 | 0.06 | 0.10 |
| *Fagus* | 0.15 | 0.28 | 0 | 0.10 | 0.62 | 0.13 | 0.34 | 0.23 |
| *Fraxinus* | 0.09 | 0.51 | 0 | 2.35 | 1.98 | 5.04 | 0.92 | 1.56 |
| Gentianaceae | 0 | 0 | 0 | 0.01 | 0.01 | 0 | 0 | 0 |
| Geraniaceae | 0 | 0.11 | 0.03 | 0 | 0 | 0 | 0.01 | 0.02 |
| *Helianthus* | 0 | 0 | 0 | 0 | 0.01 | 0 | 0 | 0 |
| Hippocastinaceae | 0.01 | 0.15 | 0.01 | 0 | 0.04 | 0.09 | 0.03 | 0.05 |
| *Ilex* | 0 | 0 | 0 | 0.01 | 0.14 | 0 | 0 | 0.02 |
| Iridaceae | 0.02 | 0.02 | 0.10 | 0 | 0.01 | 0.01 | 0.04 | 0.03 |
| *Jacaranda* | 0.02 | 0 | 0 | 0 | 0.34 | 0.02 | 0 | 0.05 |
| Juglandaceae | 0.03 | 0.07 | 0 | 0.01 | 0 | 0.05 | 0 | 0.02 |
| *Juncus* | 0.12 | 0.12 | 0.01 | 0 | 0.01 | 0.02 | 0.02 | 0.04 |
| Lamiaceae | 0 | 0 | 0 | 0.01 | 0.04 | 0.04 | 0 | 0.01 |
| Liliaceae | 0.01 | 0.04 | 0.93 | 0.03 | 0.01 | 0 | 0.01 | 0.15 |
| *Liquidambar* | 0.01 | 0.02 | 0 | 0.08 | 0 | 0 | 0.06 | 0.02 |
| Loranthaceae | 0 | 0.03 | 0 | 0 | 0.01 | 0 | 0 | 0 |
| Malvaceae | 0.03 | 0.20 | 0.01 | 0.03 | 0.02 | 0.09 | 0.08 | 0.07 |
| *Melia* | 0 | 0 | 0 | 0 | 0 | 0.03 | 0 | 0 |
| Moraceae | 1.45 | 13.17 | 0.19 | 7.82 | 25.29 | 8.19 | 2.9 | 8.43 |
| Myricaceae | 0.78 | 1.88 | 5.48 | 0.01 | 0.05 | 1.17 | 0.22 | 1.27 |
| Myrtaceae | 10.92 | 5.41 | 2.80 | 2.58 | 2.96 | 0.99 | 0.74 | 3.77 |
| *Nuxia* sp | 0.96 | 0.32 | 0 | 0 | 0 | 0 | 0.08 | 0.12 |
| Oenotheracea | 0 | 0.41 | 0.07 | 0 | 0 | 0 | 0 | 0.07 |
| Oleaceae | 2.57 | 1.41 | 3.56 | 1.73 | 1.90 | 5.93 | 3.81 | 2.99 |
| Oxalidaceae | 0 | 0 | 0.23 | 0 | 0 | 0 | 0.09 | 0.05 |
| *Parietaria* | 0.04 | 0.52 | 0 | 0.09 | 0.12 | 0 | 0.19 | 0.14 |
| *Persicaria* | 0.07 | 0.39 | 0 | 0.15 | 0 | 0 | 0.29 | 0.13 |
| Picea | 0.01 | 0 | 0 | 0.01 | 0 | 0 | 0.01 | 0 |
| Pinaceae | 0.02 | 0 | 0 | 0 | 0 | 0 | 0 | 0.02 |
| *Pinus* | 9.46 | 3.38 | 5.04 | 2.08 | 1.83 | 3.25 | 1.09 | 3.73 |
| Plantaginaceae | 1.20 | 2.23 | 0.60 | 0.33 | 0.32 | 1.22 | 0.3 | 0.89 |
| *Platanus* | 4.06 | 0.79 | 0.20 | 31.48 | 12.34 | 9.38 | 1.27 | 8.50 |
| Poaceae | 17.26 | 21.88 | 28.08 | 14.79 | 21.02 | 26.13 | 62.4 | 27.37 |
| *Podocarpus* | 0.21 | 0.91 | 0.47 | 1.44 | 0.36 | 0.39 | 0.21 | 0.57 |
| Polygonaceae | 0.24 | 0.52 | 0.26 | 0.04 | 0.01 | 0.10 | 0.56 | 0.25 |
| Polypodiaceae | 0.19 | 6.64 | 0.10 | 0.51 | 0.33 | 0.07 | 0.18 | 1.15 |
| *Populus* | 0.14 | 0.41 | 0.06 | 0.63 | 0.50 | 3.77 | 0.39 | 0.84 |
| *Prosopis* | 0.02 | 0.14 | 0 | 0.01 | 1.18 | 0.11 | 0.35 | 0.26 |
| Proteaceae | 0.18 | 0.38 | 1.27 | 0.01 | 0.02 | 0.02 | 0.08 | 0.28 |
| *Quercus* | 1.85 | 0.55 | 0 | 5.17 | 1.75 | 0.41 | 0.2 | 1.42 |
| Ranunculaceae | 0.12 | 0.07 | 0.03 | 0 | 0 | 0 | 0.05 | 0.04 |
| Restionaceae | 0.62 | 0.73 | 1.44 | 0 | 0.01 | 0 | 0.05 | 0.41 |
| Rhamnaceae | 0.03 | 0.01 | 0.12 | 0.01 | 0.01 | 0.02 | 0.01 | 0.03 |
| *Rhus* / *Searsia* | 0.85 | 0.40 | 1.17 | 0.73 | 0.51 | 1.60 | 0.91 | 0.84 |
| Rosaceae | 0.05 | 0 | 0 | 0 | 0 | 0 | 0 | 0.01 |
| *Rumex* | 0.76 | 0.59 | 0.04 | 0.02 | 0.08 | 0.05 | 0.53 | 0.30 |
| Rutaceae | 0.09 | 0.03 | 0.04 | 0 | 0 | 0 | 0.01 | 0.02 |
| *Salix* | 0.03 | 0.03 | 0 | 0.27 | 0.32 | 0.03 | 0.1 | 0.11 |
| Sapotaceae | 0 | 0.03 | 0 | 0 | 0.01 | 0 | 0 | 0.01 |
| *Saxifraga* | 0 | 0.03 | 2.01 | 0.01 | 0 | 0 | 0.06 | 0.30 |
| *Schinus* | 0.67 | 0 | 0 | 0 | 0 | 0 | 0 | 0.67 |
| Sclerocarya | 0 | 0 | 0.25 | 0 | 0.03 | 0 | 0 | 0.04 |
| Scrophulariaceae | 0.03 | 0.03 | 0 | 0 | 0 | 0.01 | 0.01 | 0.01 |
| Solanaceae | 0.01 | 0.02 | 0 | 0.03 | 0 | 0 | 0 | 0.01 |
| *Stoebe* | 0.25 | 0.40 | 2.38 | 0.78 | 1.09 | 0.05 | 0.16 | 0.94 |
| *Taraxacum* | 0.03 | 0.01 | 0.12 | 0 | 0.02 | 0.01 | 0.01 | 0.03 |
| *Theylepteris* | 0 | 0.01 | 0 | 0.01 | 0 | 0 | 0 | 0 |
| Thymelaceae | 0.17 | 0.25 | 2.39 | 0.09 | 0.04 | 0 | 0.13 | 0.52 |
| Tiliaceae | 0.10 | 0.09 | 0.26 | 0.13 | 0.02 | 0.03 | 0.02 | 0.09 |
| Typhaceae | 1.38 | 0.22 | 2.83 | 0.22 | 0.18 | 0.02 | 0.18 | 0.72 |
| *Ulmus* | 0.86 | 1.55 | 0.01 | 1.50 | 0.04 | 1.03 | 0.34 | 1.12 |
| Umbelliferae | 0 | 0.05 | 0 | 0.01 | 0.01 | 0 | 0 | 0.01 |
| Urticaceae | 2.43 | 5.08 | 0.12 | 0.05 | 0.23 | 0.51 | 1.84 | 1.46 |
| *Zea mays* | 0.04 | 0.18 | 0.03 | 0.13 | 0.15 | 0.02 | 0.21 | 0.11 |
| Zygophyllaceae | 0.01 | 0 | 0.06 | 0 | 0 | 0.05 | 0.01 | 0.02 |

**Table S4** Summary of the number of pollen taxa that were recorded at each city in South Africa

| **Site** | **Total** | **Grass** | **Trees** | **Weeds** |
| --- | --- | --- | --- | --- |
| Albany Thicket (Gqeberha) | 70 | 2 | 31 | 37 |
| Grassland (Bloemfontein) | 78 | 2 | 41 | 35 |
| Savanna (Pretoria) | 84 | 2 | 44 | 38 |
| Fynbos (Cape Town) | 83 | 2 | 41 | 40 |
| Indian Ocean  Coastal Belt (Durban) | 82 | 2 | 37 | 43 |
| Grassland (Johannesburg) | 82 | 2 | 40 | 40 |
| Savanna (Kimberley) | 78 | 2 | 38 | 38 |
| Total | 105 | 2 | 51 | 52 |
